# A Fast Lysine Cross-linker DOPA Enables Mass Spectrometry Analyses of Protein Unfolding and Weak Protein-protein Interactions

**DOI:** 10.1101/2020.11.05.369280

**Authors:** Jian-Hua Wang, Yu-Liang Tang, Rohit Jain, Fan Xiao, Zhou Gong, Yu Zhou, Dan Tan, Qiang Li, Xu Dong, Shu-Qun Liu, Chun Tang, Niu Huang, Keqiong Ye, Meng-Qiu Dong, Xiaoguang Lei

**Affiliations:** National Institute of Biological Sciences (NIBS), Beijing 102206, China; Tsinghua Institute of Multidisciplinary Biomedical Research, Tsinghua University, Beijing 102206, China; Beijing National Laboratory for Molecular Sciences, Key Laboratory of Bioorganic Chemistry and Molecular Engineering of Ministry of Education, Synthetic and Functional Biomolecules Center, College of Chemistry and Molecular Engineering, Peking-Tsinghua Center for Life Science, Peking University, Beijing 100871, China; University of Massachusetts Medical School, Worcester MA 01605, USA; Wuhan Institute of Physics and Mathematics, Chinese Academy of Sciences (CAS), Wuhan 430071, China; State Key Laboratory for Conservation and Utilization of Bio-Resources in Yunnan, Yunnan University, Kunming 650091, Yunnan, China; Key Laboratory of RNA Biology, CAS Center for Excellence in Biomacromolecules, Institute of Biophysics, Chinese Academy of Sciences, Beijing 100101, China; University of Chinese Academy of Sciences, Beijing 100049, China

## Abstract

Chemical cross-linking of proteins coupled with mass spectrometry analysis (CXMS) has become a widely used method for protein structure analysis. Central to this technology are chemical cross-linkers. The most popular cross-linkers are N-hydroxysuccinimide (NHS) esters, which react with protein amino groups relatively slowly over 10 minutes or more while in competition with the hydrolysis reaction of NHS esters. To improve the speed of cross-linking, we developed a new class of amine-selective and non-hydrolyzable <u>d</u>i-*<u>o</u>rtho*-<u>p</u>hthal<u>a</u>ldehyde (DOPA) cross-linkers. DOPA can cross-link proteins in 10 seconds under near physiological conditions, which is 60 times faster than the NHS ester cross-linker DSS. DOPA also works at low pH, low temperature, or in the presence of high concentrations of denaturants such as 8 M urea or 6 M guanidine hydrochloride. Further, DOPA-mediated pulse cross-linking captured the dynamic conformational changes associated with RNase A unfolding. Lastly, DOPA outperformed DSS at capturing weak but specific protein-protein interactions.

## Introduction

Proteins constantly undergo conformational changes. Protein motions span a wide range of temporal and spatial scales, from the fast fluctuations of the amino acid side chains in picoseconds to the moderate fluctuations of the surface loops in nanoseconds and to the slow collective motions of the domain/entire protein in microseconds to seconds^1, 2^. The biological function of a protein is rooted in its physical motions and dynamic properties^1, 3^. Subunits of protein complexes may initially form encounter complexes (EC) before assembling into the final complex^4^. Proteins experience the most dramatic conformational changes during folding and unfolding and they occur at the large time-scales of subseconds to minutes^5^. Characterizing the different conformational states (i.e., folded, partially folded, and unfolded states) of proteins and probing the conformational transitions between these states are of fundamental importance to biology and of practical value to drug development.

A variety of tools have been developed for conformational studies of proteins. X-ray crystallography and single particle cryo-electron microscopy (cryo-EM) have revolutionized structural biology by taking “snapshots” of proteins or protein complexes at atomic resolution^6, 7^. However, these technologies are less effective for protein dynamics studies. NMR spectroscopy can provide valuable information on the dynamics of a protein, but NMR is typically limited to proteins of < 50 kDa^8, 9^. Fluorescence resonance energy transfer^10, 11^ and electron paramagnetic resonance^12^ circumvent protein size limitations by targeted insertion of donor and acceptor fluorophores, or two spin labels, respectively. However, this requires a prior hypothesis and elaborate protein manipulation. Small-angel X-ray scattering (SAXS) is increasingly used to characterize the protein dynamics in solution and without fluorophores ^13, 14^. However, it can only yield changes in radius of gyration, dimensions and macromolecular shape^15^.

Mass spectrometry-based methods, such as hydrogen-deuterium exchange^16^, hydroxyl radical footprinting^17^, can be used to probe the dynamics of a protein based on the difference in the solvent accessibility among various parts of the protein structure, but they do not provide the three-dimensional (3D) structure information directly. Chemical cross-linking of proteins coupled with mass spectrometry analysis (CXMS, XL-MS, or CLMS) is a straightforward approach for investigating protein structures and protein-protein interactions^18–23^. CXMS has a great potential to capture the conformational changes in proteins, owing to the rapid linkage of two amino acid residues lying close to each other in the tertiary structures of proteins. Encouragingly, CXMS has already been utilized successfully to visualize the co-existing protein conformational states^24^ and to detect the conformational change of a protein under different conditions^25^. However, to our knowledge, CXMS has not been used to study continuous protein conformational changes, because cross-linking reactions are usually too slow to report them.

Although a number of chemical cross-linkers can target Lys^26–29^, Arg^30^, Cys^31^, or acidic amino acids^32, 33^, CXMS has largely focused on the lysine-targeting N-hydroxysuccinimide (NHS) ester cross-linkers such as disuccinimidylsuberate (DSS), bis(sulfosuccinimidyl)suberate (BS^3^), and disuccinimidyl sulfoxide (DSSO) owing to their high reactivity and an abundance of lysine residues on protein surface^20^. However, the NHS ester-based cross-linking reaction is slow and typically takes 30-60 minutes on protein substrates^34^. Besides, NHS ester is susceptible to rapid hydrolysis in aqueous solutions. The half-life of an NHS ester is about tens of minutes under the typical reaction conditions^35^. It is thus necessary to develop a fast and stable cross-linker.

The earliest report of *<u>o</u>rtho*-<u>p</u>hthal<u>a</u>ldehyde (OPA) reacting with an amino group to form an isoindolinone skeleton can be traced back to 1909^36^. Since then, many OPA-related applications have been reported, including determination of serum protein concentration^37^, polymer synthesis^38^, and disinfection of surgical equipments^39^. A recent study reported that OPA can selectively modify amino groups on natural protein surfaces^40^. Attracted by the unique properties of OPA—amine-selectivity, fast reaction, and no hydrolysis, we set out to develop a new class of amine-selective non-hydrolyzable <u>d</u>i-*<u>o</u>rtho*-<u>p</u>hthal<u>a</u>ldehyde (DOPA) cross-linkers. In this work, we show that DOPA2, with a spacer arm of two ethylene glycol units, cross-linked proteins ~60 times faster than DSS. Furthermore, DOPA2 can cross-link proteins under certain extreme conditions, e.g., low temperature, low pH, or in the presence of high concentrations of denaturants such as 8 M urea or 6 M guanidine hydrochloride (GdnHCl). These unique properties of DOPA2 make it an ideal cross-linker for probing the protein folding/unfolding intermediate states and for capturing the sequential changes during protein unfolding, as demonstrated with the model protein bovine pancreatic ribonuclease A (RNase A). Lastly, we demonstrate that DOPA2 outperforms the NHS ester cross-linkers in capturing weak but specific protein-protein interactions.

## Results

### I. Development of a new class of amine-reactive cross-linkers

First, we verified that OPA reacts with lysine faster than an NHS ester (Supplementary Figure 1), as reported before^40^. Next, we tested the OPA reaction with 10 synthetic peptides covering all 20 common amino acids (Supplementary Table 1). The liquid chromatography coupled with tandem mass spectrometry (LC-MS/MS) analysis confirmed that the major products were N-substituted phthalimidines^40^ formed on lysine ε-NH_2_ or N-terminal α-NH_2_ (Δ mass = +116.0262 Da), as expected (product 1 in Supplementary Table 2 and Supplementary Figure 2). The loop-linked side product (product 2, Δ mass = +98.0156 Da) resulted from the conjugation of OPA with an amino group and with another nucleophilic group on the same peptide; such nucleophilic groups included α-NH_2_, ε-NH_2_, a sulfhydryl group from cysteine, and a phenolic hydroxyl group from tyrosine (Supplementary Table 2 and Supplementary Figure 2). Apart from corroborating previous reports^40, 41^, our results established a foundation for further development of OPA-based cross-linkers.

Next, we synthesized three <u>d</u>i-*<u>o</u>rtho*-<u>p</u>hthal<u>a</u>ldehyde (DOPA) cross-linkers (Scheme 1). In DOPA-C_2_, the two OPA moieties are connected via one ethylene group. In DOPA1 and DOPA2, the spacer arm consists of one and two ethylene glycol units, respectively (Scheme 1). Both DOPA1 and DOPA2 have better water solubility than DOPA-C_2_.

Using bovine serum albumin (BSA) as a model protein, we optimized the cross-linking conditions for the DOPA cross-linkers. The highest number of cross-linked peptide pairs was obtained after a 10-minute reaction with BSA at a protein:cross-linker ratio of 16:1 (BSA:DOPA, w/w), 25 °C, pH 7.4, and the three amine-free buffer systems—HEPES, PBS, and trimethylamine—worked similarly well (Supplementary Figure 3a-c). Cross-linking at a protein:DOPA2 ratio of 4:1 (w/w) also performed well, provided that two parallel protease digestions—trypsin plus Asp-N or trypsin alone—were carried out. It bears emphasis that these experiments identified a critical factor for successful DOPA cross-linking: pre-dilution of DOPA to two times its final concentration (Supplementary Figure 3d).

### II. Performance of DOPA vs DSS on model proteins

Using a ten-protein mixture as a test sample, we found that peptide pairs cross-linked by DOPA2 (161) outnumbered those by DOPA1 (117) or DOPA-C_2_ (57), and two thirds or more of the DOPA1 or DOPA-C_2_ cross-linked lysine pairs were also found among the DOPA2 dataset (Figure 1a-c and Supplementary Table 3). Next, we conducted a side-by-side comparison of DOPA2 and the widely used NHS ester cross-linker DSS on a panel of six model proteins. For all six proteins with the exception of catalase, DSS cross-links outnumbered DOPA2 cross-links (Figure 1d). However, the Euclidean distance analysis of each cross-linked residue pair revealed that 44.8% of the DSS cross-links exceeded the maximum allowed cross-linking distance (24.0 Å for DSS, Cα-Cα), indicative of a structural compatibility rate of 55.2% for DSS cross-links, which is much lower than that of DOPA2 cross-links (88.3% within the maximum allowed cross-linking distance of 30.2 Å) (Figure 1e). The same conclusion can be made using the Solvent Accessible Surface Distance (SASD) (Figure 1e). DOPA-C_2_ also achieved a higher structural compatibility rate than DSS (79.0% vs 55.2% by Euclidean distance, or 52.9% vs 34.3% by SASD), although they have similar maximum allowed cross-linking distances (24.9 Å and 24.0 Å, respectively) (Figure 1e).

**Figure 1.**
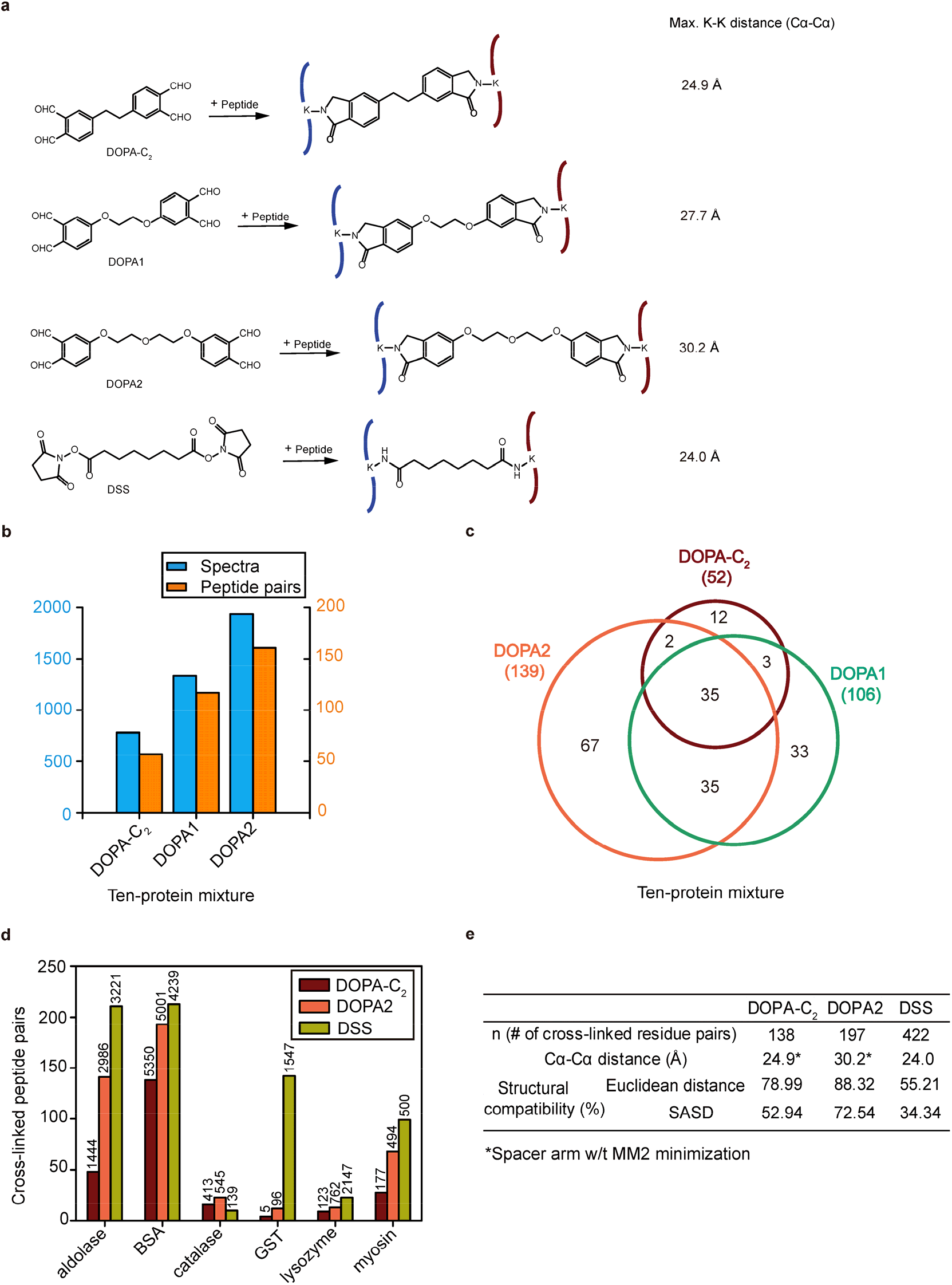
Evaluating the performance of DOPA. (a) Cross-linking products of DOPA-C_2_, DOPA1, DOPA2, and DSS with peptides. (b) Cross-links identified from a ten-protein mixture using DOPA-C_2_, DOPA1, and DOPA2. The numbers of cross-linked spectra are plotted with blue columns, and the number of cross-linked peptide pairs are plotted with orange columns. (c) Venn diagram showing the overlap of residue pairs produced by DOPA-C_2_, DOPA1, and DOPA2 from the ten-protein mixture. Identified cross-links were filtered by requiring an FDR < 0.01 at the spectra level. (d) Performance of DOPA-C_2_, DOPA2, and DSS on model proteins. Numbers of cross-linked peptide pairs are indicated with the colored columns, and spectra identified from each sample are shown above the columns. Identified cross-links were filtered by requiring an FDR < 0.01 at the spectra level. (e) The table displays the percentage of residue pairs that are consistent with the structures of the model proteins, calculated by the use of the Euclidean distance or the solvent accessible surface distance; the table also gives the maximum distance restraints and the number of cross-links belonging to each linker. Identified cross-linking residue pairs were filtered by requiring FDR < 0.01 at the spectra level and with spectral counts ≥ 3.

### III. Advantages of DOPA cross-linking

Although DOPA2 performed no better than DSS in terms of the number of cross-links identified in conventional cross-linking experiments (Figure 1d), it performed better under special conditions, e.g., low temperature, low pH, and high denaturant concentration. Furthermore, DOPA2 cross-linking reactions are about 60-fold faster than DSS on protein substrates, as described below.

As shown in Figure 2a, after just ten seconds of DOPA2 cross-linking at room temperature, the vast majority of BSA molecules appeared as covalent dimers on SDS-PAGE. LC-MS/MS analysis showed that as the cross-linking time increases, the number of identified cross-links increased gradually until reaching a plateau of around 150 pairs after ten minutes. Interestingly, 88 (> 50%) cross-links were identified after only 10 seconds of DOPA2 treatment (Supplementary Figure 4). In contrast, 10 seconds of DSS treatment yielded a tiny amount of covalent dimer, and it took 20 minutes to obtain a similar degree of cross-linking to what DOPA2 achieved in 10 seconds (Figure 2a). We thus estimate in a conservative manner that DOPA2 cross-linking is 60 times faster than DSS on protein substrates. Also worth noting is that even at 0 °C, DOPA2 readily cross-linked BSA within 30 seconds, whereas DSS hardly generated any BSA dimer bands at 0 °C after 30 minutes (Figure 2b). Upon decreasing the pH from 7.4 to 6.0, DOPA2 rather than DSS successfully cross-linked BSA in 10 seconds (Figure 2c). A further decrease in the reaction pH to 3.0 abolished cross-linking of BSA by either DSS or DOPA2. This is surprising because additional experiments revealed that the performance of DOPA2 for reacting with peptide substrates was highly similar at pH 3.0 and pH 7.4 (Figure 2d mono and inter-linked products and Supplementary Table 4a-b). Therefore, the disappearance of DOPA2 cross-linked BSA dimer at pH 3.0 is likely the consequence of acid-induced dissociation of proteins. Additionally, we found that DOPA2 but not DSS was able to cross-link proteins in the presence of urea or GdnHCl within 10 seconds (Figure 2e-g and Supplementary Table 4c). Successful DSS cross-linking was observed in the presence of urea or GdnHCl, but it took a longer time, such as 15 minutes in 1-4 M urea (not shown) or 6 minutes in 1 M GdnHCl (Figure 2f). Again, under high concentrations of denaturants (> 4 M urea or > 1 M GdnHCl), the disappearance of cross-linked BSA dimer bands was largely attributable to dissociation of BSA dimers. The above results clearly indicate that DOPA2 has the capability to cross-link proteins in the denaturing conditions.

**Figure 2.**
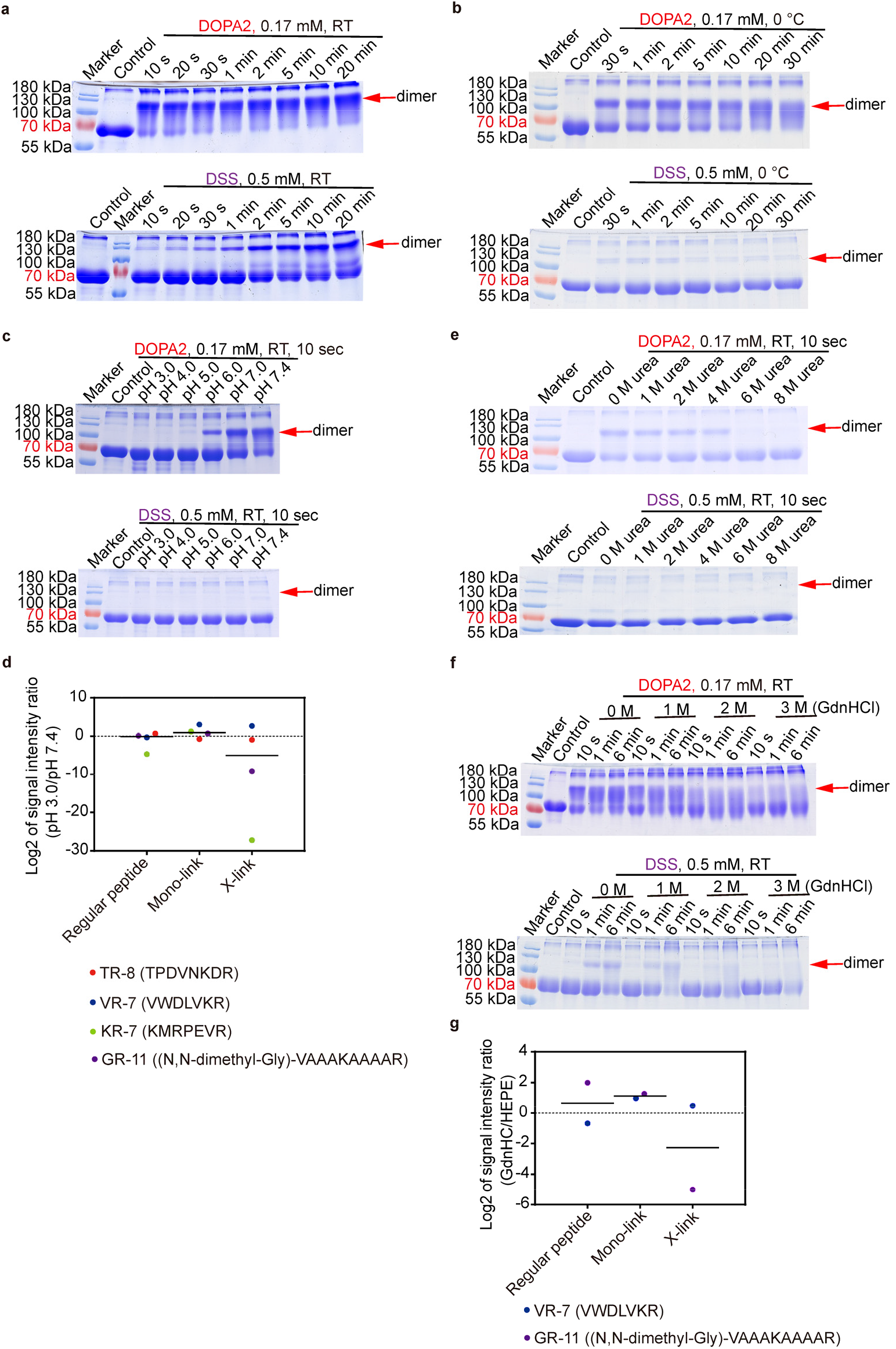
Unique properties of DOPA2 compared with DSS. (a) SDS-PAGE analysis of DOPA2 or DSS-cross-linked BSA under different reaction times at room temperature. (b) SDS-PAGE of DOPA2 or DSS-cross-linked BSA under different reaction times at 0 °C. (c) SDS-PAGE of DOPA2 or DSS-cross-linked BSA under different pH conditions. (d) The log2 values of the signal intensity ratio for different peptide products in a low pH value buffer (3.0) vs. a physiological buffer (7.4). “Regular peptide” refers to free peptides without cross-linking; “X-link” refers to the situation wherein two peptides are linked with one molecule of DOPA2; “Mono-link” refers to a peptide that has been modified but is not cross-linked by a cross-linker. (e and f) SDS-PAGE of DOPA2 or DSS-cross-linked BSA in the presence of the indicated denaturants (urea or GdnHCl). (g) The log2 values of the signal intensity ratio for different peptide products in GdnHCl (6 M) vs. a physiological buffer (HEPES, pH 7.4). The label of products was the same as in d.

### IV. Probing the unfolded states of RNase A by DOPA cross-linking

Because DOPA2 is capable of cross-linking proteins under denaturing conditions, we used it as a probe to investigate the partially or fully unfolded states of bovine pancreatic ribonuclease A (RNase A), a classic model for the protein unfolding/refolding studies^42–45^. In the first set of experiments, equal amounts of RNase A were incubated at room temperature for one hour in 0, 1, 2, 4, 6, or 8 M urea, followed by DOPA2 treatment for ten seconds, protease digestion, and LC-MS/MS analysis (Figure 3a). We made sure that only the most reliable cross-links were used in subsequent analysis by applying a stringent cut-off (FDR<0.01, at least four MS2 spectra at E-value < 1×10^−8^) to the pLink 2^46^ search results.

**Figure 3.**
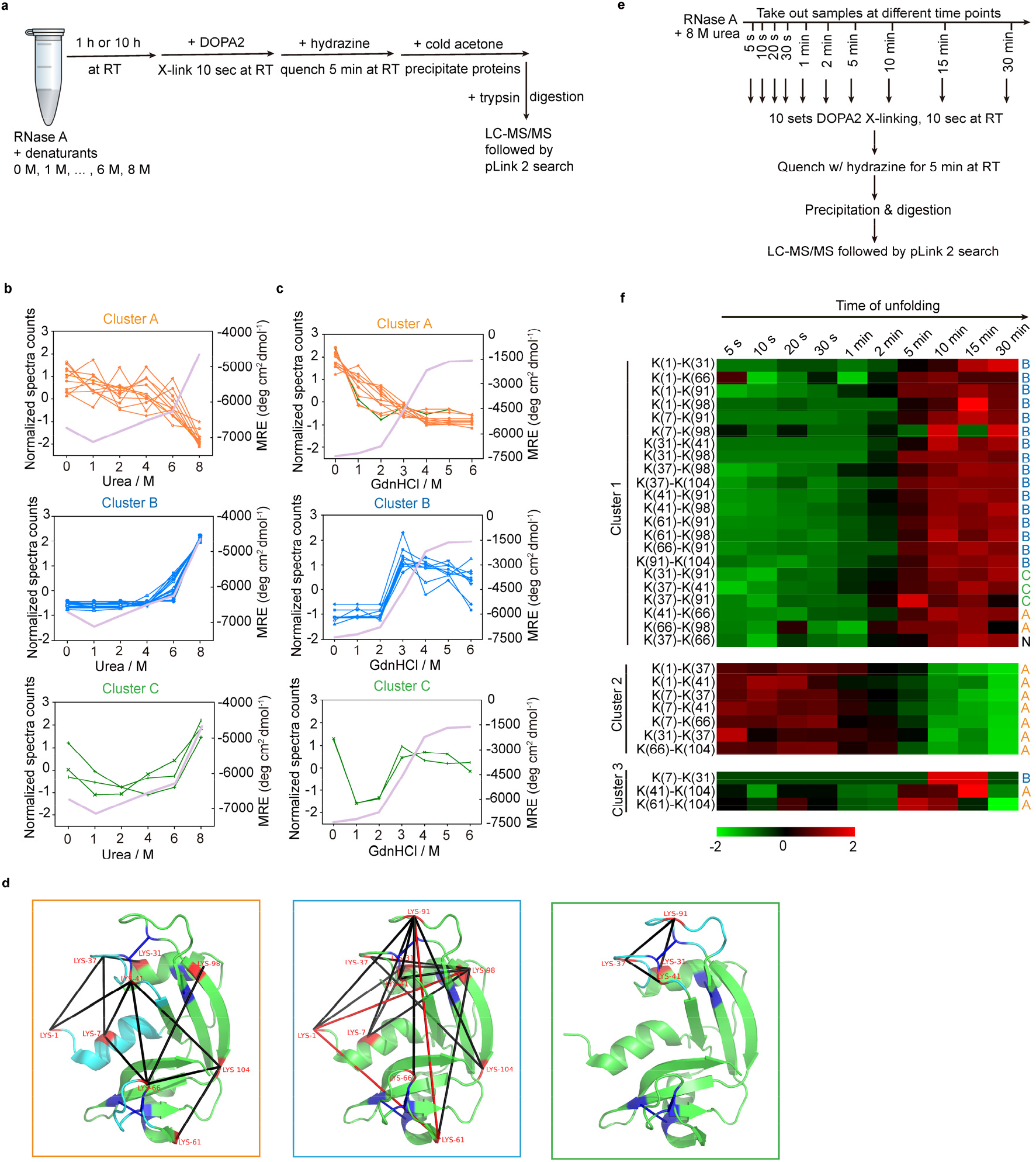
Studying the unfolding of RNase A in chemical denaturants. (a) Experimental workflow of cross-linking RNase A in different concentrations of denaturants (urea or GdnHCl) using DOPA2. (b) The changes of spectral counts for each identified cross-linked residue pair (normalized across the six conditions, shown as Z-scores on the left axis) and the mean residue ellipticity (MRE) of RNase A monitored by CD at 222 nm (on the right axis) in different concentrations of urea. The residue pairs were classified into three clusters by K-means (Cluster A, Cluster B and Cluster C). (c) As in b, but in different concentrations of GdnHCl. (d) The cross-links identified in different concentrations of urea were mapped on the crystal structure of RNase A (PDB code: 6ETK). Cluster A in orange frame (11 pairs); Cluster B in blue frame (17 pairs); and Cluster C in green frame (3 pairs). The black lines denote that the distance of cross-linked residue pairs is within the restraints of cross-linkers, while the red lines denote that the distance of cross-linked residue pairs is out of the restraints of cross-linkers. The four pairs of disulfide bonds are colored by dark blue. (e) Experimental workflow of cross-linking RNase A at different time points in 8 M urea using DOPA2. (f) The changes of spectral counts for each identified cross-linked residue pair in 8 M urea for different time. The residue pairs were classified into three clusters by K-means (Cluster 1, Cluster 2 and Cluster 3). “N” means that the cross-link is not identified in the different concentrations of urea buffers. Cross-linking residue pairs were filtered by requiring FDR < 0.01 at the spectra level, E-value < 1×10^−8^ and spectral counts > 3.

Based on how their abundances changed from 0 M to 8 M urea, the cross-linked residue pairs were clustered into three groups, as shown in Figure 3b and Supplementary Table 5a. The numbers of matched MS2 spectra for each residue pair were first normalized within a sample and then normalized across the six conditions (shown as Z-scores on the left axis). Cluster A represents the high-in-the-native-conformation cluster (native cluster), consisting of DOPA2-linked residue pairs that slowly decrease in abundance from 0 M to 6 M urea, followed by a large drop between 6 M and 8 M urea. Cluster B, the high-in-non-native-conformation cluster (non-native cluster), behaves opposite to the native cluster: the relative abundance of these DOPA2-linked lysine pairs remained low in urea concentrations below 6 M, followed by a small increase in 6 M urea and a large increase in 8 M urea. Cluster C, which has only three members, displays a U-shaped curve as the urea concentration increases.

Consistent with the observations above, circular dichroism (CD) analysis of non-cross-linked RNase A in different concentrations of urea detected little change in mean residue ellipticity (MRE, 222 nm) from 0 M to 6 M urea, then an increase at 7 M and 8 M urea (Supplementary Figure 5a), indicating a substantial loss of secondary structures between 6 M and 8 M urea. Nevertheless, RNase A in 8 M urea did not unfold completely, for its MRE values (−4749 deg cm^2^ dmol^−1^) are far from the MRE value of unfolded RNase A (−1793 deg cm^2^ dmol^−1^) in > 4 M GdnHCl (Supplementary Figure 5).

GdnHCl is a stronger denaturant than urea and the GdnHCl concentration at 50% denaturation of RNase A is ~3 M based on the MRE values at 222 nm (−4683 deg cm^2^ dmol^−1^) (Supplementary Figure 5b). Therefore, we tracked RNase A unfolding by performing CXMS unfolding experiment in GdnHCl (Figure 3a, Figure 3c and Supplementary Table 5b). In agreement with the CD result, as the concentration of GdnHCl increased, the decrease in abundance of the native cluster and the increase in abundance of the non-native cluster quickly reached the maximum level at 3 M GdnHCl (Figure 3c). Strikingly, most cross-links are the same for unfolding in urea and GdnHCl; only one cross-link belonging to the U-shaped cluster from the urea dataset (Figure 3b) had a change of profile in the GdnHCl dataset by joining the native cluster (Figure 3b-c). The above results suggest that protein folding-unfolding induced by urea or GdnHCl can now be investigated by CXMS using DOPA2 despite the different properties of these two denaturants.

A closer examination of the native cluster finds that these DOPA2 cross-links involve mainly lysine residues located on either Helix I (amino acid residues 3-13, colored by cyan) or two surface loops (amino acid residues 34-43, 64-72, colored by cyan) of RNase A (Figure 3d, in orange box), suggesting that these regions are most sensitive to urea. The CD data indicate that at ≤ 4 M urea, Helix I itself is largely intact while the cross-links decrease between Helix I and either of the two loop regions (amino acid residues 34-43, 64-72) (Figure 3b and Supplementary Figure 5a). So, we think that the increased separation between Helix I and the two surface loops in urea is likely due to deformation or displacement of three loop regions—the two above plus the one (amino acid residues 14-33) connecting Helix I to the rest of the protein.

Additionally, we find that the four pairs of disulfide bonds of RNase A (colored dark blue in Figure 3d) are critically involved in defining the path of unfolding. Once these disulfide bonds were disrupted by a reducing reagent, with or without subsequent alkylation, the cross-links displayed completely different abundance profiles from the ones of non-reduced RNase A (compare Supplementary Figure 6 to Figure 3b).

In the non-native cluster, three lysine pairs Lys1-Lys98, Lys61-Lys91, and Lys1-Lys61 (Figure 3d, in blue box) caught our attention, because the two residues in each pair are far apart in primary sequence and in 3D space in the native state. Indeed, these over-length cross-links were hardly detectable from 0 M to 4 M urea and then they spiked in 8 M urea. Therefore, it is unlikely that the non-native conformation is exclusively a stretched linear one; certain globular conformations likely exist.

### V. Analyzing the kinetic RNase A unfolding by DOPA2 cross-linking

Having established that DOPA2 cross-linking can probe the native, partially or fully unfolded states of a protein, we then advanced to studying the kinetics of protein unfolding, i.e., the time-dependence of the transition from the native state to the non-native state. As shown in Figure 3e, RNase A was exposed to 8 M urea for varying amounts of time from 5 seconds to 30 minutes, followed by 10 seconds of DOPA2 cross-linking. The samples were then quenched and processed for LC-MS/MS analysis. After identifying cross-linked peptides through pLink 2 search, we performed cluster analysis to classify the cross-links into groups based on their abundance profile in the unfolding process (Figure 3f and Supplementary Table 5c). Cluster 1 consisted of 22 DOPA2-linked residue pairs whose relative abundance at 8 M urea was low in the two minutes or so and high afterwards. Cluster 2 consisted of seven members and showed an opposite profile with a sharp decrease in abundance after about two minutes of urea exposure (Figure 3f). Cluster 3 was small and less informative. Of note, Cluster 1 is highly enriched (16/22) for members of the non-native cluster (Cluster B in Figure 3b), and Cluster 2 is made up entirely of members of the native cluster (Cluster A in Figure 3b). This shows that a large number of cross-links behave similarly in equilibrium and kinetic unfolding of RNase A. Overall, DOPA2 cross-linking is able to capture the crucial states reflecting the concerted conformational changes during RNase A unfolding.

### VI. Capturing weak but specific protein-protein interactions by DOPA cross-linking

Encouraged by the successful application of rapid DOPA2 cross-linking to the protein unfolding analysis, we set out to find out whether DOPA2 can facilitate analysis of weak protein-protein interactions. We tested this idea by using the bacterial glucose phosphotransferase system^47, 48^, in which the phosphate group from phosphoenolpyruvate is transferred to glucose via Enzyme I, the phosphocarrier protein (HPr), and Enzyme II. Enzyme I consists of an N-terminal domain (EIN) and a C-terminal domain (EIC). Enzyme II (EII) comprises three functional subunits EIIA^Glc^, EIIB^Glc^, and EIIC^Glc47, 48^. Weak interactions are responsible for the transferring of phosphate between EIN and HPr^49, 50^ and between EIIA^Glc^ and EIIB^Glc51^, and the *K*_D_ values for these two protein complexes are ~7 μM^52^ and ~25 μM^53^, respectively (Figure 4a).

**Figure 4.**
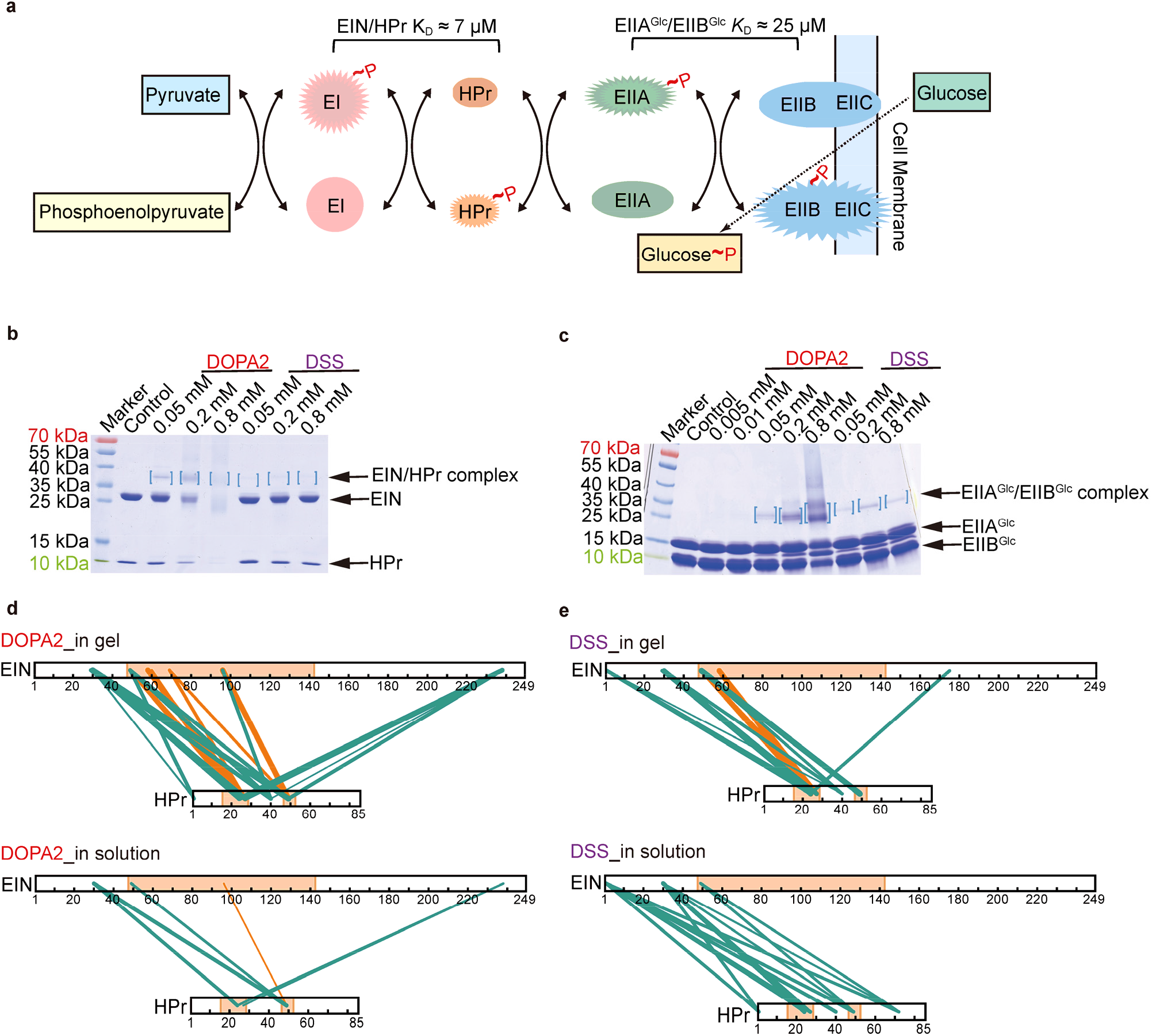
DOPA2 can capture weak interactions within the EIN/HPr and EIIA^Glc^/EIIB^Glc^ complexes. (a) Conceptual diagram of carbohydrate transport and phosphorylation by the phosphotransferase system. (b) SDS-PAGE of DOPA2 or DSS-cross-linked EIN/HPr. The concentration of EIN/HPr is 10× *K*_D_. Dimers are marked by square brackets. (c) As in b, but for the EIIA^Glc^/EIIB^Glc^ complex. (d) The inter-molecular residue pairs identified by DOPA2 in the dimer gel or in the solution were mapped on the primary sequence of EIN/HPr (visualized using xiNET^61^). (e) As in d, but for cross-links identified by DSS. Cross-links were filtered by requiring FDR < 0.01 at the spectra level, E-value < 1×10^−8^ and spectral counts > 3. The interactional interface of EIN and HPr on the specific complex was indicated by light orange color. The green lines and the orange lines denoted the cross-links that belong to encounter complexes and specific complex, respectively. The numbers of the corresponding spectra of each cross-link were indicated by the thickness of the line.

Consistent with the weak association between EIN and HPr and between EIIA^Glc^ and EIIB^Glc^, DSS cross-linking yielded no discernable covalent dimer bands on SDS-PAGE at a protein concentration of 0.25 mg/mL (for EIN/HPr) or 0.63 mg/mL (for EIIA^Glc^/EIIB^Glc^) (Supplementary Figure 7a-b). Note that the protein concentration used here are in the range for conventional cross-linking experiments. Interestingly, faint covalent dimer bands of EIIA^Glc^/EIIB^Glc^ appeared when the same samples were cross-linked with DOPA2 (Supplementary Figure 7b). In above experiments, the protein concentrations were set to about 1× *K*_D_, i.e., EIN and HPr each at 7 μM (0.06 mg/mL and 0.19 mg/mL, respectively), EIIA^Glc^ and EIIB^Glc^ each at 25 μM (0.40 mg/mL and 0.23 mg/mL, respectively). Under these conditions, the equilibrium concentrations of the heterodimers, EIN/HPr and EIIA^Glc^/EIIB^Glc^, were 2.67 μM and 9.55 μM, respectively. In other words, only 38% of the protein subunits were in the complex form. When the protein concentrations were increased to 10× *K*_D_, the complex percentage increased to 73%, which corresponds to an equilibrium concentration of 51.09 μM for the EIN/HPr heterodimer and of 182.46 μM for EIIA^Glc^/EIIB^Glc^. Thus, this is a 19-fold increase in the quantity of the protein complexes relative to that at 1× *K*_D_. At 10× *K*_D_ concentration, compared to cross-linking with DSS, cross-linking using DOPA2 generated more prominent dimer bands on SDS-PAGE for both complexes (Figure 4b-c and Supplementary Figure 8a-b). This validates the idea that the faster DOPA2 cross-linking is superior to DSS cross-linking in capturing weak protein-protein interactions.

LC-MS/MS analysis of the cross-linked dimers from the gel bands showed that the DOPA2 samples contain more intra- and inter-molecular cross-links than the DSS samples (Supplementary Figure 8c-d). For example, there were 34 DOPA2 and 21 DSS inter-molecular cross-links between EIN and HPr, and 37 DOPA2 and 24 DSS inter-molecular cross-links between EIIA^Glc^ and EIIB^Glc^, respectively (Supplementary Figure 8c-d). Consistent with the results obtained from six model proteins (Figure 1d-e), a higher percentage of the DOPA2-linked residue pairs are compatible with the crystal structures (Supplementary Figure 8e-g).

We showed in our previous work^54^ that a vast majority of the EIN/HPr inter-molecular cross-links generated by BS^2^G or BS^3^, both NHS ester cross-linkers, reflected ECs, which are a collection of conformations with higher energy and less specific electrostatic attractions between the two protein partners compared to the finally formed complex. These fleeting ECs are too weak to be captured by crystallization, but are detectable by a special NMR method^55^. The lower energy-state stereospecific complex (SC) of EIN/HPr as depicted in the NMR structure is mainly stabilized by the stereospecific hydrogen bonds and hydrophobic interactions^49, 55^. Only one out of the 13 inter-molecular BS^2^G or BS^3^ cross-links comes from SC^54^.

In this study, we obtained a higher percentage of cross-links representing SC by the use of either DOPA2 cross-linking or the gel band containing covalently linked EIN/HPr. When the EIN/HPr samples were analyzed by LC-MS/MS without SDS-PAGE separation (in-solution samples) as done before^54^, one of the six inter-molecular DOPA2 cross-links fitted with SC (16.7%); in contrast, none of the 13 inter-molecular DSS cross-links fitted with SC (Figure 4d-e, Supplementary Table 6 and Supplementary Table 7a-b). From the gel band of cross-linked EIN/HPr, four (27%) out of the 15 inter-molecular DSS cross-links represented SC (Figure 4e, Supplementary Table 6 and Supplementary Table 7c). Of note is that the cross-links examined here all passed a stringent filter as descried above. We estimated that the SC-specific cross-links were enriched by DOPA2 and gel band by factors of about 3 and 2, respectively (Supplementary Table 6). As the SC cross-links are already favored by the use of DOPA2, further increase by analyzing the gel band became less obvious (Figure 4d-e and Supplementary Table 6). Nevertheless, LC-MS/MS analysis of the gel bands containing DOPA2-linked EIN/HPr yielded the best result as 8 out of 25 (32%) inter-molecular cross-links represented SC (Figure 4d, Supplementary Table 6 and Supplementary Table 7d). Together, the data above show that DOPA2 outperforms DSS in capturing weak but specific protein-protein interactions.

## Discussion

In this study, we have developed an amine-selective non-hydrolyzable di-*ortho*-phthalaldehyde cross-linker, DOPA2. It offers fast cross-linking under extreme conditions (e.g., 8 M urea or 6 M GdnHCl) where the commonly used NHS ester cross-linker is ineffective. Our results demonstrate that DOPA2-based CXMS could be a valuable technique to extract information about conformational states, including partially and fully unfolded states, and about continuous conformational changes associated with protein unfolding. We also demonstrate that under the typical, near physiological conditions, DOPA2 is advantageous over NHS ester cross-linkers in capturing weak but specific protein-protein interactions.

In using DOPA2, one thing to watch out for is not to over-cross-link with it. Because DOPA2 does not hydrolyze, its optimal concentration in protein cross-linking reactions is three- or four-fold lower than that of the NHS ester cross-linker (Figure 2a and Supplementary Figure 3b). When initiating a cross-linking reaction, the protein solution should be mixed with an equal volume of 2× DOPA solution to avoid high local concentration. If proteins are cross-linked using a slightly higher than optimal concentration of DOPA2 (e.g., at 4:1 instead of 16:1 protein/DOPA2, w/w, for BSA), the sample can be digested with a second protease Asp-N following trypsin (Supplementary Figure 3d) to generate shorter, more readily identifiable peptides.

DOPA2 cross-links proteins very fast (Figure 2a and Supplementary Figure 4). We succeeded in monitoring RNase A unfolding in time intervals as short as five seconds by subjecting each time point sample to DOPA2 cross-linking for only 10 seconds (Figure 3e). It seems that 10 seconds are the minimum reaction time. As shown in Supplementary Figure 4, when the reaction time is reduced from 10 minutes to 5 seconds, we see an overwhelming amount of mono-links and a small number of cross-links. This is understandable because in most or all cross-linking reactions, one end of a cross-linker makes a covalent attachment before the other. Extrapolating out the trend line, it is predictable that sub-second DOPA2 cross-linking will generate few cross-links.

As shown in Figure 2a, cross-linking of BSA by DOPA2 is ~60-fold faster than that by DSS. An earlier study reports that covalent conjugation of OPA to a peptide (equivalent to a mono-link of DOPA) is about eight times as fast as an NHS ester^40^. In our hands, the pseudo-first-order reaction rate constant between OPA and free lysine is too fast to measure. Even when the ratio of free lysine:OPA is adjusted to 1:2, the second-order rate constant is still too fast to be measured accurately with the equipment we have (Supplementary Figure 1b, the amount of product approached the maximum at 30 s). What we can say for certain is that OPA reacts markedly faster than an NHS ester. Possibly in a protein cross-linking reaction, which involves two consecutive OPA or NHS ester reactions, success of the first reaction likely accelerates the second one by reducing the degree of freedom, i.e. creating a proximity effect. This could account for the larger difference in the reaction speed between DOPA and DSS than that between OPA and an NHS ester.

One intriguing result of our study is the differential preferences of DOPA2 and DSS towards SC and EC, respectively. Namely, for the weak EIN/HPr complex, the relatively slow cross-linking reagent DSS prefers fleeting ECs whereas the faster cross-linking reagent DOPA2 prefers the longer-lived SC. The possible reasons are discussed below.

The process of capturing a protein-protein interaction by cross-linking may be conceptualized as follows. We start from the moment when one end of the cross-linker already is covalently attached to either of the two protein partners (A and B) that participate in the formation of the final complex A•B. This simplifies the second step of cross-linking to a near first-order reaction, because the attachment reaction by one end of a cross-linker increases its effective local concentration and hence accelerates a subsequent cross-linking reaction by the other end. Assuming that the cross-linker lands on a site that allows inter-protein cross-linking and that everything else is equal, then the likelihood for the formation of an inter-protein cross-link will be determined by the reaction rate of the amine-targeting group and the half-life of the A•B complex. If the reaction rate is not fast enough compared to the half-life of the A•B complex, no inter-protein cross-link will form for this round of association. However, the cross-linker that has been planted onto a protein partner by one end still has a chance to form an inter-protein cross-link in the next round of association before it loses reactivity due to the attack of an intra-molecular amine group or a quenching reagent.

We propose that DSS or BS^3^ cross-linking reactions are too slow to capture either the SC or the fleeting ECs in one round of association/dissociation cycle, and the observed cross-links may come from a later association event between a planted subunit and its partner. As an association event starts presumably with nonspecific electrostatic attractions that could result in ECs, it is not surprising that a single-end attached DSS or BS^3^ cross-linker is able to capture such conformations. This is in line with an earlier work demonstrating that a slow reacting sulfonyl fluoride group can capture weak and transient interactions once planted onto a protein^56^. This explanation fits with the observation that the number of inter-molecular DSS or BS^3^ cross-links identified from EIN/HPr is small and most of them represent ECs. In contrast, DOPA2 has a faster reaction rate and thus is more efficacious in capturing SC before it dissociates. Because ECs exist as an ensemble of distinct conformations while SC represents a stable conformational state, cross-linked ECs are more heterogeneous than cross-linked SC, thus explaining why ECs are less visible on SDS-PAGE as they spread along the lane and why the visible covalent complex enriches for SC cross-links (Supplementary Table 6).

In summary, DOPA not only provides a powerful tool for CXMS to identify weak protein-protein interactions, but also introduces CXMS to new application areas such as studying the conformational changes during protein unfolding.

## Materials and methods

### Chemical instrumentation and methods

All reactions were carried out in oven-dried glassware under an argon atmosphere, unless otherwise stated. Air and moisture sensitive reagents were transferred by syringe or cannula. Brine refers to a saturated aqueous solution of NaCl. Analytical thin layer chromatography (TLC) was performed on 0.25 mm silica gel 60-F plates, using 254 nm UV light as the visualizing agent or staining with potassium permanganate or phosphomolybdic acid and heat as developing agents. Flash chromatography was performed using 200-400 mesh silica gel. Yields refer to chromatographically and spectroscopically pure materials, unless otherwise stated.

^1^H NMR spectra were recorded on a Varian 400 or 500 MHz spectrometer at ambient temperature with CDCl_3_ as the solvent, unless otherwise stated. ^13^C NMR spectra were recorded on a Varian 100 or 125 MHz spectrometer (with complete proton decoupling) at ambient temperature. Chemical shifts are reported in parts per million relative to chloroform (^1^H, δ 7.26 ppm; ^13^C, δ 77.16 ppm). Data for ^1^H NMR are reported as follows: chemical shift, integration, multiplicity (s = singlet, d = doublet, t = triplet, q = quartet, m = multiplet) and coupling constants. High-resolution mass spectra were obtained at Peking University Mass Spectrometry Laboratory using a Bruker APEX instrument.

Synthetic compounds were analyzed by UPLC/MS on a Waters UPLC H Class and SQ Detector 2 system. The system was equipped with a Waters C18 1.7 μm Acquity UPLC BEH column (2.1*50 mm), equilibrated with HPLC grade water (solvent A) and HPLC grade acetonitrile (solvent B) with a flow rate of 0.3 mL/min.

### Reagents and solvents

All chemical reagents were from J&K, Alfa Aesar or TCI Chemicals without further purification unless otherwise stated. DCM and CH3CN were distilled from calcium hydride; THF was distilled from sodium/benzophenone ketyl prior to use.

For the MS analysis, DSS, tris(2-carboxyethyl) phosphine (TCEP), and 2-Iodoacetamide (IAA) were purchased from Pierce biotechnology (Thermo Scientific). Guanidine hydrochloride (GdnHCl) was purchased from MP Biomedicals LLC. Dimethylsulfoxide (DMSO), HEPES, NaCl, KCl, urea, CaCl_2_, methylamine, and other general chemicals were purchased from Sigma-Aldrich. Acetonitrile, formic acid, acetone, and ammonium bicarbonate were purchased from J.T. Baker. Trypsin, Asp-N (gold mass spectrometry grade) was purchased from Promega.

### Preparation of protein samples

Aldolase, BSA, catalase, lysozyme, myosin, lactoferrin, carbonic anhydrase 2, and β-amylase were obtained from Sigma-Aldrich. RNase A was purchased from Thermo Fisher. Recombinant GST containing an N-terminal His tag was expressed in *E. coli* BL21 cells from the pDYH24 plasmid and purified with glutathione sepharose (GE Healthcare). PUD-1/PUD-2 heterodimers were purified on a HisTrap column followed by gel filtration. Stock solutions of model proteins were individually buffer exchanged into 20 mM HEPES, pH 8.0 by ultrafiltration. The mixture of ten model proteins was mixed manually and contained: aldolase, BSA, catalase, carbonic anhydrase 2, lysozyme, lactoferrin, β-amylase, myosin, GST and PUD-1/PUD-2. Each of the ten proteins was present at a final concentration of 0.1 mg/mL (total protein concentration, 1 mg/mL).

The N-terminal domain of *E. coli* enzyme I (EIN, residues 1-249) and the histidine-containing phosphorcarrier protein HPr, EIIA^Glc^ and EIIB^Glc^ were purified as previously described^49, 57^. Eluted proteins were exchanged into 20 mM HEPES, 150 mM NaCl, pH 7.4.

### Kinetics experiments for second-order rate constant determination

1. Equation of calculating the second-order rate constants

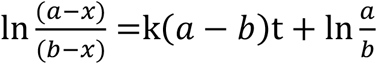 **a** represented the amount of OPA/NHS ester analogue in the beginning. **b** represented the amount of Boc-OMe-lysine in the beginning. **x** represented the consumption of OPA/NHS ester analogue/Boc-OMe-lysine at time **t**. (The consumption is equivalent) **(a-x)** represented the amount of remaining OPA/ NHS ester analogue at time **t**. This amount is equal to the amount of quenched product. **(b-x)** represented the amount of remaining Boc-OMe-lysine at time **t**. **k** represented the second-order reaction kinetic constants.
2. Kinetics experiments of Boc-OMe-lysine with OPA Boc-OMe-lysine (5 μmol, 1 eq.) was dissolved in 10 mL TEAB buffer at 25 °C. OPA (10 μmol, 2 eq.) was added in the stirred reaction mixture. The reaction was monitored at 0.5 min, 1 min, 1.5 min, 2 min, 2.5 min, 3 min, 3.5 min, 4 min. At each time point, 10 μl reaction mixture was quenched by 1 μl 98% hydrazine. The conversion was monitored by LC-MS.
3. Kinetics experiments of Boc-OMe-lysine with NHS ester analogue Boc-OMe-lysine (5 μmol, 1 eq.) was dissolved in 10 mL TEAB buffer at 25 °C. NHS (10 μmol, 2 eq.) was added in the stirred reaction mixture. The reaction was monitored at 2 min, 3 min, 4 min, 5 min, 6 min, 7 min, 8 min, 9 min. At each time point, 10 μl reaction mixture was quenched by 1 μl 50% hydroxylamine. The conversion was monitored by LC-MS.

### Characterization of OPA selectivity towards different amino acids

Ten synthesized peptides (listed in Table S1) were dissolved in 20 mM HEPES, pH 7.4 at a final concentration of 2 mM. OPA was added at a final concentration of 2 mM. After one-hour reaction at room temperature, the reaction products were analyzed by LC-MS/MS.

### Peptide cross-linking

The synthesized peptides VR-7 (sequence: VWDLVKR), KR-7 (sequence: KMRPEVR), TR-8 (sequence: TPDVNKDR) and GR-11 (sequence: (N,N-dimethyl-Gly)-VAAAKAAAAR) were dissolved in 20 mM HEPES, pH 7.4 at a final concentration of 2 mM. Cross-linker DOPA2 was added at a final concentration of 2 mM. For testing the activity of DOPA2 under acidic pH or in the presence of high concentrations of denaturants, peptides VR-7, KR-7, TR-8, GR-11 were dissolved in 100 mM citric acid-Na_2_HPO_4_, pH3.0 or in 6 M GdnHCl at a final concentration of 2 mM. Cross-linker DOPA2 was added at a final concentration of 2 mM. After cross-linking at room temperature for one hour, samples were diluted and about 5 pmol of total peptides were subjected to liquid chromatography coupled with tandem mass spectrometry (LC-MS/MS) analysis.

### Protein cross-linking

The cross-linking conditions, reaction time, reaction temperature, reaction buffers, concentrations of cross-linkers were optimized using protein BSA at the final concentration of 1 mg/mL. Six model proteins and the ten-protein mixture were diluted to 1 mg/mL with HEPES buffer (20 mM HEPES and 150 mM NaCl, pH 7.4) and were cross-linked with DOPA-C_2_, DOPA1 or DOPA2 (the protein-to-cross-linker mass ratio at 16:1) at room temperature for 10 minutes.

The reactivity of cross-linkers DOPA2 and DSS in different pH conditions was evaluated with protein BSA. Protein BSA were dissolved in citric acid-Na_2_HPO_4_ buffers under pH 3.0, pH 4.0, pH 5.0, pH 6.0, pH 7.0, pH 7.4, respectively and then were cross-linked with DOPA2 (0.17 mM) or DSS (0.5 mM) for 10 s.

The reactivity of cross-linkers DOPA2 and DSS in different concentrations of denaturants was evaluated with protein BSA. Protein BSA were in the urea (0 M, 1 M, 2 M, 4 M, 6 M, 8 M) at room temperature for one hour, and then were cross-linked with DOPA2 (0.17 mM) or DSS (0.5 mM) for 10 s. In the GdnHCl (0 M, 1 M, 2 M, 3 M), BSA were cross-linked with DOPA2 (0.17 mM) or DSS (0.5 mM) at room temperature for 10 s, 1 min, 6 min, respectively. All the reactions were quenched with 20 mM hydrazine at room temperature for 5 minutes. The cross-linking reaction of RNase A in different concentrations of urea (0 M, 1 M, 2 M, 4 M, 6 M, 8 M) or GdnHCl (0 M, 1 M, 2 M, 3 M, 4 M, 5 M, 6 M) is similar to that of protein BSA. Protein RNase A were in the urea for one hour and in the GdnHCl for ten hours before cross-linking. Reactions were quenched as described above. After cross-linking, denaturants were diluted to the final concentration of 1.5 M with 1× PBS buffer, and then cross-linked proteins were precipitated with 4-fold volume of cool acetone.

For the cross-linking reaction of RNase A in 8 M urea at different time points, 20 μl 10 mg/mL RNase A was mixed with 160 μl 10 M urea solution and 20 μl 1× PBS buffer, pH 7.4, and the timer started timing. 20 μl protein sample was picked up quickly at the time points 5 s, 10 s, 20 s, 30 s, 1 min, 2 min, 5 min, 10 min, 15 min, 30 min, and the reactions were terminated with 20 mM hydrazine at room temperature for 5 minutes. (Note: the first four time points are too dense, and it is difficult to pick up samples continuously and control the time strictly. Thus, the samples of RNase A in 8 M urea were prepared one by one before the addition of cross-linkers.)

The *K*_D_ value of EIN/HPr complex was ~7 μM^52^. EIN/HPr complexes were diluted to 1× *K*_D_ (0.25 mg/mL) and 10× *K*_D_ (2.5 mg/mL), and then were cross-linked with DOPA2 at the final concentration of 0.05 mM, 0.2 mM, 0.8 mM at room temperature for 10 minutes, respectively. EIN/HPr complexes were also cross-linked with DSS at the final concentration of 0.05 mM, 0.2 mM, 0.8 mM at room temperature for one hour.

The *K*_D_ value of EIIA^Glc^/EIIB^Glc^ complex was estimated to be ~25 μM. According to a previous study, EIIA^Glc^/EIIB^Glc^ has a *K*m value of 1.7-25 μM^53^. We did not succeed in obtaining an accurate *K*_D_ value by either surface plasmon resonance assay or ELISA. However, it is clear that the binding is weak and the *K*_D_ value is likely greater than 10 μM. EIIA^Glc^/EIIB^Glc^ complexes were diluted to 1× *K*_D_ (0.63 mg/mL) and 10× *K*_D_ (6.3 mg/mL), and then were cross-linked with DOPA2 at the final concentration of 0.005 mM, 0.01 mM, 0.05 mM, 0.2 mM, 0.8 mM at room temperature for 10 minutes, respectively. EIIA^Glc^/EIIB^Glc^ complexes were also cross-linked with DSS at the final concentration of 0.05 mM, 0.2 mM, 0.8 mM at room temperature for one hour.

### Trypsin digestion

Cross-linked proteins were precipitated with 4-fold volume of acetone for at least 30 minutes at −20 °C. The pellets were air dried and then dissolved, assisted by sonication, in 8 M urea, 20 mM methylamine, 100 mM Tris, pH 8.5. After reduction (5 mM TCEP, RT, 20 min) and alkylation (10 mM iodoacetamide, RT, 15 min in the dark), the samples were diluted to 2 M urea with 100 mM Tris, pH 8.5. Denatured proteins were digested by trypsin at a 1/50 (w/w) enzyme/substrate ratio at 37 °C for 16-18 h, and the reactions were quenched with 5% formic acid (final conc.).

### LC-MS analysis

All synthesized peptide samples were analyzed using an EASY-nLC 1000 system (Thermo Fisher Scientific) interfaced with a Q-Exactive HF mass spectrometer (Thermo Fisher Scientific). Peptides were loaded on a pre-column (75 μm ID, 4 cm long, packed with ODS-AQ 12 nm-10 μm beads) and separated on an analytical column (75 μm ID, 12 cm long, packed with Luna C18 1.9 μm 100 Å resin). Slight modifications to the separation method were made for different samples. The ten OPA modified peptides were injected and separated with a 30 min linear gradient at a flow rate of 200 nl/min as follows: 0-5% B in 2 min, 5-30% B in 15 min, 30-100% B in 3 min, 100% B for 10 min (A = 0.1% FA, B = 100% ACN, 0.1% FA). The DOPA2 cross-linked peptides VR-7, KR-7, TR-8 and GR-11 were injected and separated with a 30 min linear gradient at a flow rate of 200 nl/min as follows: 0-5% B in 2 min, 5-28% B in 15 min, 28-100% B in 3 min, 100% B for 10 min (A = 0.1% FA, B = 100% ACN, 0.1% FA). Spectra were acquired in data-dependent mode: the top fifteen most intense precursor ions from each full scan (resolution 60,000) were isolated for HCD MS2 (resolution 15,000, NCE 27) with a dynamic exclusion time of 30 s. Precursors with more than 6+, or unassigned charge states were excluded. In order to acquire more high-quality spectra at the peaks, ten OPA modified peptides were also analyzed with a dynamic exclusion time of 10 s.

All protein samples were analyzed using an EASY-nLC 1000 system (Thermo Fisher Scientific) interfaced with an HF Q-Exactive mass spectrometer (Thermo Fisher Scientific). Peptides were loaded on a pre-column and separated on an analytical column as noted above. Slight modifications to the separation method were made for different samples. The BSA samples for optimizing the cross-linking conditions, a ten-protein mixture and RNase A were injected and separated with a 60 min linear gradient at a flow rate of 200 nl/min as follows: 0-5% B in 2 min, 5-30% B in 43 min, 30-100% B in 5 min, 100% B for 10 min (A = 0.1% FA, B = 100% ACN, 0.1% FA). EIN/HPr and EIIA^Glc^/EIIB^Glc^ complexes were injected and separated with a 75 min linear gradient at a flow rate of 200 nl/min as follows: 0-5% B in 1 min, 5-35% B in 59 min, 35-100% B in 5 min, 100% B for 10 min (A = 0.1% FA, B = 100% ACN, 0.1% FA). The top fifteen most intense precursor ions from each full scan (resolution 60,000) were isolated for HCD MS2 (resolution 15,000; NCE 27) with a dynamic exclusion time of 30 s. Precursors with 1+, 2+, more than 6+, or unassigned charge states were excluded.

### Identification of cross-links with pLink 2^46^

The search parameters used for pLink 2 were as follows: instrument, HCD; precursor mass tolerance, 20 ppm; fragment mass tolerance 20 ppm; cross-linker DOPA-C_2_ (cross-linking sites K and protein N-terminus, cross-link mass-shift 258.068, mono-link w/o hydrazine mass-shift 276.079, mono-link w/t hydrazine mass-shift 272.095); cross-linker DOPA1 (cross-linking sites K and protein N-terminus, cross-link mass-shift 290.058, mono-link w/o hydrazine mass-shift 308.068, mono-link w/t hydrazine mass-shift 304.085), cross-linker DOPA2 (cross-linking sites K and protein N-terminus, cross-link mass-shift 334.084, mono-link w/o hydrazine mass-shift 352.096, mono-link w/t hydrazine mass-shift 348.111), cross-linker DSS (cross-linking sites K and protein N-terminus, cross-link mass-shift 138.068, mono-link mass-shift 156.079); fixed modification C 57.021; peptide length, minimum 6 amino acids and maximum 60 amino acids per chain; peptide mass, minimum 600 and maximum 6,000 Da per chain; enzyme, trypsin or trypsin and Asp-N, with up to three missed cleavage sites per cross-link. Protein sequences of model proteins were used for database searching. The results were filtered by requiring a spectral false identification rate < 0.01. MS2 spectra were annotated using pLabel^58, 59^, requiring mass deviation ≤ 20 ppm.

### Ca-Ca distance calculations

The Cα-Cα Euclidean distances were calculated by an in-house Perl script with the coordinates from the PDB files. The Cα-Cα Solvent Accessible Surface Distance (SASD) were calculated using Jwalk^60^. For model proteins, it is not possible to distinguish from the sequence of the peptide whether the cross-links are intra- or inter-molecular. We thus calculated all the possible combinations and picked those with the shortest Cα-Cα distance. If the SASD of cross-linked residue pairs cannot be calculated due to a lack of surface accessibility, these residue pairs are excluded from calculation. When calculating structural compatibility, the distance cut-offs are 24.9 Å for DOPA-C_2_, 27.7 Å for DOPA1, 30.2 Å for DOPA2, and 24.0 Å for DSS.

### K-means clustering and data normalization

The spectral counts of each cross-linked site pair across all samples were first transformed to Z-Score (mean centered and scaled to the variance). K-means clustering was performed using Cluster 3.0 with the settings of 3 clusters and a maximum of 100 iterations.

### Circular dichroism (CD) measurements

CD spectra were measured on a Chirascan-plus circular dichroism spectrometer. Measurements were done at 20 °C and pH 7.4. Data were recorded between 210-260 nm and quartz cuvette with 1 mm path length. Measurements were acquired in 1 nm increments with an integration time of 0.5 s. RNase A was measured at 7 μM concentration and in the presence of different concentrations of urea (0 M-8 M) and GdnHCl (0 M-6 M). Three scans were averaged for each measurement. The averaged CD spectra for one condition and its respective buffer were subtracted and then smoothed (window size = 2). The resulting CD spectra were plotted with OriginPro 8.

## Acknowledgements

We thank Yulu Li and Dr. Jianhua Sui for trying very hard to determine the *K*_D_ value of EIIA^Glc^/EIIB^Glc^. We gratefully acknowledge financial support from the National Natural Science Foundation of China Grant (21625201, 21961142010, 21661140001, 91853202, and 21521003 to X.L.), the National Key Research and Development Program of China (2017YFA0505200 X.L.), the Beijing Outstanding Young Scientist Program (BJJWZYJH01201910001001 X.L.), the municipal government of Beijing (in the form of NIBS intramural grants), TIMBR, and Tsinghua University, the Program for Donglu Scholar in the Yunnan University.

## Author contributions

J.-H.W. performed MS experiments, analyzed and interpreted the data, and wrote the manuscript; Y.-L.T. designed and synthesized the compounds, analyzed the data, and wrote the manuscript; M.-Q.D. guided the study, interpreted the data, and wrote the manuscript; X.L., conceived the DOPA cross-liner, guided the study, and revised the manuscript; R.J. did CD analysis, helped with data analysis and interpretation, revised the manuscript; F.X. performed the kinetics experiments; Z.G., X.D., and C.T. provided the EIN/HPr and EIIA^Glc^/EIIB^Glc^ protein complexes, helped with data analysis and interpretation; Y.Z. and N.H. helped with data analysis and interpretation; D.T. performed MS experiments; Q.L. performed the chemical synthesis; S.-Q.L. helped with data analysis and interpretation, revised the manuscript; K.Y. helped with data analysis and interpretation.

---

**Scheme 1.**
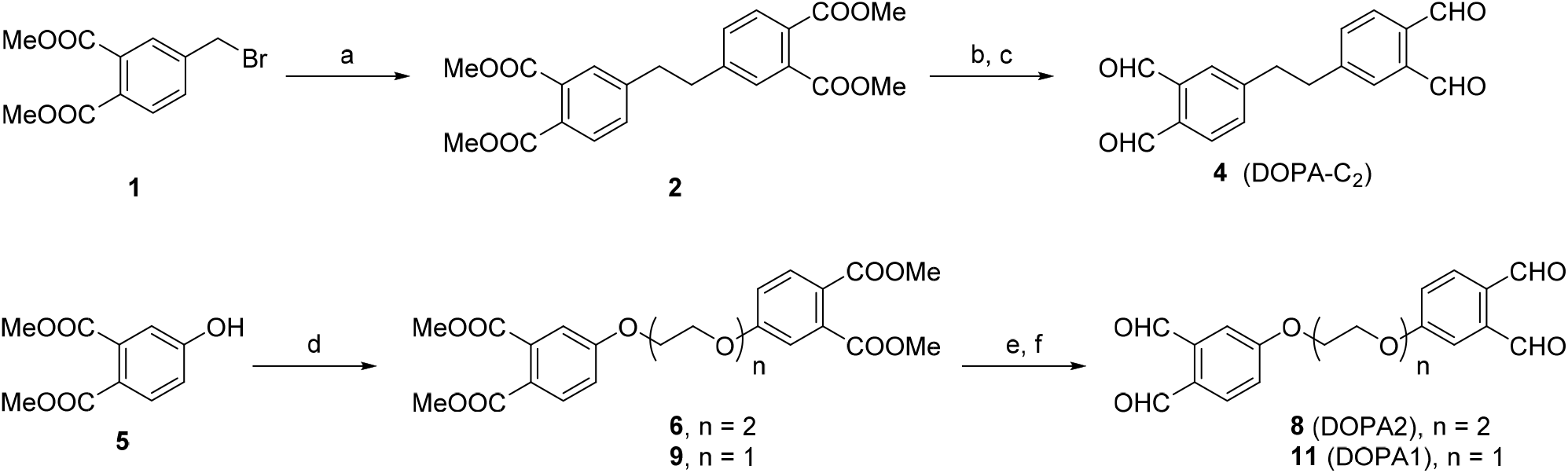
Syntheses of <u>d</u>i-*<u>o</u>rtho*-<u>p</u>hthal<u>a</u>ldehyde (DOPA) cross-linkers^a^. ^a^Reagents and conditions: (a) Me_2_Zn, RhCl(PPh_3_)_3_, THF, 24 h, 76%. (b) DIBAL-H, THF, 12 h. (c) DMP, DCM, 12 h, 98% for two steps. (d) Cs_2_CO_3_, bis (2-iodoethyl) ether or 1,2-dibromoethane, ACN, 80 °C, 12 h, n = 2, 77%; n = 1, 37%. (e) DIBAL-H, THF, 12 h. (f) DMP, DCM, 12 h, n = 2, 93% for two steps; n = 1, 80% for two steps. DIBAL-H = diisobutylaluminum hydride, DMP = Dess-Martin periodinane.

## Supplementary Information

**Supplementary Figure 1.**
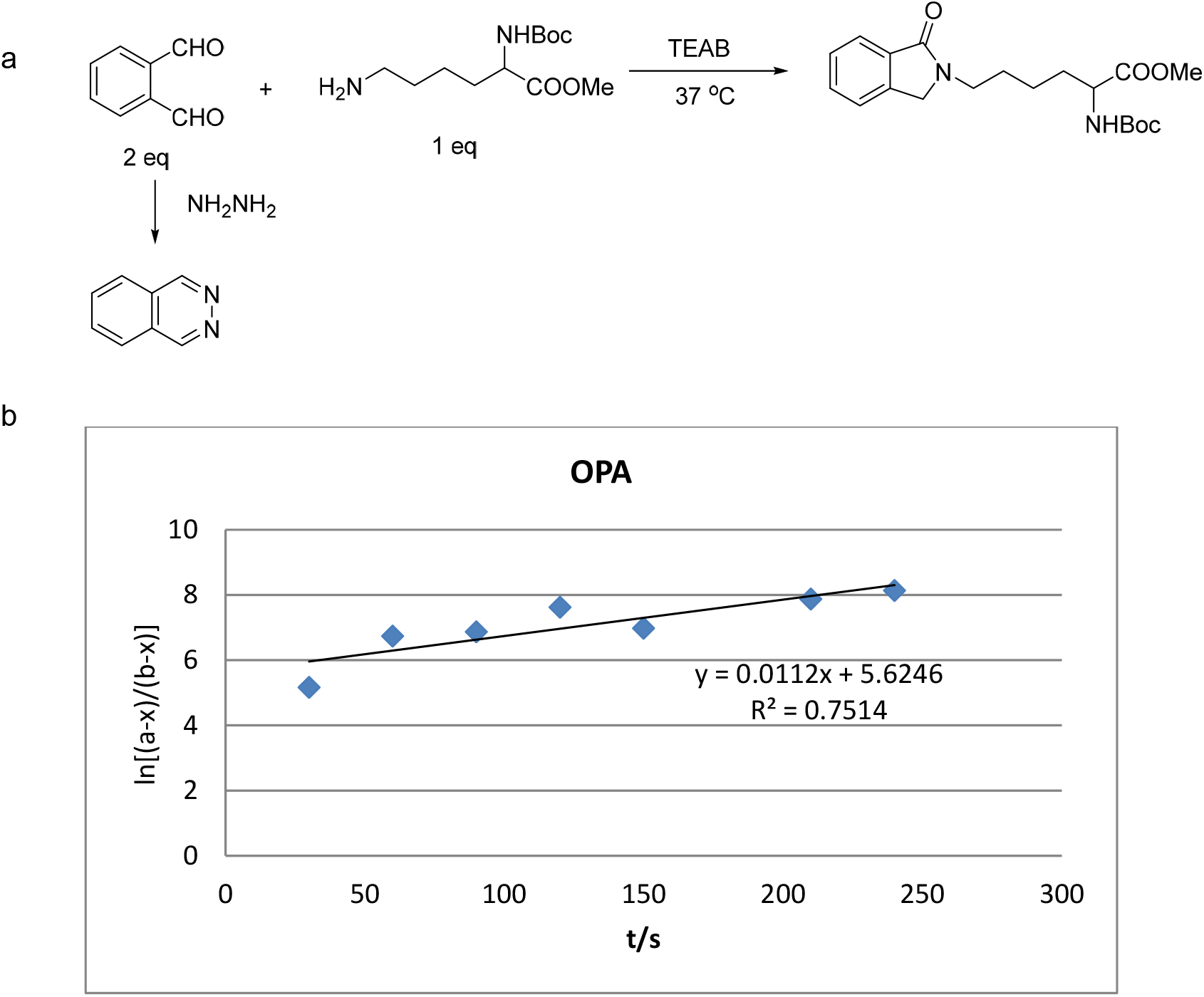
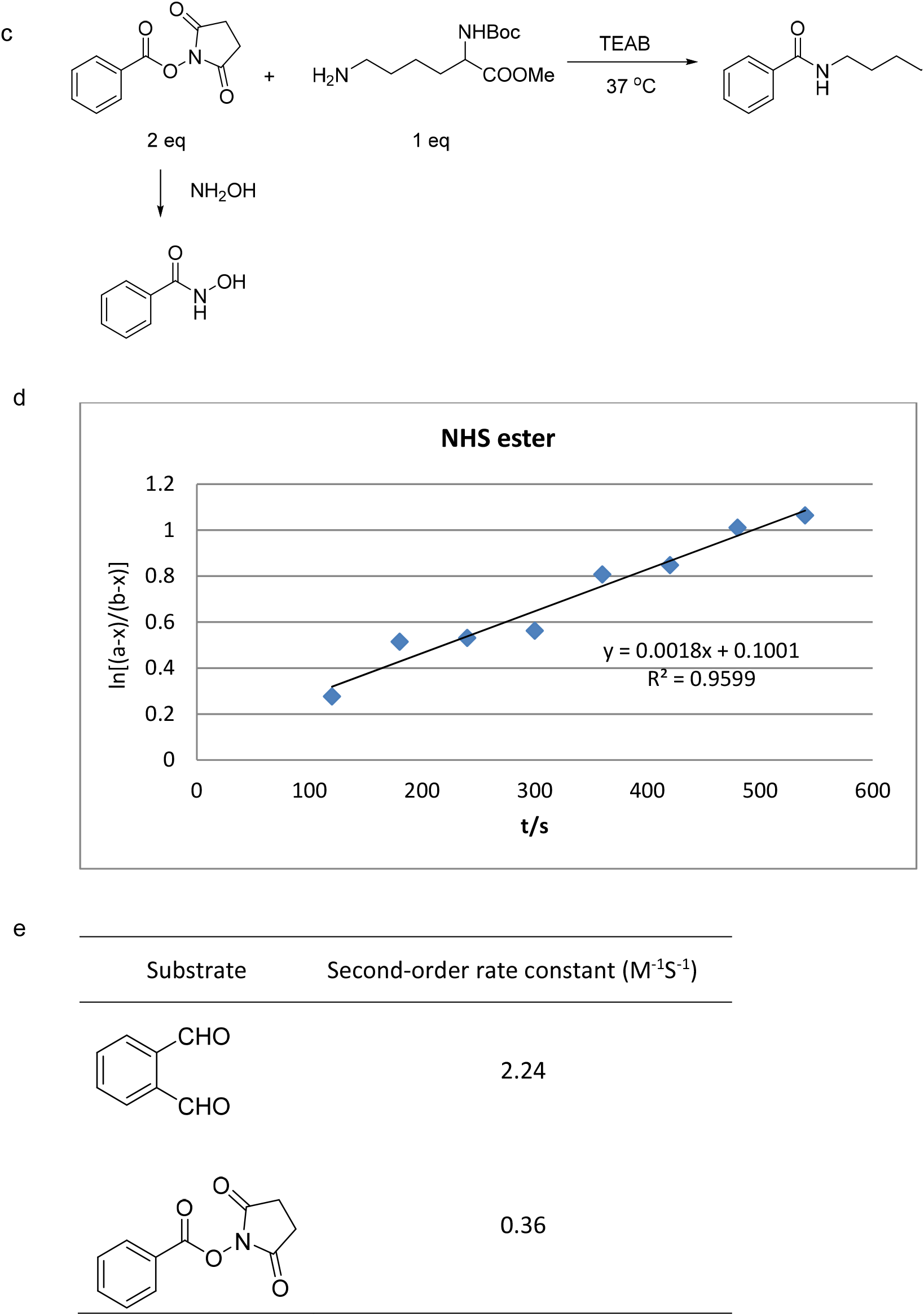
Rate kinetics of OPA and NHS ester with Boc-OMe-lysine. (a) The reaction of OPA with lysine afforded the phthalimidine and the excessive OPA was quenched by hydrazine to afford phthalazine. (b) The rate curve of the reaction of OPA with lysine. (c) The reaction of NHS ester with lysine afforded the amide and the excessive NHS ester was quenched by hydroxylamine. (d) The rate curve of the reaction of NHS ester with lysine. (e) The second-order rate constant of OPA and NHS ester with lysine.

**Supplementary Figure 2.**
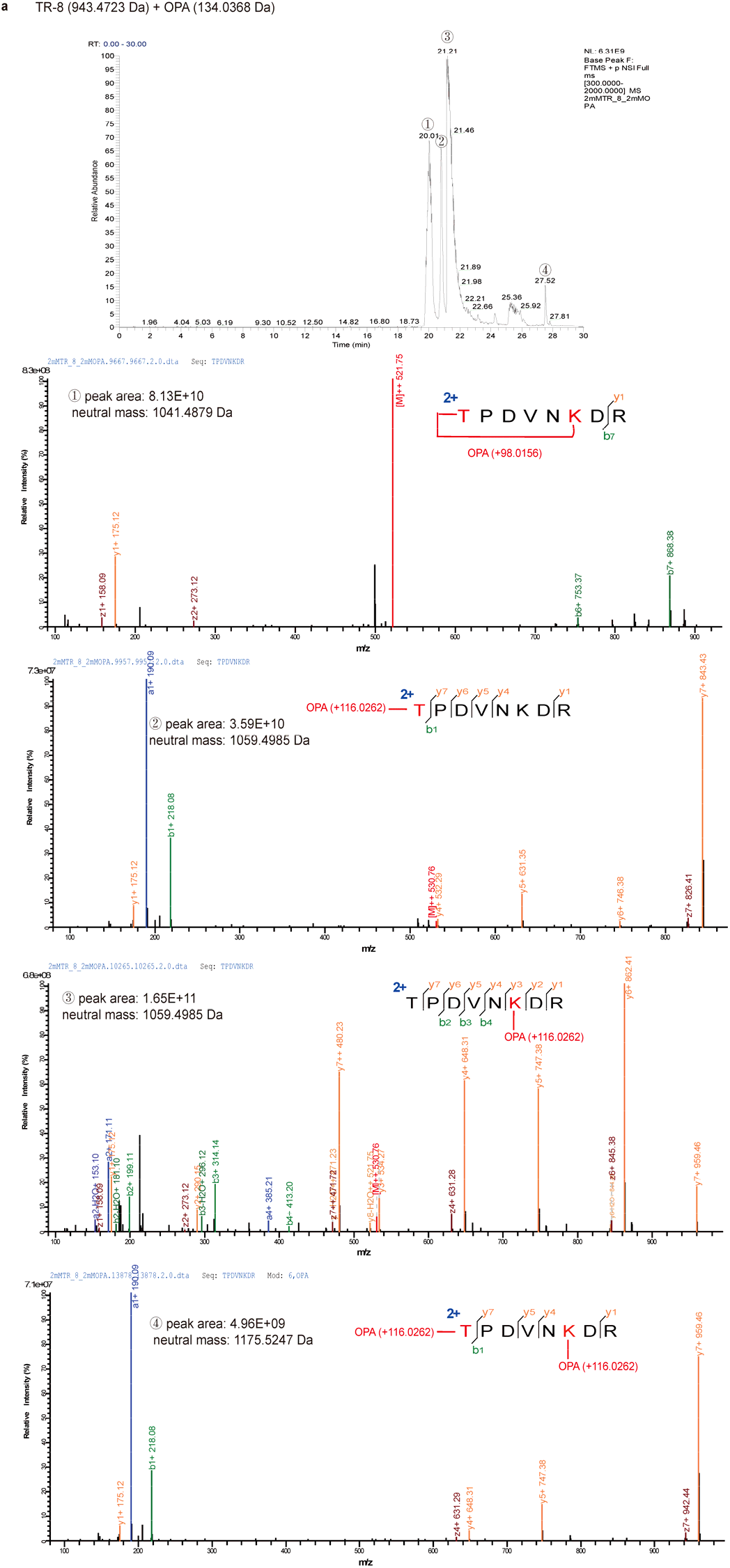
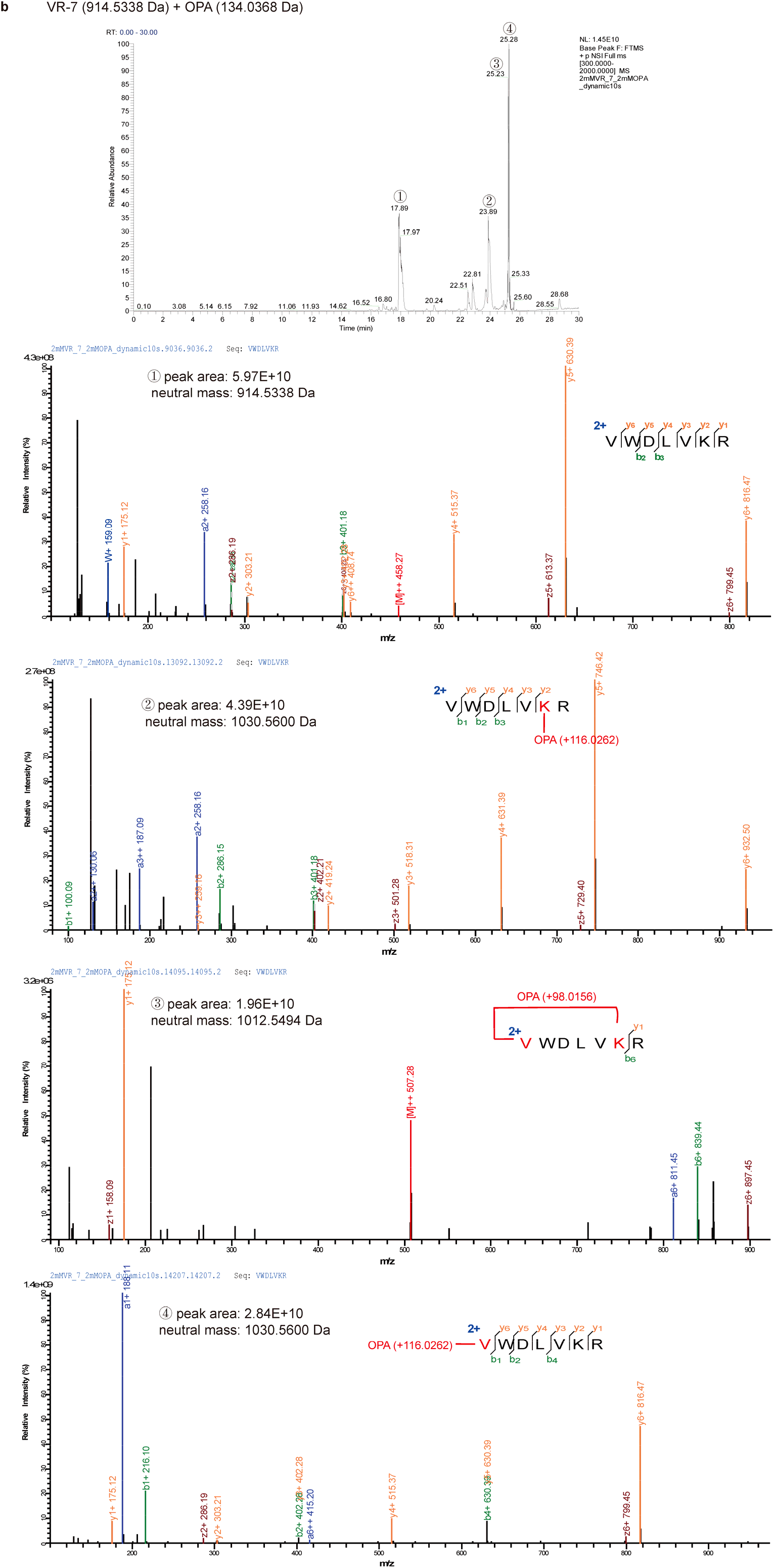
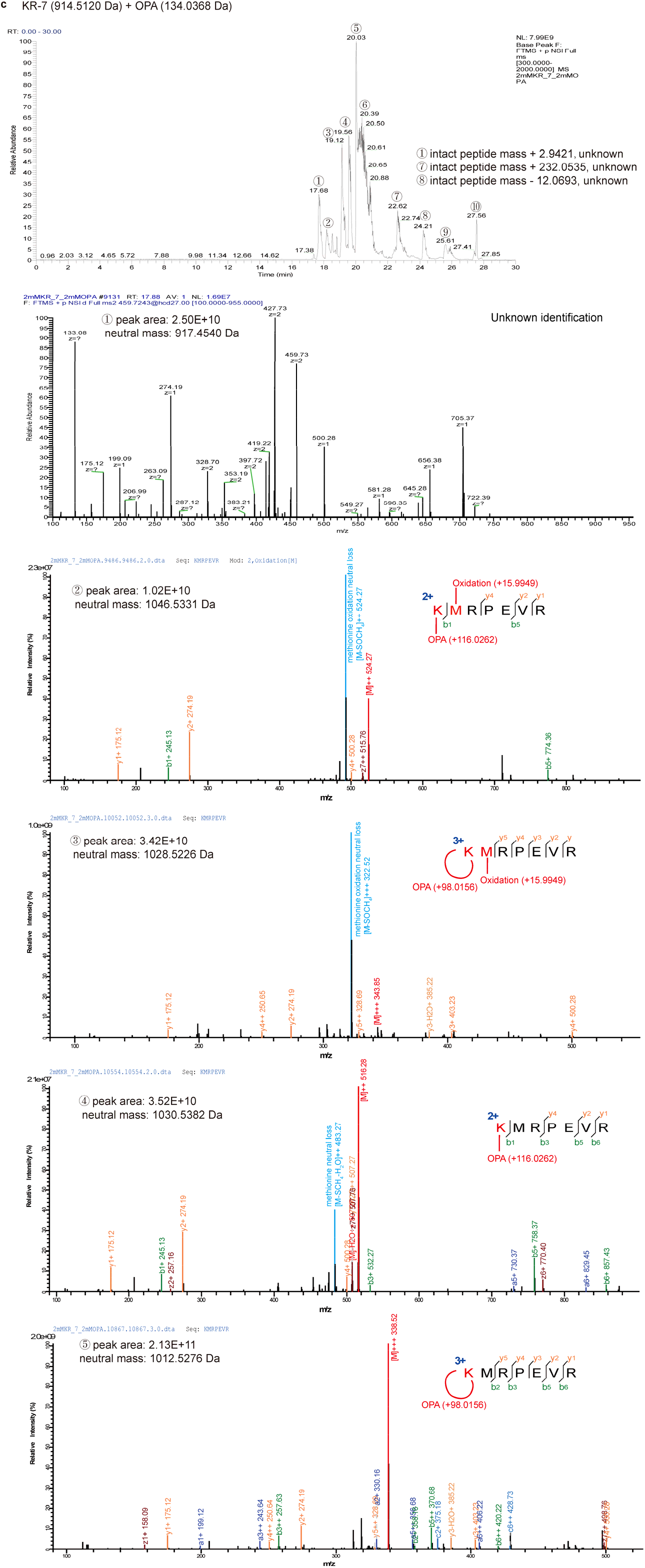
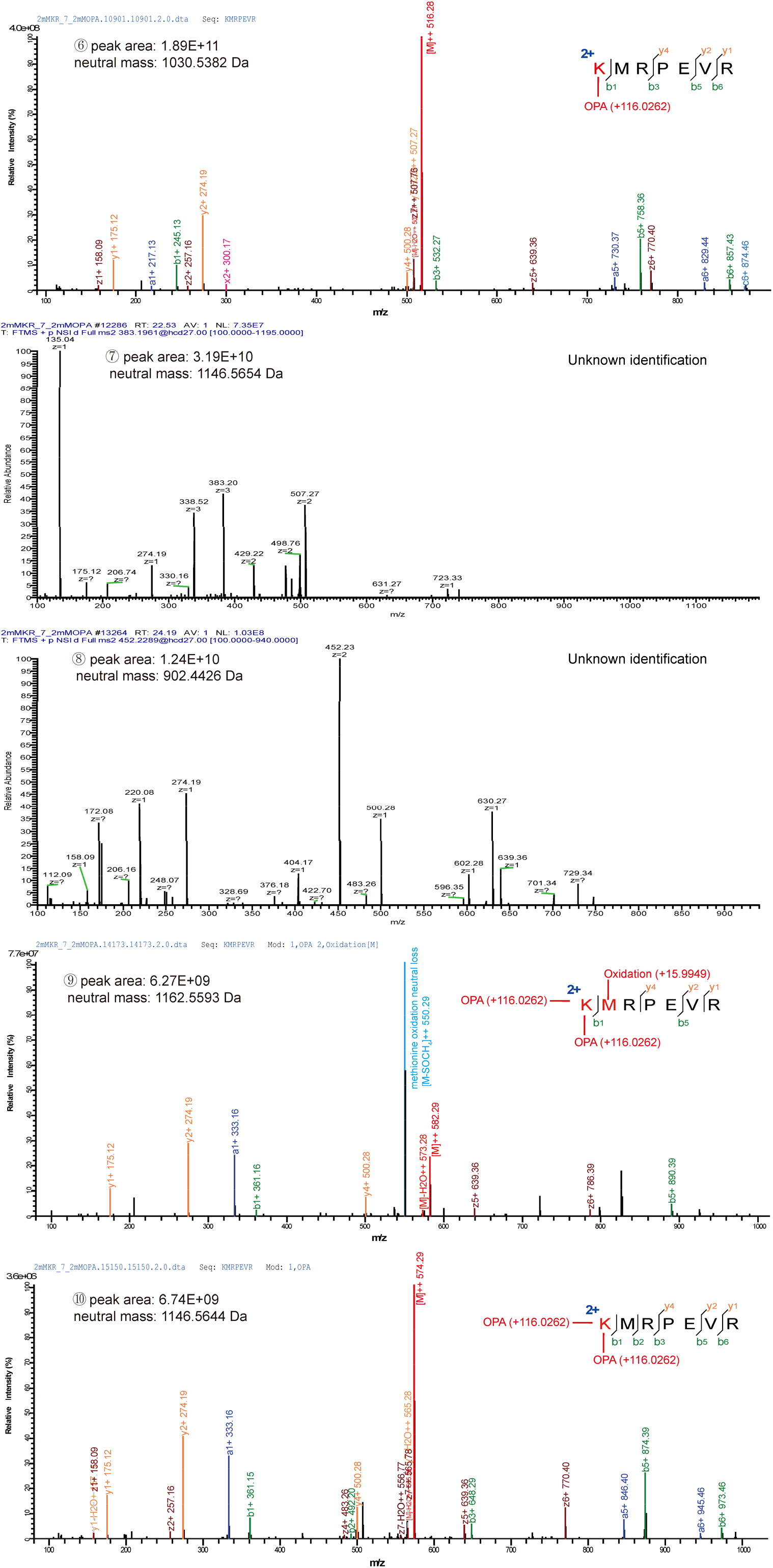
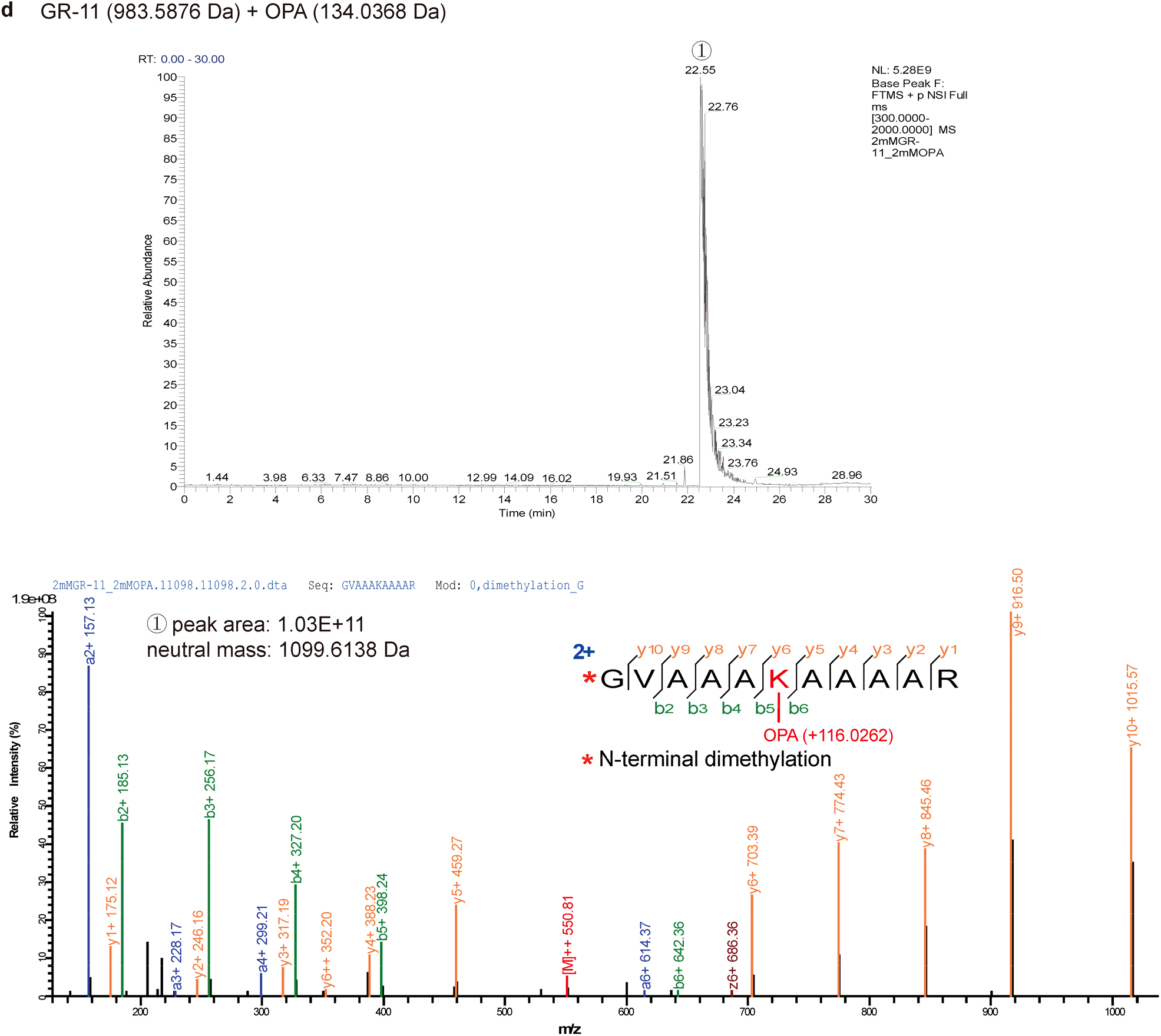
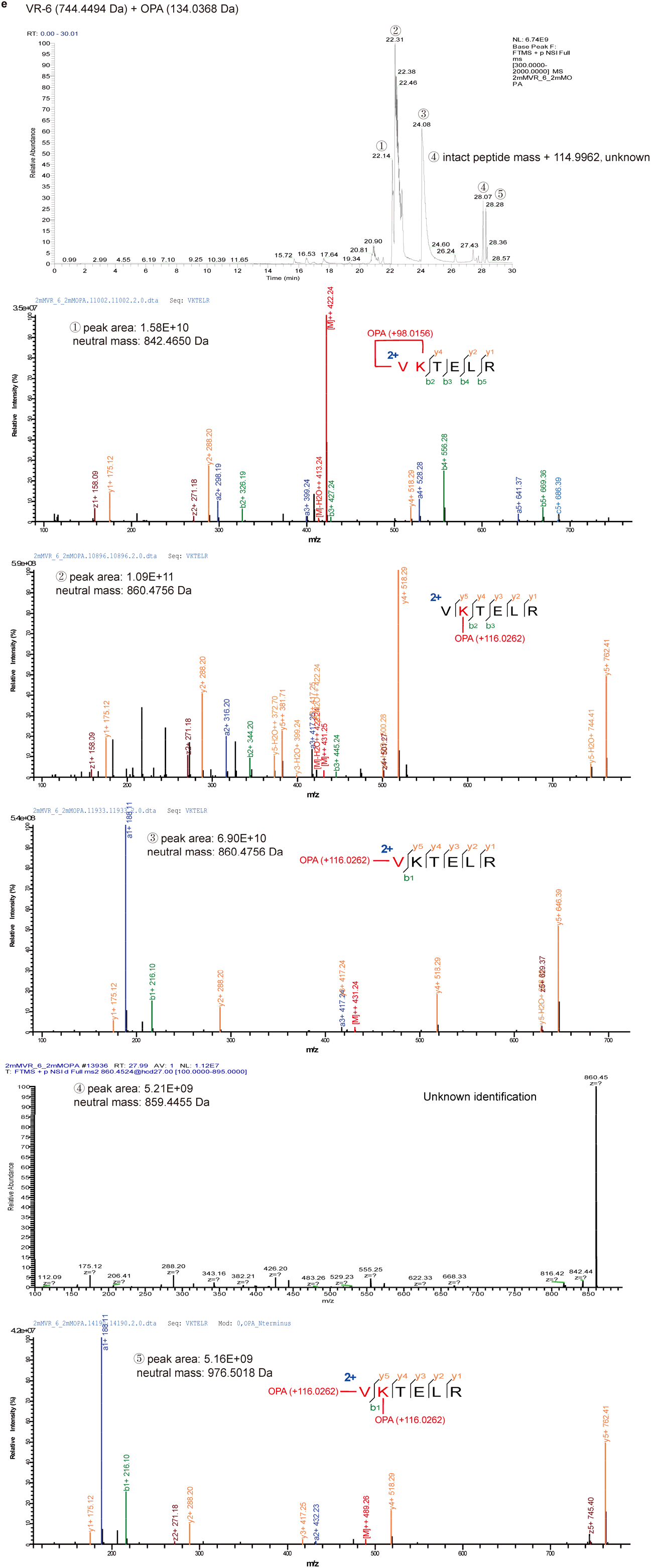
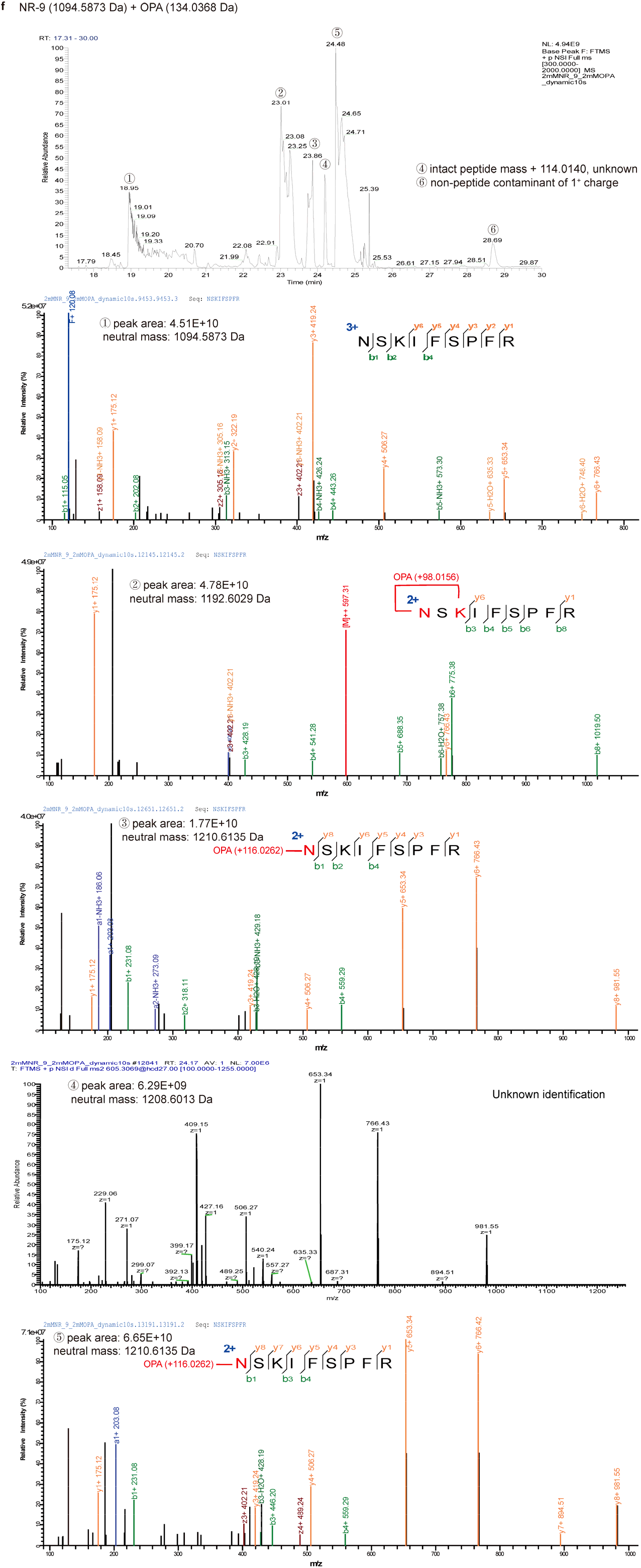
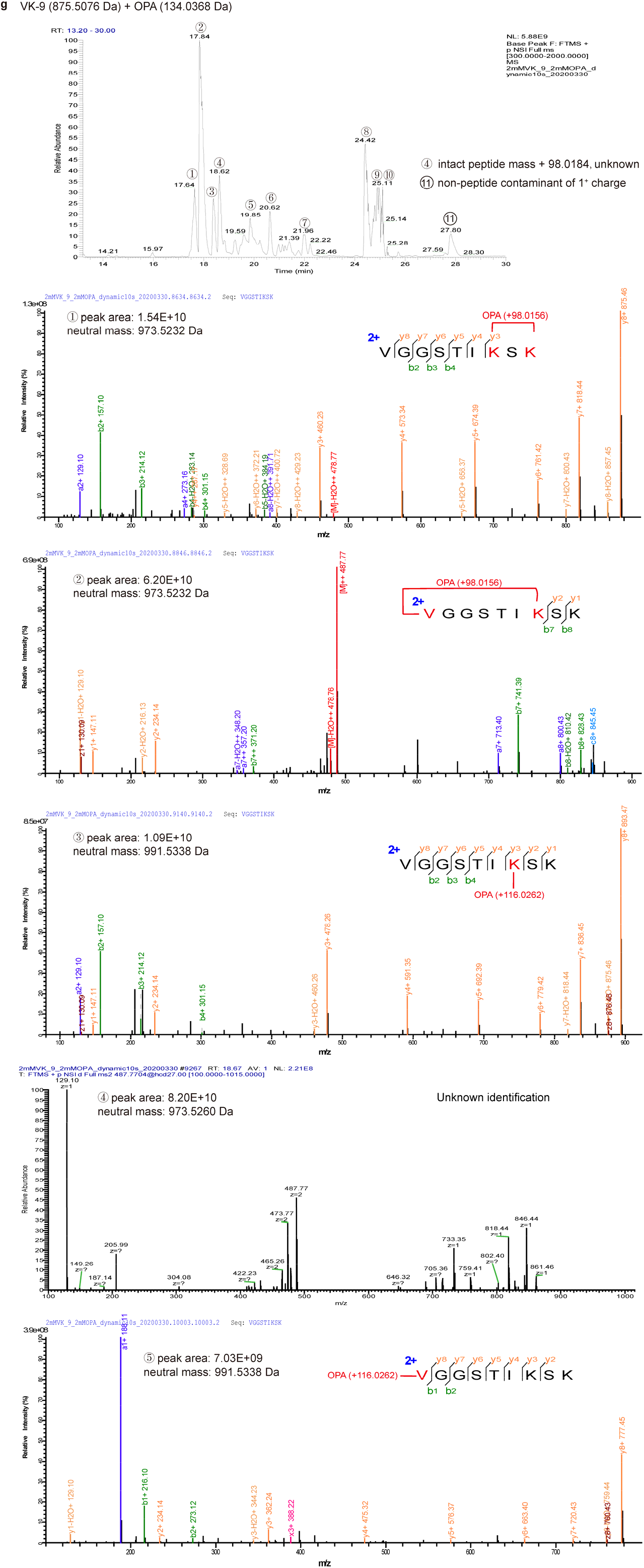
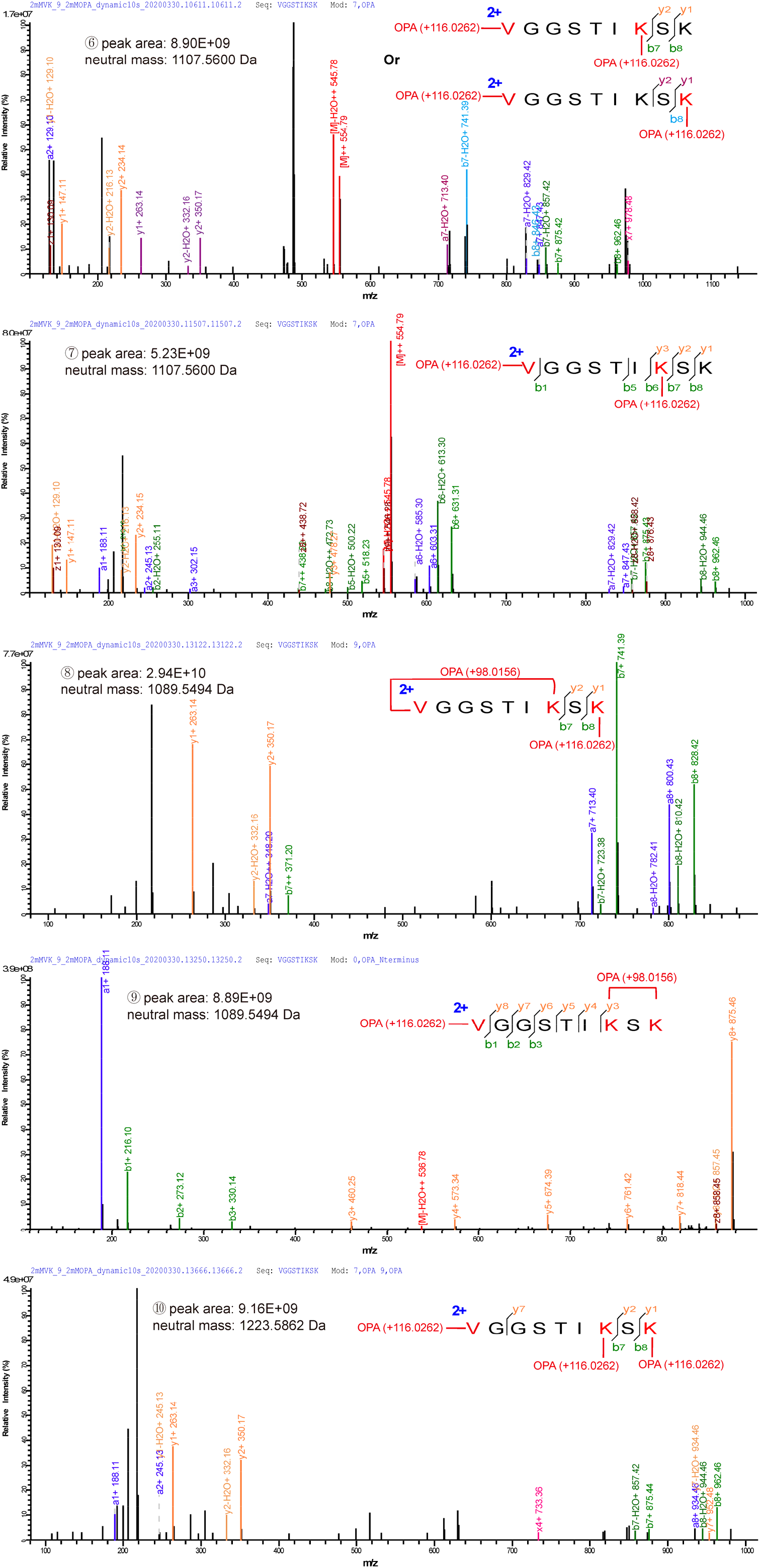
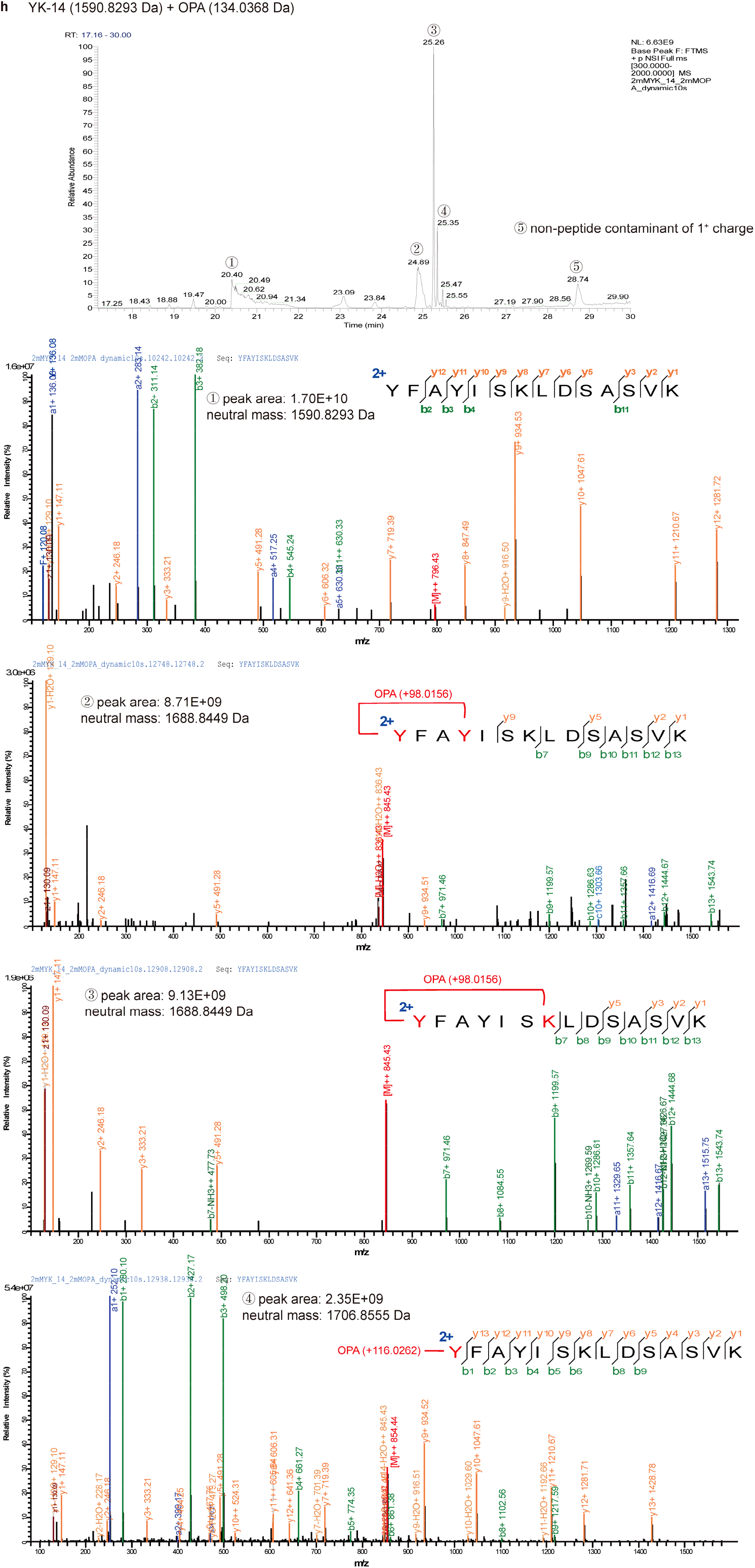
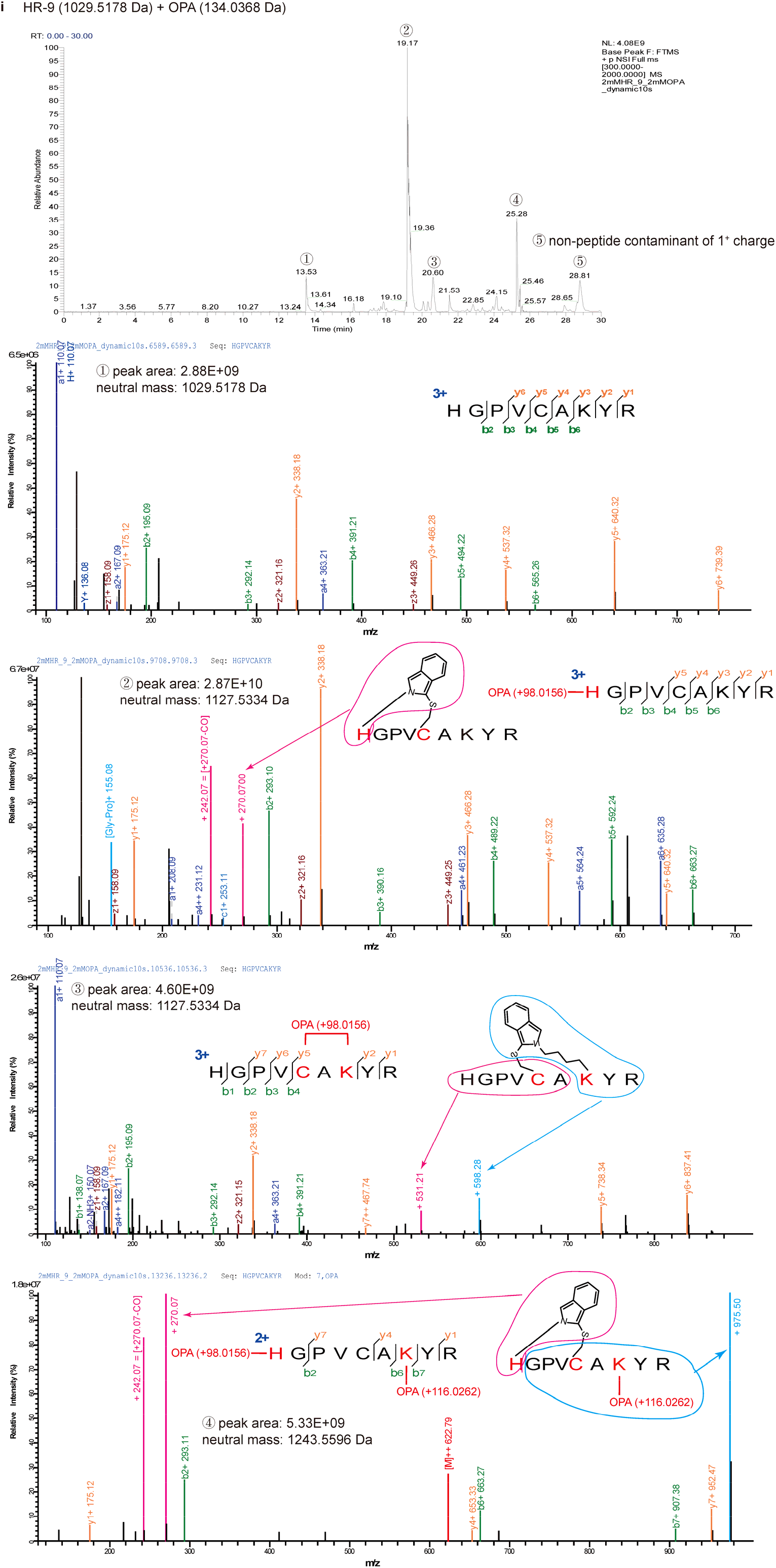
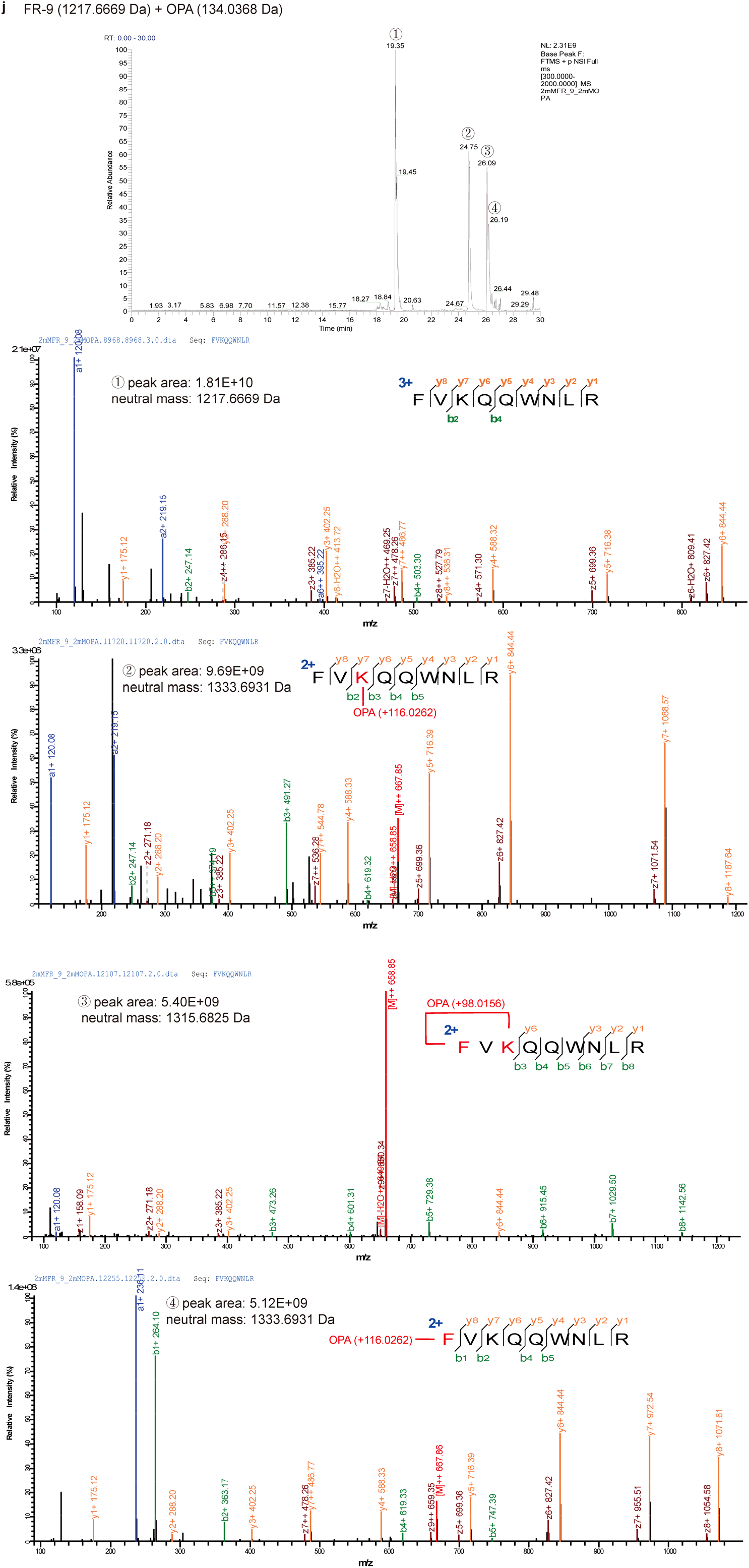
Annotating the reaction products of ten synthetic peptides and OPA in the selectivity test. (a - j) MS1 chromatographic peak and the corresponding HCD spectrum of every peak. Lysine ε-NH_2_ or peptide N-terminal α-NH_2_ reacted with OPA, forming N-substituted phthalimidines (product 1, Δ mass = + 116.0262 Da). The loop-linked side product (product 2, Δ mass = +98.0156 Da) resulted from OPA jointing an amino group and another nucleophilic group on the same peptide. The modified residues are indicated in red. (Note: peptide GR-11 (d) has no loop-linked side products due to the dimethylation of its N-terminus; the sixth peak in peptide VK-9 (g) was a mixture of two products, as labelled in MS2 spectrum; the second and fourth peaks in peptide HR-9 (i) were presumed to contain the loop-linked side products between N terminal α-NH_2_ and a sulfhydryl group from cysteine, but some fragment ions support the addition of 98.0156 Da at the N terminus without clear explanation, as labelled in MS2 spectrum.)

**Supplementary Figure 3.**
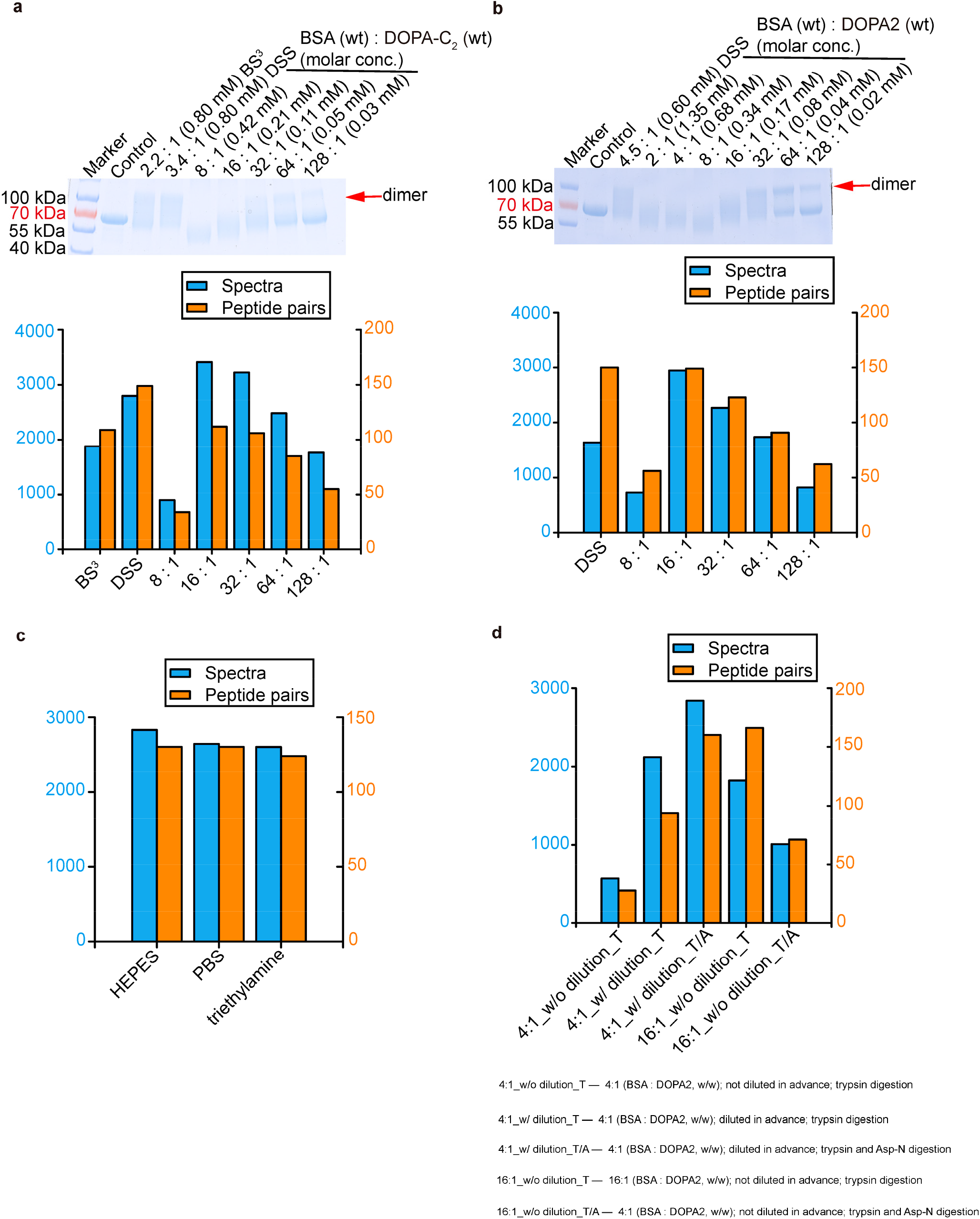
Optimization of cross-linking conditions using BSA. (a and b) Optimizing the working concentration of cross-linkers DOPA-C_2_ and DOPA2. (c) Performance of DOPA2 in HEPES buffer, PBS buffer and triethylamine buffer on BSA. (d) Comparing cross-linking effects of DOPA2 with or without pre-dilution to 2× the final concentration, followed by digestion with trypsin alone or with trypsin plus Asp-N. Identified cross-links were filtered by requiring FDR < 0.01 at the spectral level.

**Supplementary Figure 4.**
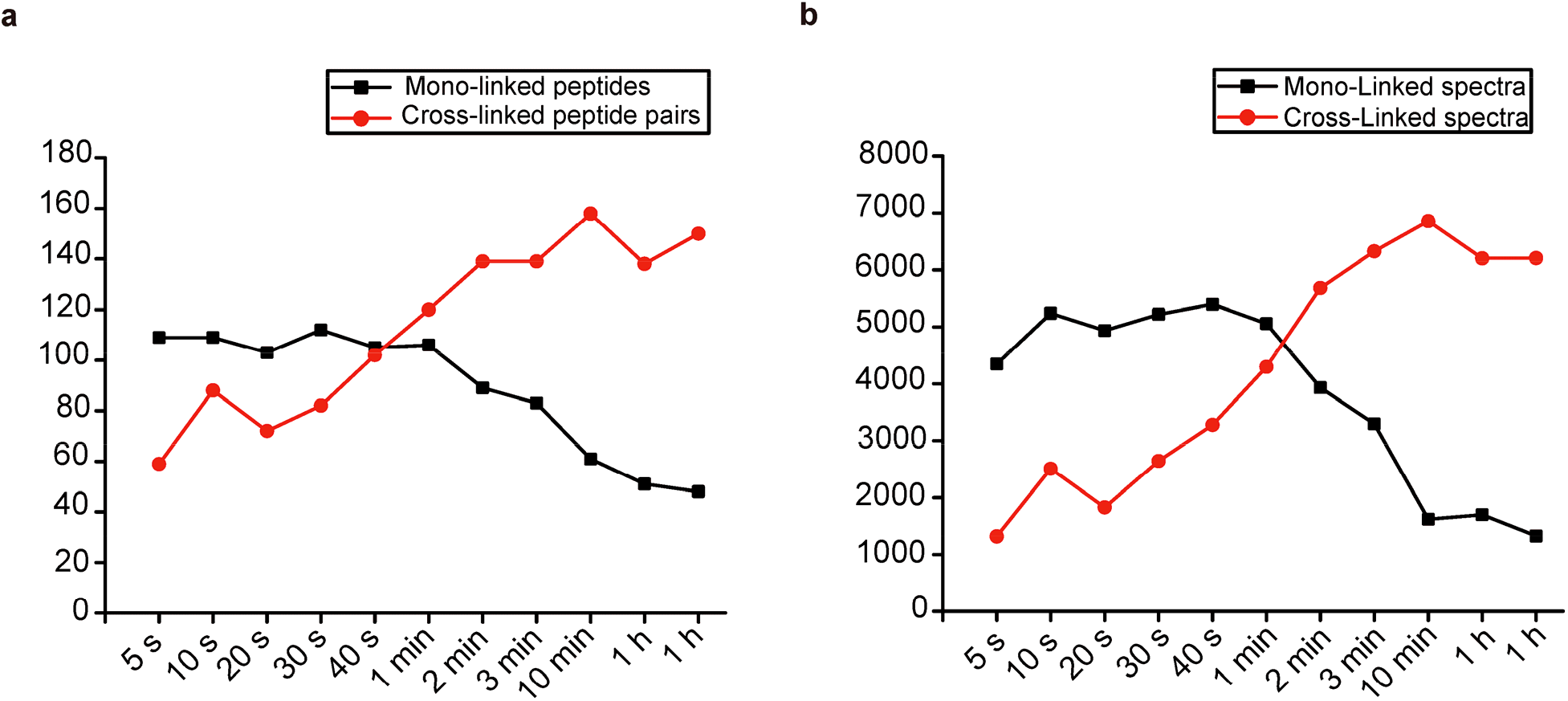
The distribution of mono-links and cross-links on BSA with the increasing reaction time. (a) The number of mono-linked peptides and cross-linked peptide pairs with the increasing reaction time. (b) The number of mono-linked spectra and cross-linked spectra with the increasing reaction time.

**Supplementary Figure 5.**
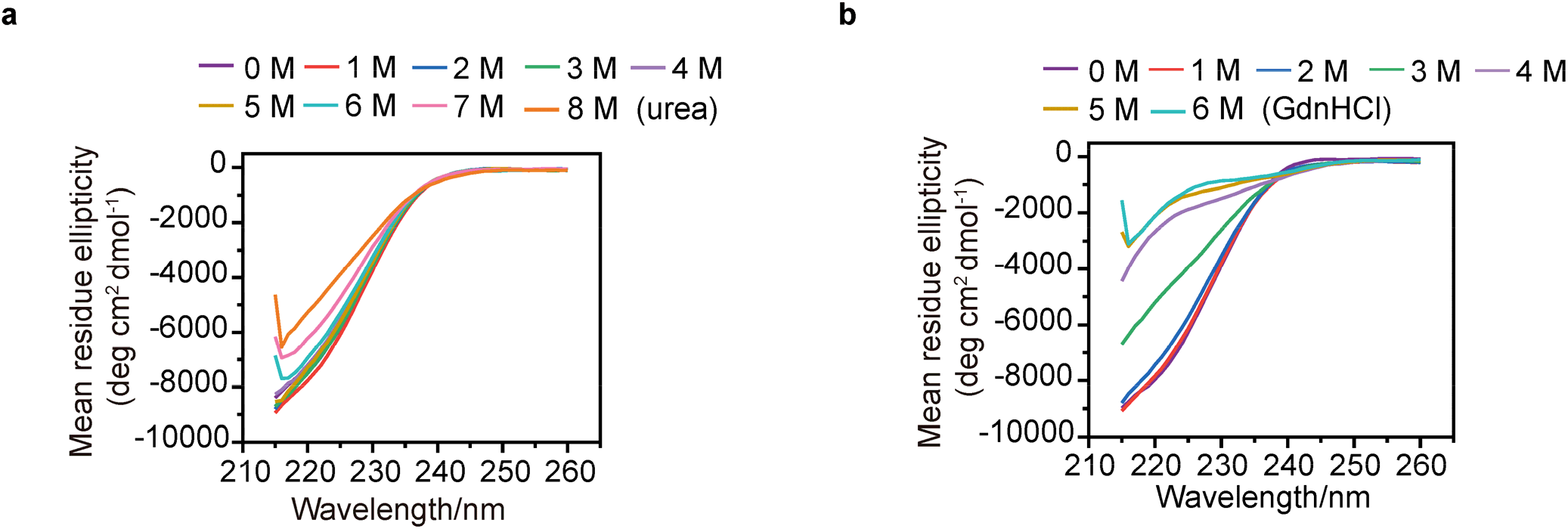
Circular dichroism spectra in far-UV region measured for RNase A in the presence of denaturants. (a) RNase A unfolding at different urea concentrations. (b) RNase A unfolding at different GdnHCl concentrations.

**Supplementary Figure 6.**
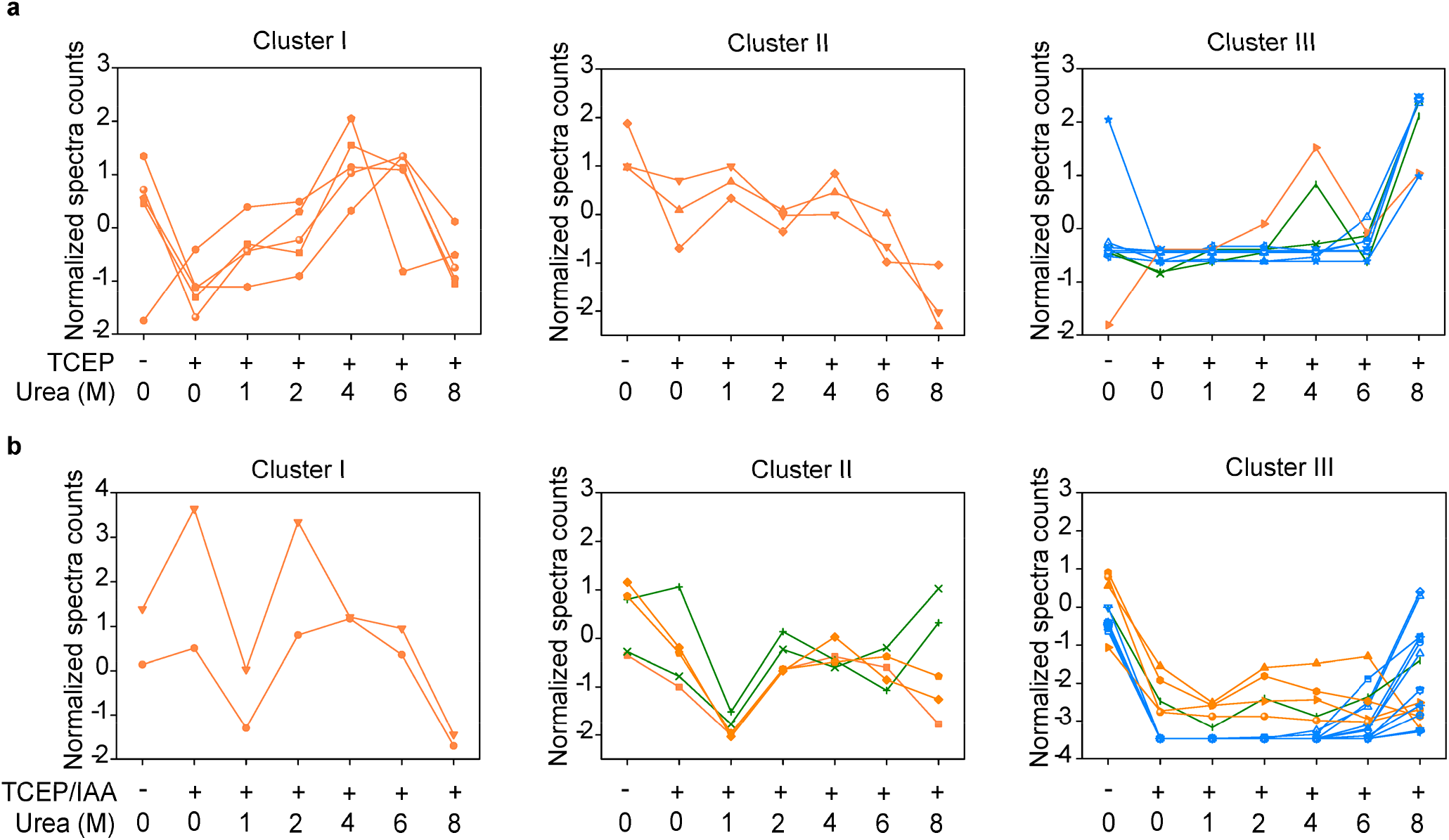
Studying the unfolding process of RNase A in urea buffer using DOPA2 under reduction conditions. (a) The changes of spectral counts for each identified cross-linked residue pair in different concentrations of urea. And RNase A samples are reduced by TCEP ahead of denaturation in urea buffer. The residue pairs identified were classified into three clusters by K-means (Cluster I, Cluster II, cluster III). The coloring of cross-links is consistent with Cluster A, Cluster B and Cluster C in Figure 3b. (b) Similar to a, but RNase A samples are respectively reduced and blocked by TCEP and IAA ahead of denaturation in urea buffer. Cross-linking residue pairs were filtered by requiring FDR < 0.01 at the spectra level, E-value < 1×10^−8^ and spectral counts > 3.

**Supplementary Figure 7.**
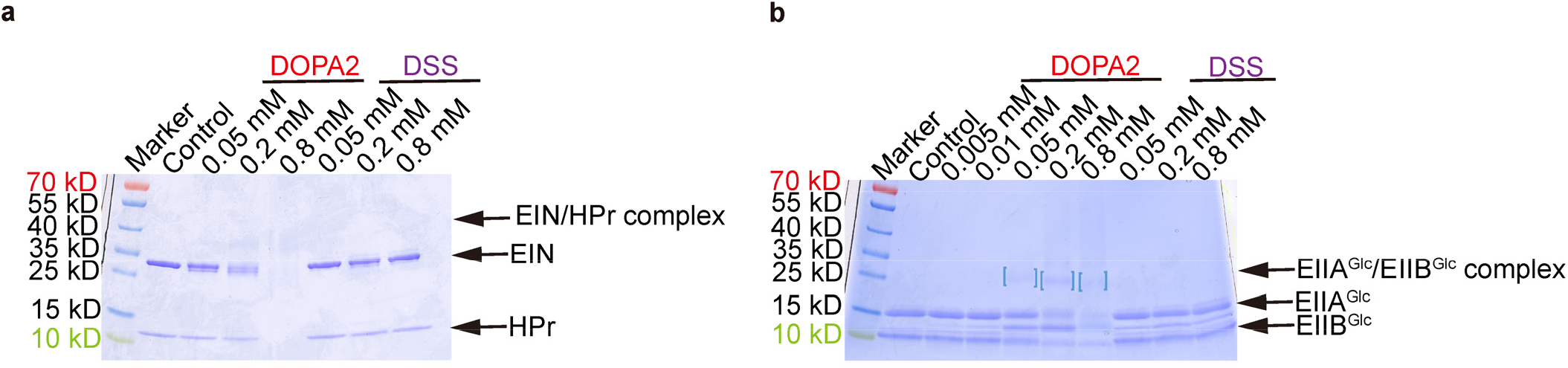
Performance of DOPA2 and DSS on weak protein interactions at the protein concentration of 1× *K*_D_. (a) SDS-PAGE of DOPA2 or DSS-cross-linked EIN/HPr. The concentration of EIN/HPr is 1 × *K*_D_. (b) As in a, but for the EIIA^Glc^/EIIB^Glc^ complex. Dimers were marked by square brackets.

**Supplementary Figure 8.**
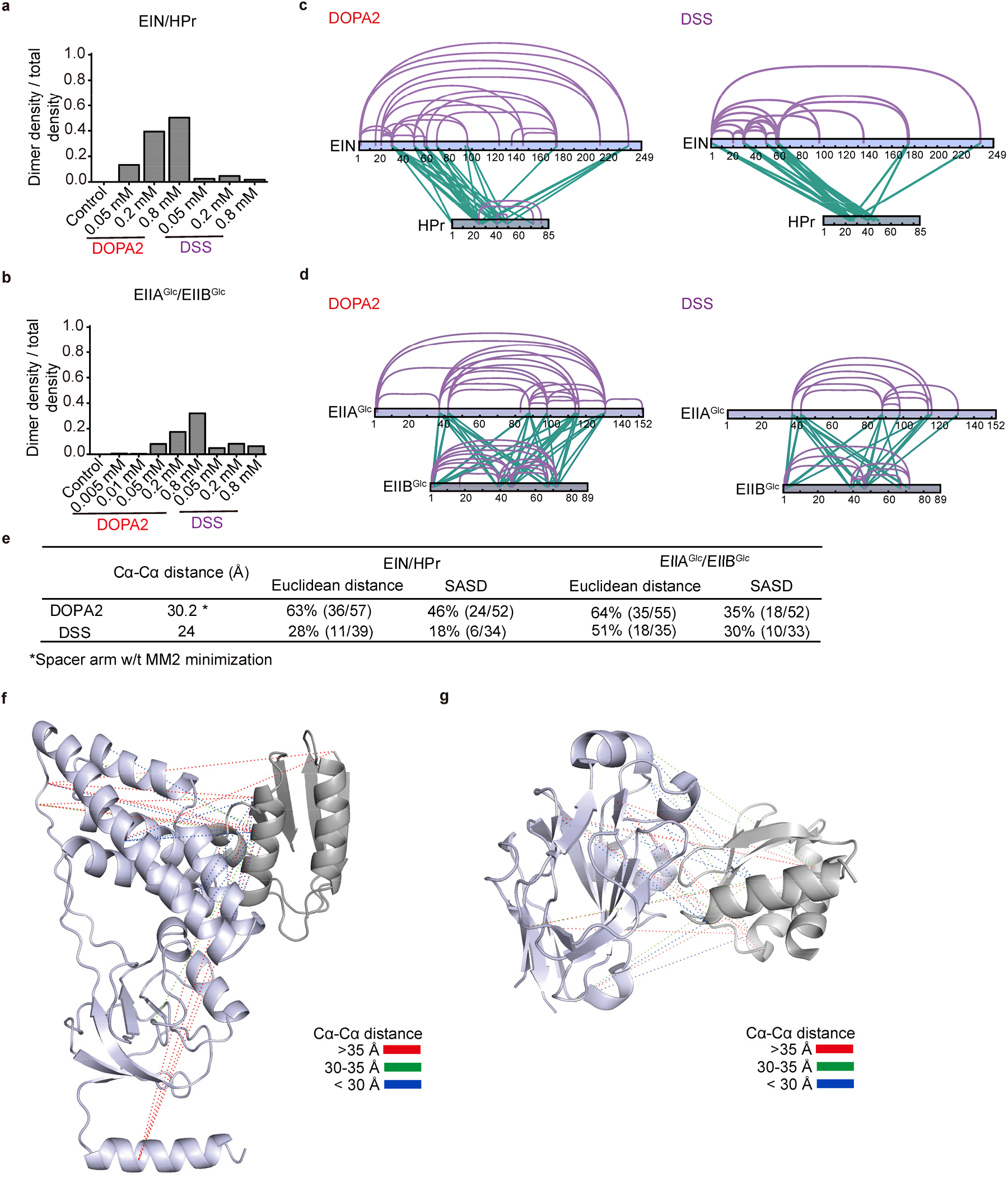
Performance of DOPA2 and DSS on weak protein interactions at the protein concentration of 10× *K*_D_. (a) The quantitative data of SDS-PAGE of DOPA2 or DSS-cross-linked EIN/HPr in Figure 4b. The ratio of dimer grey density to total grey density of each sample was shown in a. (b) As in a, but for the EIIA^Glc^/EIIB^Glc^ in Figure 4d. The ratio of dimer grey density to total grey density of each sample was shown in b. (c) DOPA2-cross-linked or DSS-cross-linked residue pairs identified from excised dimer bands that were mapped to the primary sequences of EIN/HPr complex subunits (visualized using xiNET^61^). (d) As in d, but for the EIIA^Glc^/EIIB^Glc^ complex. (e) The table displays the percentage of residue pairs (inter-links plus intra-links) that are consistent with the structures of weak protein complexes, calculated by the use of the Euclidean distance or the solvent accessible surface distance; the table also displays the maximum distance restraints imposed by each linker. (f and g) Identified inter-molecular DOPA2 cross-links mapped onto the crystal structures of EIN/HPr (PDB code: 3EZA) or EIIA^Glc^/EIIB^Glc^ (PDB code: 1O2F). The Cα-Cα Euclidean distances of cross-links are color-coded. All the cross-linking residue pairs were filtered by requiring an FDR < 0.01 at the spectra level, E-value < 1×10^−3^, and spectral counts > 3.

**Supplementary Table 1.**
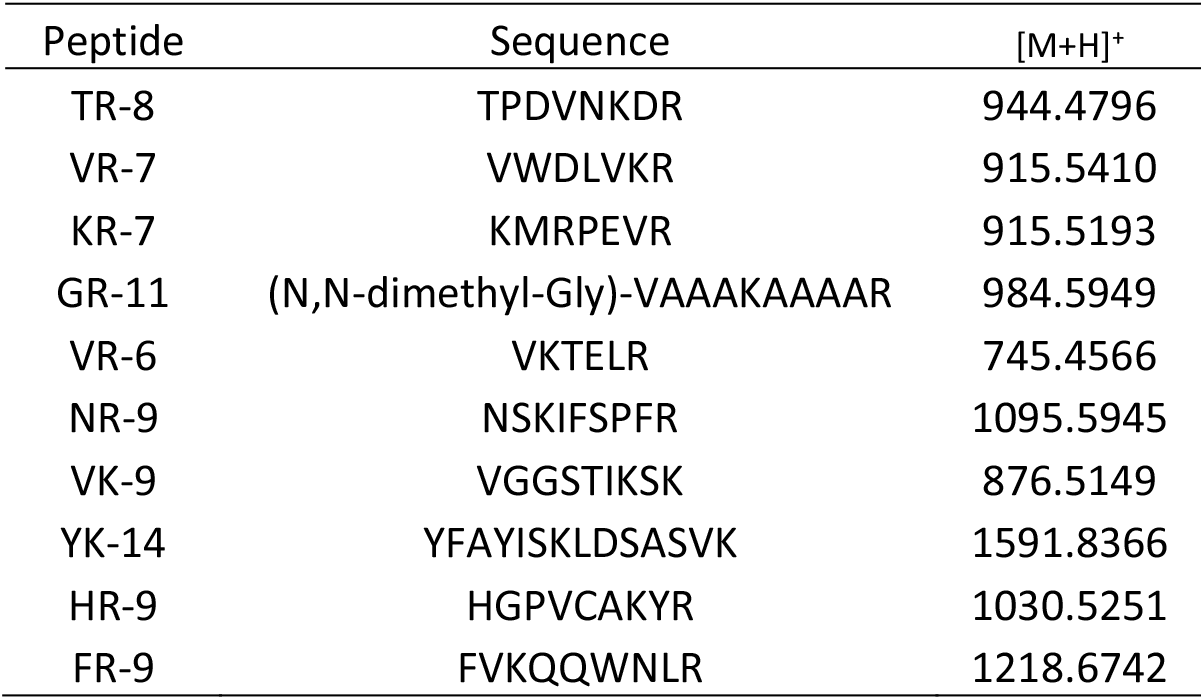
Peptides tested for OPA selectivity.

**Supplementary Table 2.**
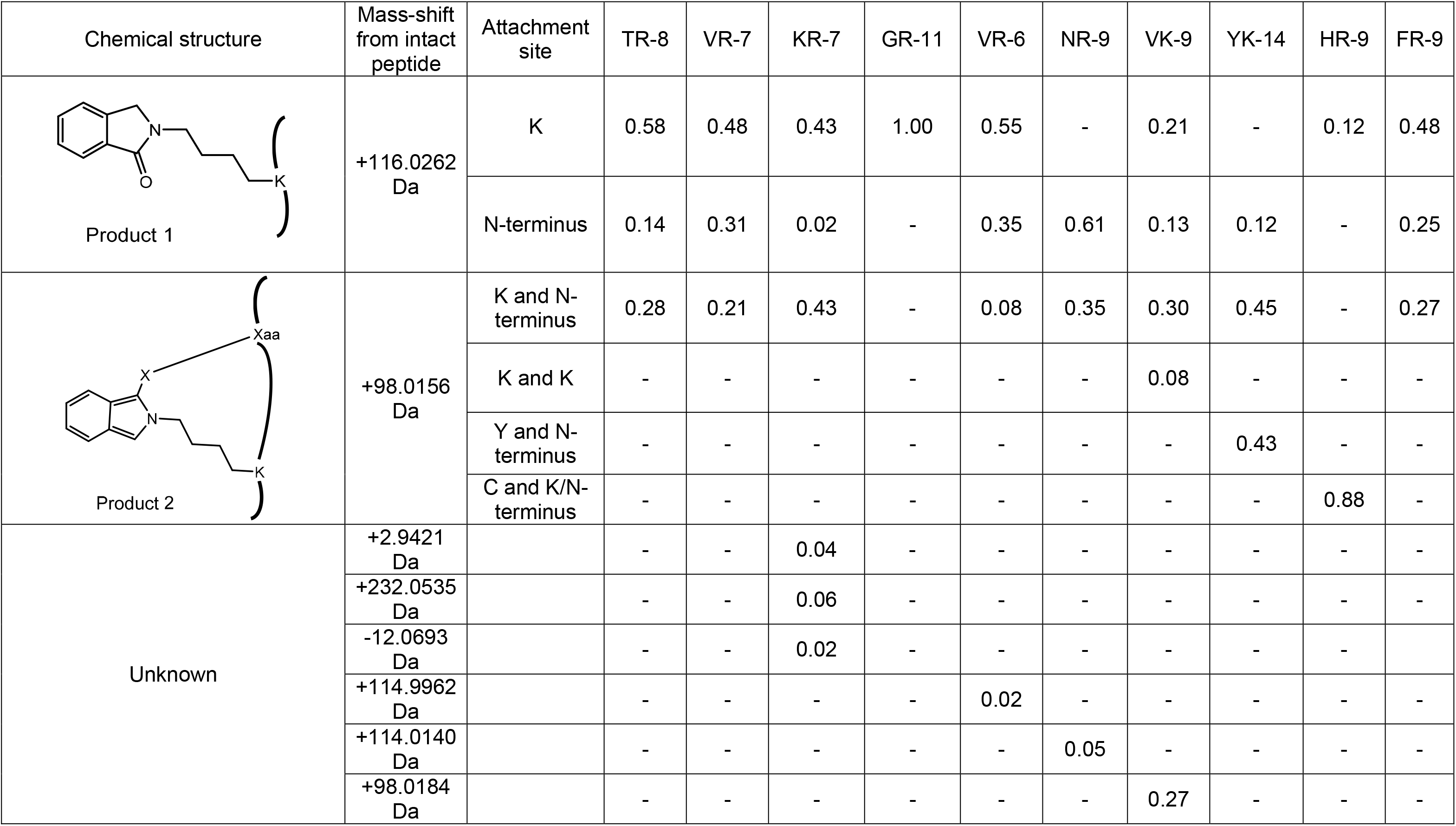
Products of ten synthesized peptides and OPA. Peptides are modified after reacting. For each product, the mass-shift from the intact peptide is indicated, and the relative abundance is calculated based on the chromatographic peak areas from the corresponding. (X = N, S, or O; Xaa = Lys, Cys, Tyr, or the peptide N-terminus)

| Chemical structure | Mass-shift from intact peptide | Attachment site | TR-8 | VR-7 | KR-7 | GR-11 | VR-6 | NR-9 | VK-9 | YK-14 | HR-9 | FR-9 |
| --- | --- | --- | --- | --- | --- | --- | --- | --- | --- | --- | --- | --- |
| Product 1 | +116.0262 Da | K | 0.58 | 0.48 | 0.43 | 1.00 | 0.55 | - | 0.21 | - | 0.12 | 0.48 |
|  |  | N-terminus | 0.14 | 0.31 | 0.02 | - | 0.35 | 0.61 | 0.13 | 0.12 | - | 0.25 |
| Product 2 | +98.0156 Da | K and N-terminus | 0.28 | 0.21 | 0.43 | - | 0.08 | 0.35 | 0.30 | 0.45 | - | 0.27 |
|  |  | K and K | - | - | - | - | - | - | 0.08 | - | - | - |
|  |  | Y and N-terminus | - | - | - | - | - | - | - | 0.43 | - | - |
|  |  | C and K/N-terminus | - | - | - | - | - | - | - | - | 0.88 | - |
| Unknown | +2.9421 Da |  | - | - | 0.04 | - | - | - | - | - | - | - |
|  | +232.0535 Da |  | - | - | 0.06 | - | - | - | - | - | - | - |
|  | -12.0693 Da |  | - | - | 0.02 | - | - | - | - | - | - | - |
|  | +114.9962 Da |  | - | - | - | - | 0.02 | - | - | - | - | - |
|  | +114.0140 Da |  | - | - | - | - | - | 0.05 | - | - | - | - |
|  | +98.0184 Da |  | - | - | - | - | - | - | 0.27 | - | - | - |

**Supplementary Table 3.** The composition of a ten-protein mixture and the molecular weight for each protein.

| # | Protein Name | MW (kDa) |
| --- | --- | --- |
| 1 | myosin | 242 |
| 2 | lactoferrin | 78 |
| 3 | BSA | 69 |
| 4 | catalase | 60 |
| 5 | β-amylase | 56 |
| 6 | aldolase | 39 |
| 7 | carbonic anhydrase 2 | 29 |
| 8 | GST | 28 |
| 9 | PUD-1/PUD-2 heterodimer | 17/17 |
| 10 | lysozyme | 16 |

**Supplementary Table 4.** The intensity of different products in peptide samples cross-linked by DOPA2 or DSS at low pH and in the presence of denaturants. (a) The chromatographic base peak intensity of different products in peptide samples cross-linked by DOPA2 in low pH value buffer and physiology buffer. “Regular peptide” refers to free peptides without cross-linking; “X-link” refers to the situation wherein two peptides are linked with one molecule of DOPA2; “Mono-link” refers to a peptide that has been modified but is not cross-linked by a cross-linker. (b) The chromatographic base peak intensity of different products in peptide samples cross-linked by DSS in low pH value buffer and physiology buffer. (c) The chromatographic base peak intensity of different products in peptide samples cross-linked by DOPA2 in 6 M GdnHCl buffer or HEPES buffer.

| Peptide sequence | DOPA2 |  |  |  |  |  |  |  |
| --- | --- | --- | --- | --- | --- | --- | --- | --- |
|  | HEPES Buffer, pH7.4 |  |  |  | Citric Acid - Na<sub>2</sub>HPO<sub>4</sub> Buffer, pH3.0 |  |  |  |
|  | Regular peptide (A) | X-link (B) | Mono-link (C) | (B+C)/(A+B+C) | Regular peptide (A) | X-link (B) | Mono-link (C) | (B+C)/(A+B+C) |
| TR-8 | 5.34E+07 | 2.36E+06 | 7.19E+07 | 0.58156 | 1.07E+08 | 1.56E+06 | 5.45E+07 | 0.34478 |
| VR-7 | 8.33E+09 | 2.63E+06 | 2.24E+08 | 0.02647 | 3.72E+09 | 9.15E+06 | 9.88E+08 | 0.21130 |
| KR-7 | 2.36E+08 | 7.38E+08 | 7.38E+08 | 0.86227 | 1.91E+06 | 0.00E+00 | 3.57E+08 | 0.99469 |
| GR-11 | 8.40E+09 | 1.52E+09 | 1.52E+09 | 0.26542 | 1.37E+10 | 4.02E+06 | 3.68E+09 | 0.21221 |

b
| Peptide sequence | DSS |  |  |  |  |  |  |  |
| --- | --- | --- | --- | --- | --- | --- | --- | --- |
|  | HEPES Buffer, pH7.4 |  |  |  | Citric Acid - Na<sub>2</sub>HPO<sub>4</sub> Buffer, pH3.0 |  |  |  |
|  | Regular peptide (A) | X-link (B) | Mono-link (C) | (B+C)/(A+B+C) | Regular peptide (A) | X-link (B) | Mono-link (C) | (B+C)/(A+B+C) |
| TR-8 | 1.12E+08 | 0 | 6.00E+07 | 0.34863 | 8.77E+08 | 0 | 3.33E+04 | 0.00004 |
| VR-7 | 6.16E+09 | 0 | 5.32E+08 | 0.07951 | 1.11E+10 | 0 | 3.97E+07 | 0.00357 |
| KR-7 | 1.03E+08 | 0 | 5.07E+08 | 0.83055 | 9.69E+07 | 0 | 2.23E+06 | 0.02245 |
| GR-11 | 9.89E+09 | 0 | 1.53E+06 | 0.00015 | 1.53E+10 | 0 | 1.89E+06 | 0.00012 |

c
| Peptide sequence | DOPA2 |  |  |  |  |  |  |  |
| --- | --- | --- | --- | --- | --- | --- | --- | --- |
|  | HEPES Buffer, pH7.4 |  |  |  | 6 M GdnHCl |  |  |  |
|  | Regular peptide (A) | X-link (B) | Mono-link (C) | (B+C)/(A+B+C) | Regular peptide (A) | X-link (B) | Mono-link (C) | (B+C)/(A+B+C) |
| VR-7 | 1.71E+10 | 8.27E+08 | 6.37E+09 | 0.29607 | 2.37E+10 | 2.56E+09 | 2.77E+10 | 0.56072 |
| GR-11 | 3.01E+08 | 1.55E+10 | 1.00E+10 | 0.98834 | 7.47E+08 | 3.01E+08 | 1.51E+10 | 0.95381 |

**Supplementary Table 5.** The cross-links identified from RNase A by DOPA2. (a) The identified cross-links from RNase A in various concentrations of urea using DOPA2, including linked amino acid positions, the Cα-Cα distance of each cross-link on crystal structures and the number of spectra seen in different concentrations of urea. (b) As in a, but for identified cross-links from RNase A in various concentrations of GdnHCl. (c) Similar to a, but for the identified cross-links from RNase A exposed to 8 M urea for varying amounts of time using DOPA2. Cross-linking residue pairs were filtered by requiring FDR < 0.01 at the spectra level, E-value < 1×10^−8^ and spectral counts > 3. The estimated FDR at the residue pair level is zero.

| | Linked aa positions | Cα-Cα distance (Å)<br>(PDB code: 6ETK) | # of spectra seen in various conc. of urea | | | | | |
| --- | --- | --- | --- | --- | --- | --- | --- | --- |
|  |  |  | 0 M urea | 1 M urea | 2 M urea | 4 M urea | 6 M urea | 8 M urea |
| Cluster A | RNase_A(1)-RNase_A(37) | 17.0 | 206 | 220 | 196 | 230 | 183 | 88 |
|  | RNase_A(7)-RNase_A(37) | 14.5 | 189 | 184 | 177 | 182 | 139 | 82 |
|  | RNase_A(1)-RNase_A(41) | 19.8 | 152 | 198 | 161 | 195 | 125 | 43 |
|  | RNase_A(7)-RNase_A(41) | 13.8 | 480 | 444 | 396 | 356 | 218 | 61 |
|  | RNase_A(66)-RNase_A(98) | 23.3 | 35 | 27 | 49 | 25 | 43 | 11 |
|  | RNase_A(41)-RNase_A(66) | 18.6 | 129 | 125 | 101 | 100 | 83 | 61 |
|  | RNase_A(41)-RNase_A(104) | 21.6 | 132 | 101 | 108 | 91 | 76 | 28 |
|  | RNase_A(66)-RNase_A(104) | 17.0 | 148 | 150 | 124 | 206 | 140 | 34 |
|  | RNase_A(7)-RNase_A(66) | 19.3 | 28 | 18 | 12 | 14 | 19 | 1 |
|  | RNase_A(61)-RNase_A(104) | 14.6 | 19 | 25 | 8 | 6 | 8 | 4 |
|  | RNase_A(31)-RNase_A(37) | 7.2 | 210 | 180 | 183 | 178 | 150 | 147 |
| Cluster B | RNase_A(41)-RNase_A(91) | 10.9 | 0 | 0 | 0 | 21 | 140 | 264 |
|  | RNase_A(37)-RNase_A(104) | 28.2 | 3 | 0 | 0 | 2 | 30 | 80 |
|  | RNase_A(37)-RNase_A(98) | 18.0 | 13 | 9 | 16 | 69 | 149 | 286 |
|  | RNase_A(1)-RNase_A(98) | 30.5 | 0 | 0 | 0 | 0 | 1 | 36 |
|  | RNase_A(1)-RNase_A(91) | 28.2 | 0 | 0 | 0 | 1 | 39 | 82 |
|  | RNase_A(7)-RNase_A(98) | 22.3 | 0 | 0 | 0 | 0 | 4 | 31 |
|  | RNase_A(41)-RNase_A(98) | 12.6 | 0 | 0 | 0 | 7 | 102 | 254 |
|  | RNase_A(31)-RNase_A(98) | 13.5 | 0 | 0 | 0 | 1 | 11 | 50 |
|  | RNase_A(66)-RNase_A(91) | 27.7 | 0 | 0 | 0 | 0 | 10 | 50 |
|  | RNase_A(1)-RNase_A(31) | 20.3 | 1 | 0 | 1 | 0 | 4 | 22 |
|  | RNase_A(7)-RNase_A(91) | 23.9 | 0 | 0 | 0 | 8 | 50 | 114 |
|  | RNase_A(7)-RNase_A(31) | 14.8 | 2 | 0 | 2 | 1 | 4 | 35 |
|  | RNase_A(61)-RNase_A(91) | 38.2 | 0 | 0 | 0 | 0 | 4 | 29 |
|  | RNase_A(91)-RNase_A(104) | 29.0 | 0 | 0 | 0 | 1 | 21 | 86 |
|  | RNase_A(31)-RNase_A(41) | 10.0 | 0 | 0 | 0 | 0 | 1 | 57 |
|  | RNase_A(61)-RNase_A(98) | 30.0 | 0 | 0 | 0 | 0 | 0 | 27 |
|  | RNase_A(1)-RNase_A(61) | 30.3 | 0 | 0 | 0 | 0 | 0 | 7 |
| Cluster C | RNase_A(37)-RNase_A(91) | 12.3 | 162 | 128 | 129 | 156 | 174 | 218 |
|  | RNase_A(37)-RNase_A(41) | 9.0 | 320 | 273 | 246 | 234 | 246 | 329 |
|  | RNase_A(31)-RNase_A(91) | 13.3 | 102 | 96 | 91 | 101 | 103 | 162 |

b
|  | Linked aa positions | Ca-Ca distance (Å)<br>(PDB code: 6ETK ) | # of spectra seen in various conc. of GdnHCl |  |  |  |  |  |  |
| --- | --- | --- | --- | --- | --- | --- | --- | --- | --- |
|  |  |  | 0 M<br>GdnHCl | 1 M<br>GdnHCl | 2 M<br>GdnHCl | 3 M<br>GdnHCl | 4 M<br>GdnHCl | 5 M<br>GdnHCl | 6 M<br>GdnHCl |
| Cluster A | RNase_A(37)-RNase_A(41) | 9.0 | 154 | 68 | 34 | 56 | 43 | 51 | 41 |
|  | RNase_A(1)-RNase_A(41) | 19.8 | 55 | 19 | 16 | 12 | 0 | 0 | 0 |
|  | RNase_A(66)-RNase_A(98) | 23.3 | 67 | 9 | 1 | 7 | 2 | 2 | 0 |
|  | RNase_A(41)-RNase_A(66) | 18.6 | 43 | 7 | 0 | 2 | 4 | 4 | 0 |
|  | RNase_A(41)-RNase_A(104) | 21.6 | 118 | 59 | 29 | 24 | 6 | 0 | 0 |
|  | RNase_A(7)-RNase_A(41) | 13.8 | 284 | 215 | 125 | 45 | 1 | 0 | 0 |
|  | RNase_A(7)-RNase_A(37) | 14.5 | 168 | 150 | 109 | 49 | 39 | 30 | 28 |
|  | RNase_A(1)-RNase_A(37) | 17.0 | 80 | 65 | 60 | 42 | 19 | 20 | 16 |
|  | RNase_A(66)-RNase_A(104) | 17.0 | 87 | 62 | 52 | 21 | 0 | 0 | 0 |
|  | RNase_A(7)-RNase_A(66) | 19.3 | 54 | 32 | 12 | 1 | 0 | 0 | 0 |
|  | RNase_A(61)-RNase_A(104) | 14.6 | 41 | 48 | 25 | 12 | 1 | 0 | 0 |
| Cluster B | RNase_A(37)-RNase_A(98) | 18.0 | 6 | 18 | 28 | 105 | 106 | 86 | 93 |
|  | RNase_A(1)-RNase_A(91) | 28.2 | 0 | 1 | 0 | 35 | 47 | 41 | 29 |
|  | RNase_A(41)-RNase_A(98) | 12.6 | 0 | 0 | 0 | 66 | 64 | 67 | 56 |
|  | RNase_A(1)-RNase_A(98) | 30.5 | 0 | 0 | 0 | 16 | 15 | 17 | 10 |
|  | RNase_A(61)-RNase_A(91) | 38.2 | 0 | 0 | 2 | 55 | 44 | 43 | 33 |
|  | RNase_A(61)-RNase_A(98) | 30.0 | 0 | 0 | 0 | 30 | 4 | 10 | 0 |
|  | RNase_A(41)-RNase_A(91) | 10.9 | 0 | 0 | 1 | 91 | 82 | 69 | 61 |
|  | RNase_A(7)-RNase_A(91) | 23.9 | 0 | 0 | 0 | 15 | 12 | 9 | 0 |
|  | RNase_A(91)-RNase_A(104) | 29.0 | 0 | 7 | 0 | 35 | 22 | 26 | 36 |
| Cluster C | RNase_A(37)-RNase_A(91) | 12.3 | 69 | 35 | 38 | 59 | 62 | 61 | 52 |
|  | RNase_A(31)-RNase_A(91) | 13.3 | 64 | 3 | 7 | 58 | 44 | 41 | 42 |

C
|  | Linked aa positions | Ca-Ca distance<br>(PDB code: 6ETK ) | # of spectra seen in 8 M urea at different time points |  |  |  |  |  |  |  |  |  |
| --- | --- | --- | --- | --- | --- | --- | --- | --- | --- | --- | --- | --- |
|  |  |  | 5 s | 10 s | 20 s | 30 s | 1 min | 2 min | 5 min | 10 min | 15 min | 30 min |
| Cluster 1 | RNase_A(41)-RNase_A(91) | 10.9 | 18 | 24 | 39 | 33 | 64 | 117 | 166 | 216 | 227 | 217 |
|  | RNase_A(41)-RNase_A(98) | 12.6 | 20 | 30 | 40 | 50 | 69 | 99 | 139 | 192 | 183 | 213 |
|  | RNase_A(37)-RNase_A(91) | 12.3 | 255 | 242 | 268 | 265 | 267 | 311 | 379 | 332 | 345 | 309 |
|  | RNase_A(37)-RNase_A(98) | 18.0 | 81 | 93 | 120 | 126 | 176 | 190 | 232 | 271 | 298 | 292 |
|  | RNase_A(1)-RNase_A(91) | 28.2 | 1 | 7 | 11 | 16 | 30 | 39 | 63 | 62 | 73 | 62 |
|  | RNase_A(37)-RNase_A(41) | 9.0 | 208 | 234 | 253 | 256 | 288 | 314 | 344 | 359 | 391 | 367 |
|  | RNase_A(1)-RNase_A(98) | 30.5 | 0 | 0 | 1 | 0 | 0 | 0 | 6 | 8 | 17 | 10 |
|  | RNase_A(61)-RNase_A(98) | 30.0 | 4 | 4 | 8 | 4 | 15 | 26 | 49 | 80 | 60 | 76 |
|  | RNase_A(37)-RNase_A(104) | 28.2 | 16 | 27 | 36 | 23 | 33 | 40 | 90 | 101 | 98 | 98 |
|  | RNase_A(66)-RNase_A(98) | 23.4 | 11 | 9 | 23 | 10 | 7 | 23 | 28 | 28 | 30 | 20 |
|  | RNase_A(7)-RNase_A(98) | 22.3 | 1 | 0 | 1 | 0 | 0 | 1 | 0 | 4 | 0 | 4 |
|  | RNase_A(31)-RNase_A(98) | 13.5 | 0 | 0 | 0 | 0 | 0 | 11 | 30 | 31 | 31 | 28 |
|  | RNase_A(1)-RNase_A(66) | 26.0 | 16 | 5 | 8 | 12 | 5 | 12 | 17 | 18 | 16 | 16 |
|  | RNase_A(1)-RNase_A(31) | 20.3 | 0 | 0 | 1 | 1 | 0 | 1 | 3 | 4 | 7 | 8 |
|  | RNase_A(31)-RNase_A(41) | 10.0 | 0 | 0 | 0 | 0 | 0 | 4 | 10 | 22 | 17 | 20 |
|  | RNase_A(41)-RNase_A(66) | 18.6 | 7 | 6 | 3 | 18 | 19 | 27 | 61 | 86 | 80 | 77 |
|  | RNase_A(31)-RNase_A(91) | 13.3 | 72 | 64 | 92 | 89 | 111 | 121 | 149 | 169 | 163 | 180 |
|  | RNase_A(7)-RNase_A(91) | 23.9 | 0 | 0 | 0 | 0 | 2 | 8 | 20 | 31 | 38 | 30 |
|  | RNase_A(91)-RNase_A(104) | 29.0 | 53 | 46 | 42 | 53 | 60 | 72 | 97 | 121 | 117 | 123 |
|  | RNase_A(66)-RNase_A(91) | 27.7 | 9 | 14 | 17 | 15 | 24 | 24 | 67 | 78 | 86 | 77 |
|  | RNase_A(61)-RNase_A(91) | 38.2 | 0 | 2 | 5 | 10 | 21 | 43 | 91 | 124 | 112 | 120 |
|  | RNase_A(37)-RNase_A(66) | 26.4 | 15 | 0 | 28 | 17 | 13 | 37 | 42 | 65 | 76 | 70 |
| Cluster 2 | RNase_A(7)-RNase_A(37) | 14.5 | 373 | 407 | 375 | 379 | 331 | 306 | 267 | 204 | 160 | 133 |
|  | RNase_A(1)-RNase_A(37) | 17.0 | 311 | 294 | 315 | 310 | 293 | 263 | 243 | 163 | 144 | 127 |
|  | RNase_A(7)-RNase_A(41) | 13.8 | 353 | 339 | 320 | 327 | 294 | 245 | 201 | 163 | 140 | 115 |
|  | RNase_A(1)-RNase_A(41) | 19.8 | 29 | 34 | 33 | 25 | 19 | 18 | 15 | 9 | 10 | 3 |
|  | RNase_A(66)-RNase_A(104) | 17.0 | 281 | 276 | 294 | 295 | 254 | 265 | 204 | 168 | 179 | 144 |
|  | RNase_A(7)-RNase_A(66) | 19.3 | 81 | 80 | 87 | 90 | 62 | 66 | 49 | 23 | 22 | 11 |
|  | RNase_A(31)-RNase_A(37) | 7.2 | 232 | 209 | 215 | 214 | 214 | 211 | 206 | 182 | 189 | 178 |
| Cluster 3 | RNase_A(41)-RNase_A(104) | 21.6 | 85 | 69 | 86 | 86 | 75 | 74 | 95 | 97 | 117 | 72 |
|  | RNase_A(61)-RNase_A(104) | 14.6 | 143 | 136 | 151 | 142 | 132 | 131 | 165 | 159 | 138 | 111 |
|  | RNase_A(7)-RNase_A(31) | 14.8 | 0 | 0 | 0 | 0 | 0 | 0 | 0 | 3 | 3 | 0 |

**Supplementary Table 6.**
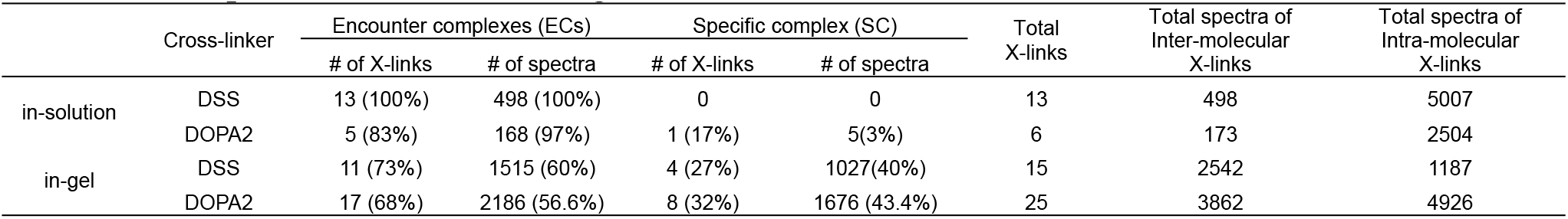
Inter-molecular cross-links identified from the EIN/HPr complex. The cross-links identification results were filtered by requiring FDR < 0.01 at the spectra level, E-value < 1×10^−8^ and spectral counts > 3. The estimated FDR at the residue pair level is zero. The cross-links were classified according to their structural compatibility with either the specific complex or the encounter complexes. The cross-linking data of 0.05, 0.2, and 0.8 mM DOPA2 or DSS were combined.

**Supplementary Table 7.** The inter-molecular cross-links identified by DOPA2 or DSS in protein complex EIN/HPr. (a) The inter-molecular cross-links identified by DSS in the solution. (b) As in a, but for cross-links identified by DOPA2. (c) The inter-molecular cross-links identified by DSS in the gel. (d) As in c, but for cross-links identified by DOPA2. Cross-links were filtered by requiring FDR < 0.01 at the spectra level, E-value < 1×10^−8^ and spectral counts > 3. The estimated FDR at the residue pair level is zero.

| Linked aa positions | Total Spec | Best E-value | Ca-Ca distance (Å)<br>(PDB code: 3EZA) | label |
| --- | --- | --- | --- | --- |
| EIN(49)-HPr(49) | 22 | 1.78E-20 | 26.48 | in-solution |
| EIN(30)-HPr(49) | 13 | 3.73E-37 | 29.90 | in-solution |
| EIN(49)-HPr(72) | 20 | 4.17E-16 | 35.22 | in-solution |
| HPr(27)-EIN(30) | 27 | 2.66E-19 | 35.30 | in-solution |
| HPr(24)-EIN(30) | 119 | 2.86E-17 | 36.78 | in-solution |
| EIN(1)-HPr(49) | 48 | 4.34E-32 | 44.96 | in-solution |
| HPr(1)-EIN(30) | 32 | 2.72E-19 | 46.92 | in-solution |
| EIN(30)-HPr(72) | 9 | 3.71E-21 | 47.21 | in-solution |
| EIN(1)-HPr(40) | 58 | 3.18E-11 | 48.40 | in-solution |
| EIN(1)-HPr(24) | 18 | 8.06E-17 | 49.07 | in-solution |
| EIN(1)-HPr(27) | 63 | 5.71E-22 | 53.15 | in-solution |
| EIN(1)-HPr(72) | 32 | 1.17E-27 | 60.88 | in-solution |
| EIN(1)-HPr(1) | 37 | 1.26E-31 | 64.70 | in-solution |

b
| Linked aa positions | Total Spec | Best E-value | Ca-Ca distance (Å)<br>(PDB code: 3EZA) | label |
| --- | --- | --- | --- | --- |
| HPr(49)-EIN(96) | 5 | 1.92E-19 | 24.33 | in-solution |
| EIN(49)-HPr(49) | 21 | 1.05E-17 | 26.48 | in-solution |
| EIN(30)-HPr(49) | 83 | 3.73E-31 | 29.90 | in-solution |
| HPr(24)-EIN(30) | 47 | 1.95E-12 | 36.78 | in-solution |
| HPr(24)-EIN(238) | 11 | 5.31E-16 | 56.10 | in-solution |
| HPr(27)-EIN(238) | 6 | 1.66E-20 | 61.06 | in-solution |

C
| Linked aa positions | Total Spec | Best E-value | Ca-Ca distance (Å)<br>(PDB code: 3EZA) | label |
| --- | --- | --- | --- | --- |
| HPr(24)-EIN(58) | 526 | 3.21E-36 | 15.38 | in-gel |
| HPr(27)-EIN(58) | 238 | 5.11E-14 | 19.62 | in-gel |
| HPr(24)-EIN(49) | 94 | 1.58E-12 | 22.32 | in-gel |
| HPr(27)-EIN(49) | 169 | 4.16E-31 | 23.42 | in-gel |
| EIN(49)-HPr(49) | 702 | 2.37E-24 | 26.48 | in-gel |
| HPr(24)-EIN(175) | 43 | 2.83E-22 | 34.05 | in-gel |
| HPr(27)-EIN(30) | 156 | 2.81E-16 | 35.30 | in-gel |
| HPr(40)-EIN(49) | 27 | 4.61E-24 | 36.39 | in-gel |
| HPr(24)-EIN(30) | 303 | 5.70E-20 | 36.78 | in-gel |
| HPr(27)-EIN(29) | 36 | 2.84E-26 | 36.85 | in-gel |
| HPr(24)-EIN(29) | 56 | 2.87E-09 | 37.81 | in-gel |
| EIN(30)-HPr(40) | 44 | 4.52E-21 | 38.01 | in-gel |
| EIN(1)-HPr(40) | 5 | 1.94E-10 | 48.40 | in-gel |
| EIN(1)-HPr(24) | 81 | 4.86E-13 | 49.07 | in-gel |
| EIN(1)-HPr(27) | 62 | 3.69E-21 | 53.15 | in-gel |

d
| Linked aa positions | Total Spec | Best E-value | Ca-Ca distance (Å)<br>(PDB code: 3EZA) | label |
| --- | --- | --- | --- | --- |
| HPr(24)-EIN(58) | 233 | 8.11E-40 | 15.38 | in-gel |
| HPr(27)-EIN(58) | 504 | 4.98E-23 | 19.62 | in-gel |
| HPr(24)-EIN(60) | 15 | 1.31E-09 | 20.21 | in-gel |
| HPr(27)-EIN(69) | 45 | 1.65E-14 | 20.35 | in-gel |
| HPr(49)-EIN(69) | 28 | 1.58E-18 | 22.20 | in-gel |
| HPr(24)-EIN(49) | 127 | 2.89E-33 | 22.32 | in-gel |
| HPr(27)-EIN(49) | 86 | 3.44E-23 | 23.42 | in-gel |
| HPr(49)-EIN(96) | 638 | 3.05E-41 | 24.33 | in-gel |
| EIN(49)-HPr(49) | 78 | 6.50E-18 | 26.48 | in-gel |
| EIN(30)-HPr(49) | 623 | 1.18E-34 | 29.90 | in-gel |
| EIN(29)-HPr(49) | 24 | 6.25E-16 | 30.92 | in-gel |
| HPr(40)-EIN(96) | 56 | 1.95E-10 | 31.79 | in-gel |
| HPr(27)-EIN(30) | 274 | 2.13E-21 | 35.30 | in-gel |
| HPr(40)-EIN(49) | 77 | 3.25E-28 | 36.39 | in-gel |
| HPr(24)-EIN(30) | 209 | 8.37E-23 | 36.78 | in-gel |
| HPr(27)-EIN(29) | 78 | 3.22E-26 | 36.85 | in-gel |
| HPr(24)-EIN(29) | 37 | 1.21E-10 | 37.81 | in-gel |
| EIN(30)-HPr(40) | 130 | 2.27E-31 | 38.01 | in-gel |
| EIN(29)-HPr(40) | 24 | 5.62E-16 | 39.11 | in-gel |
| HPr(1)-EIN(49) | 66 | 6.96E-26 | 41.09 | in-gel |
| HPr(1)-EIN(30) | 7 | 2.05E-19 | 46.92 | in-gel |
| HPr(24)-EIN(238) | 229 | 2.10E-13 | 56.10 | in-gel |
| HPr(49)-EIN(238) | 65 | 1.76E-20 | 56.45 | in-gel |
| HPr(40)-EIN(238) | 4 | 3.15E-09 | 60.32 | in-gel |
| HPr(27)-EIN(238) | 205 | 2.49E-32 | 61.06 | in-gel |

### Synthetic procedures and analytical data

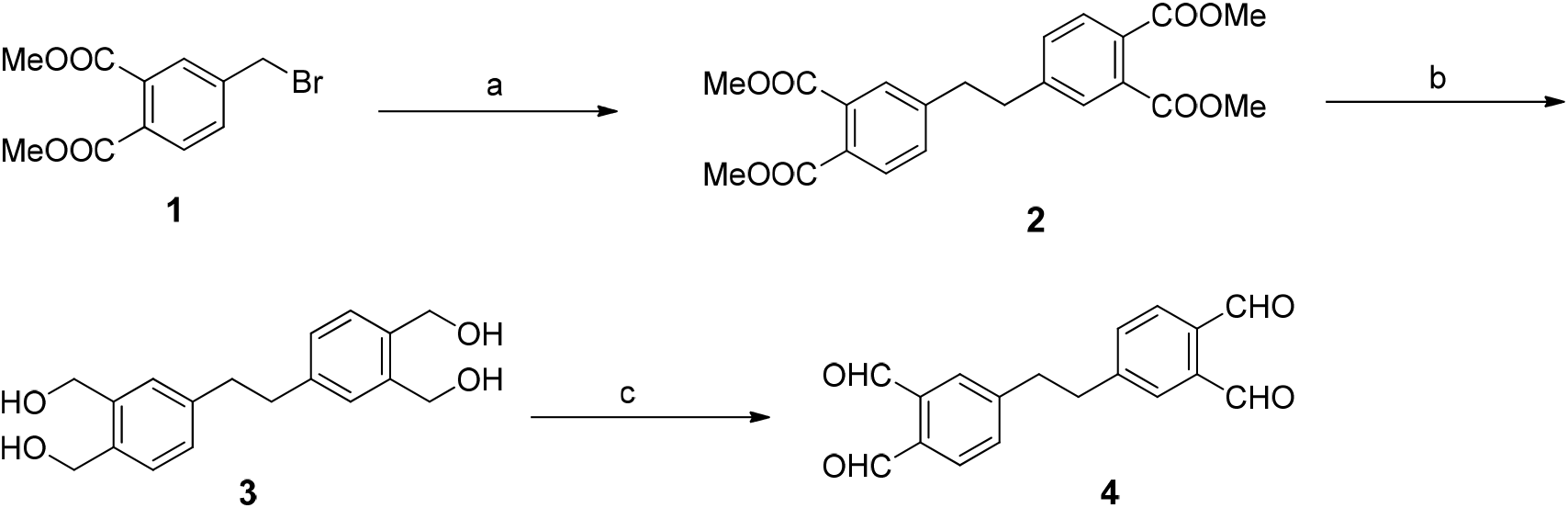

Reagents and conditions: (a) Me_2_Zn, RhCl(PPh_3_)_3_, THF, 24 h, 76%; (b) DIBAL-H, THF, 12 h; (c) Dess-Martin periodinane, DCM, 12 h, 98% for two steps.

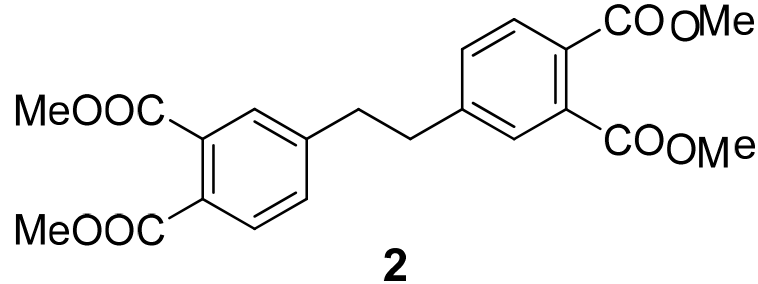

Compound **2**: To a solution of RhCl (PPh_3_)_3_ (18.5 mg, 0.02 mmol) and compound **1** (287.0 mg, 1.00 mmol) in anhydrous and thoroughly degassed THF (5.0 mL) was added Me_2_Zn (1.0 mL, 1.00 mmol, 1.0 M in hexanes) at room temperature. The resulting mixture was stirred under argon atmosphere for 24 h. After that, the reaction mixture was quenched with 2 M HCl (5.0 mL), and extracted with EtOAc (20.0 mL×3). The combined organic layers were washed with brine (10.0 mL). After dried over Na_2_SO_4_, the solution was concentrated *in vacuo* and purified by silica gel flash chromatography (petrol ether/EtOAc, 4:1 - 7:3) to afford the desired compound **2** as a white solid (315.0 mg,76%).

^1^H NMR (400 MHz, CDCl_3_) δ 7.67 (d, *J* = 8.0 Hz, 2H), 7.50 (d, *J* = 2.0 Hz, 2H), 7.27 (dd, *J* = 8.0, 2.0 Hz, 2H), 3.90 (d, *J* = 8.0 Hz, 12H), 2.99 (s, 4H);

^13^C NMR (100 MHz, CDCl_3_) δ 168.44, 167.83, 144.75, 132.82, 131.12, 129.58, 129.50, 128.85, 52.84, 52.75, 37.10;

HRMS (ESI) *m/z* calculated for C_22_H_23_O_8_ (M+H)^+^ 414.1387, found 415.1387.

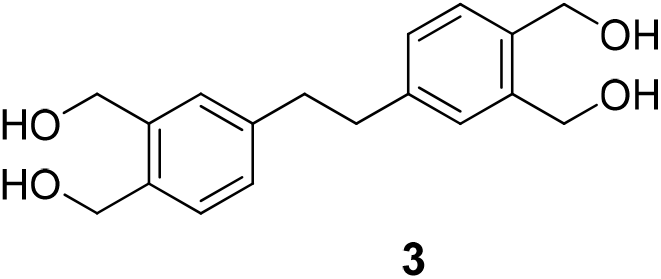

Compound **3**: To a solution of compound **2** (127.0 mg, 0.31 mmol) in anhydrous THF (9.0 mL) was added DIBAL-H (3.0 mL, 3.00 mmol, 1.0 M in hexanes) at room temperature. The resulting mixture was stirred under argon atmosphere for 12 h. After that, the reaction mixture was quenched with potassium sodium tartrate solution (10.0 mL). Most organic solvents were removed under reduced pressure and the mixture was extracted with *i*-PrOH:CHCl_3_ = 1:3 (40.0 mL×3). The combined organic layers were washed with brine (10.0 mL). After dried over Na_2_SO_4_, the solution was concentrated *in vacuo* to afford the desired compound **3** as a white solid without further purification.

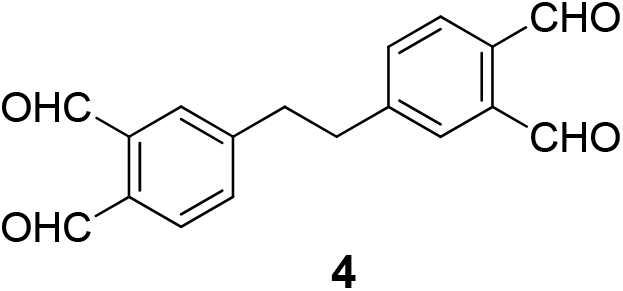

Compound **4**: To a solution of compound **3** (90.0 mg, 0.30 mmol) in anhydrous DCM (18.0 mL) was added Dess-Martin periodinane (1.27 g, 3.70 mmol) at room temperature. The resulting mixture was stirred under argon atmosphere for 12 h. After that, the reaction mixture was quenched with Sat. NaHCO_3_ (10.0 mL) and Sat. Na_2_S_2_O_3_ (10.0 mL). The mixture was stirred for 1 h and extracted with DCM (30.0 mL×3). The combined organic layers were washed with brine (10.0 mL). After dried over Na_2_SO_4_, the solution was concentrated *in vacuo* and purified by silica gel flash chromatography (petrol ether/EtOAc, 4:1 - 2:3 with 5% dichloromethane) to afford the desired compound **4** as a white solid (86.7 mg, 98% for two steps).

^1^H NMR (500 MHz, CDCl_3_) δ 10.58 (s, 2H), 10.45 (s, 2H), 7.89 (d, *J* = 8.0 Hz, 2H), 7.79 (d, *J* = 2.0 Hz, 2H), 7.52 (dd, *J* = 8.0, 2.0 Hz, 2H), 3.15 (s, 4H);

^13^C NMR (125 MHz, CDCl_3_) δ 192.26, 191.99, 147.26, 136.87, 134.94, 133.78, 132.30, 130.60, 37.03;

HRMS (ESI) m/z calculated for C_18_H_15_O_4_ (M+H)^+^ 295.0970, found 295.0965.

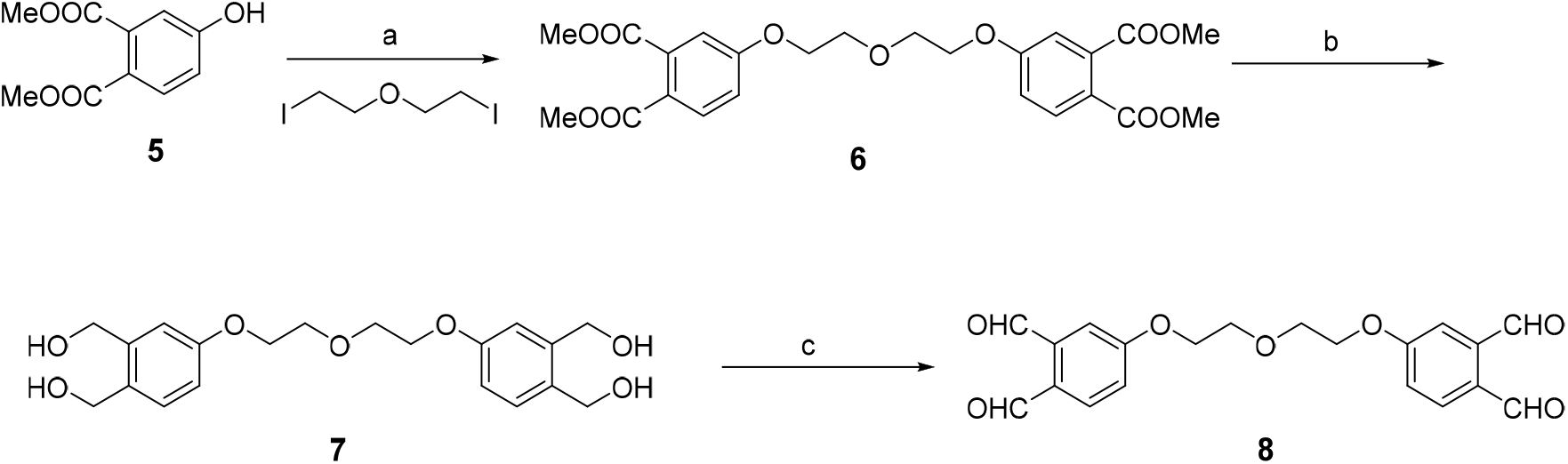

Reagents and conditions: (a) Cs_2_CO_3_, DMF, 80 °C, 12 h, 77%; (b) DIBAL-H, THF, 50 °C, 2.5 h; (c) Dess-Martin periodinane, DCM, 12 h, 93% for two steps.

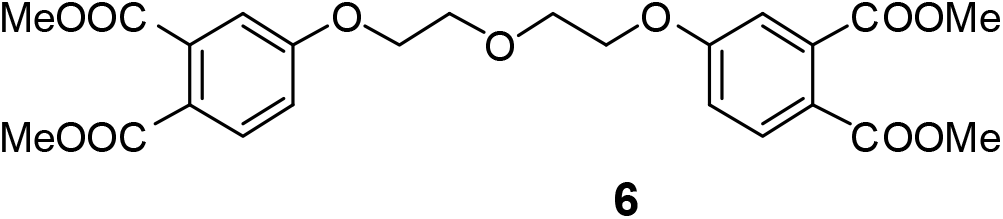

Compound **6**: To a solution of Cs_2_CO_3_ (600.0 mg, 1.84 mmol) and compound **5** (322.0 mg, 1.53 mmol) in anhydrous DMF (6.0 mL) was added 1,5-diiodo-3-oxopentane (200.0 mg, 0.61 mmol) at 80 °C. The resulting mixture was stirred under argon atmosphere for 12h. After that, the reaction mixture was filtered to remove the solid then the filtrate was concentrated *in vacuo* and purified by silica gel flash chromatography (petrol ether/EtOAc, 4:1 - 1:1) to afford the desired compound **6** as a white solid (578.2 mg, 77%).

^1^H NMR (400 MHz, CDCl_3_) δ 7.79 (d, *J* = 8.0 Hz, 2H), 7.09 (d, *J* = 2.0 Hz, 2H), 7.00 (dd, *J* = 8.0, 2.0 Hz, 2H), 4.20 (m, 4H), 3.93 (m, 4H), 3.88 (d, *J* = 16.0 Hz, 12H);

^13^C NMR (100 MHz, CDCl_3_) δ 168.86, 166.92, 161.30, 135.75, 131.70, 122.63, 116.50, 114.25, 69.84, 68.00, 52.91, 52.54;

HRMS (ESI) m/z calculated for C_24_H_27_O_11_ (M+H)^+^ 491.1553, found 491.1548.

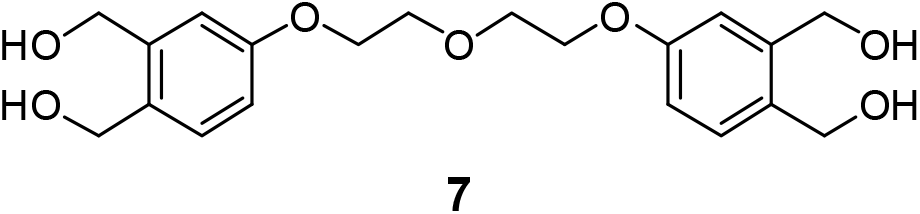

Compound **7**: To a solution of compound 6 (73.5 mg, 0.15 mmol) in anhydrous THF (3.5 mL) was added DIBAL-H (1.5 mL, 1.50 mmol, 1.0 M in hexanes) at room temperature. The resulting mixture was stirred under argon atmosphere for 2.5 h under 50 °C. After that, the reaction mixture was quenched with potassium sodium tartrate solution (5.0 mL). Most organic solvents were removed under reduced pressure and the mixture was extracted with *i*-PrOH:CHCl_3_ = 1:3 (40.0 mL×3).The combined organic layers were washed with brine (5.0 mL). After dried over Na_2_SO_4_, the solution was concentrated *in vacuo* to afford the desired compound **7** as a white solid without further purification.

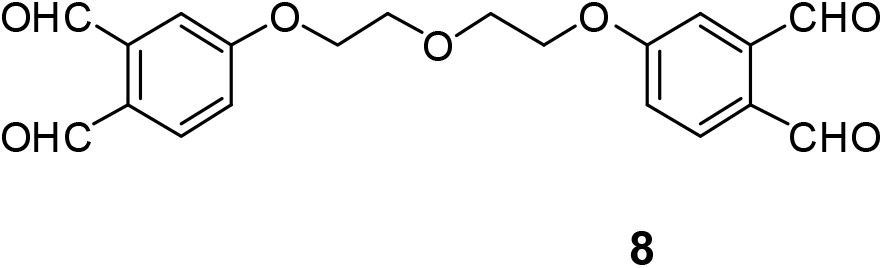

Compound **8**: To a solution of compound **7** (~56.0 mg, 0.15 mmol) in anhydrous DCM (9.0 mL) was added Dess-Martin periodinane (382.0 mg, 0.90 mmol) at room temperature. The resulting mixture was stirred under argon atmosphere for 12 h. After that, the reaction mixture was quenched with Sat. NaHCO_3_ (5.0 mL) and Sat. Na_2_S_2_O_3_ (5.0 mL). The mixture was stirred for 1 h and extracted with DCM (15.0 mL×3). The combined organic layers were washed with brine (5.0 mL). After dried over Na_2_SO_4_, the solution was concentrated *in vacuo* and purified by silica gel flash chromatography (petrol ether/EtOAc, 3:1 - 3:2) to afford the desired compound **8** as a white solid (51.7 mg, 93% for two steps).

^1^H NMR (500 MHz, CDCl_3_) δ 10.65 (s, 2H), 10.32 (s, 2H), 7.91 (d, *J* = 8.0 Hz, 2H), 7.46 (d, *J* = 2.0 Hz, 2H), 7.23 (dd, *J* = 8.0, 2.0 Hz, 2H), 4.30 (m, 4H), 3.98 (m, 4H);

^13^C NMR (125 MHz, CDCl_3_) δ 191.95, 191.07, 163.14, 138.78, 134.82, 129.90, 119.48, 115.33, 69.85, 68.30;

HRMS (ESI) *m/z* calculated for C_20_H_19_O_7_ (M+H)^+^ 371.1125, found 371.1123.

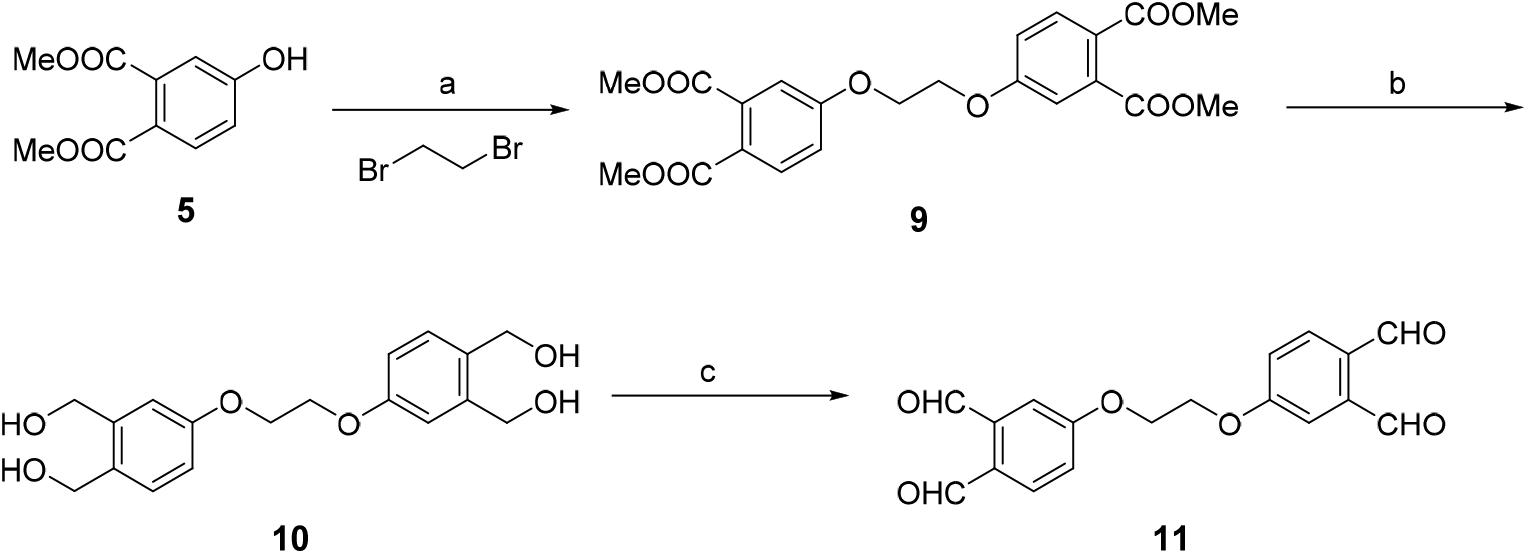

Reagents and conditions: (a) Cs_2_CO_3_, DMF, 80 °C, 12 h, 37%; (b) DIBAL-H, THF, 12 h; (c) Dess-Martin periodinane, DCM, 6 h, 80% for two steps.

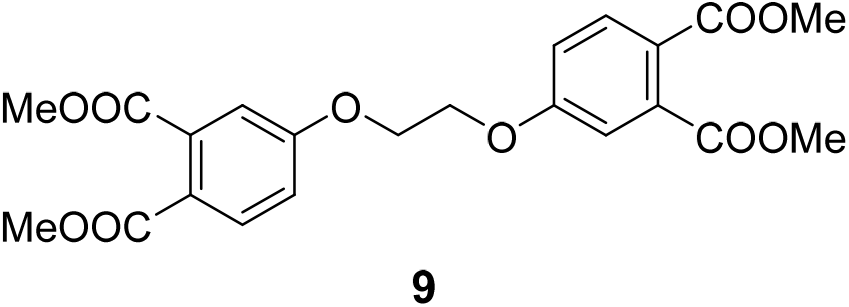

Compound **9**: To a solution of Cs_2_CO_3_ (6.2 g, 19.00 mmol) and compound **5** (2.1 g, 10.00 mmol) in anhydrous DMF (100.0 mL) was added ethylene dibromide (7.1 g, 38.00 mmol) at 80 °C. The resulting mixture was stirred under argon atmosphere for 12h. After that, the reaction mixture was filtered to remove the solid then the filtrate was concentrated *in vacuo* and purified by silica gel flash chromatography (petrol ether/EtOAc, 10:1 - 5:1) to afford the desired compound **9** as a white solid. If anyone wants to improve the yield of this step, you can decrease the ratio of dibromide (1.6g, 37%).

^1^H NMR (400 MHz, CDCl_3_) δ 7.81 (d, *J* = 8.0 Hz, 2H), 7.12 (d, *J* = 2.0 Hz, 2H), 7.03 (dd, *J* = 8.0, 2.0 Hz, 2H), 4.39 (s, 4H), 3.89 (d, *J* = 16.0 Hz, 12H);

^13^C NMR (100 MHz, CDCl_3_) δ 168.73, 166.87, 160.97, 135.77, 131.74, 123.02, 116.57, 114.20, 66.72, 52.93, 52.57;

HRMS (ESI) *m/z* calculated for C_22_H_23_O_10_ (M+H)^+^ 447.1286, found 447.1282.

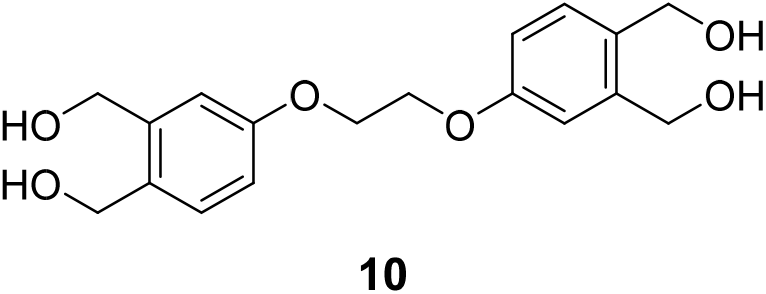

Compound **10**: To a solution of compound **9** (717.2 mg, 1.61 mmol) in anhydrous THF (40.0 mL) was added DIBAL-H (16.1 mL, 16.10 mmol, 1.0 M in hexanes) at room temperature. The resulting mixture was stirred under argon atmosphere for 12 h. After that, the reaction mixture was quenched with potassium sodium tartrate solution (25.0 mL). Most organic solvents were removed under reduced pressure and the mixture was extracted with *i*-PrOH:CHCl_3_ = 1:3 (100.0 mL×3).The combined organic layers were washed with brine (50.0 mL). After dried over Na_2_SO_4_, the solution was concentrated *in vacuo* to afford the desired compound **10** as a white solid without further purification.

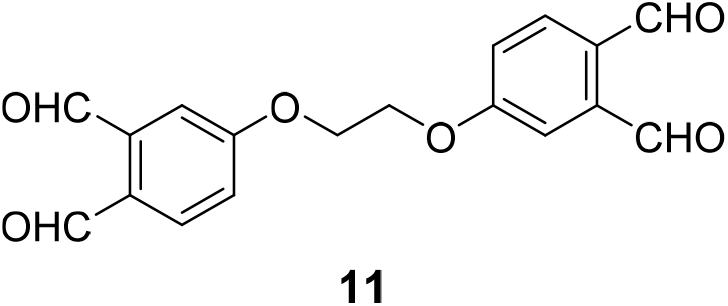

Compound **11**: To a solution of compound **10** (~150.0 mg, 0.45 mmol) in anhydrous DCM (40.0 mL) was added Dess-Martin periodinane (1.1 g, 2.69 mmol) at room temperature. The resulting mixture was stirred under argon atmosphere for 6 h. After that, the reaction mixture was quenched with Sat. NaHCO_3_ (20.0 mL) and Sat. Na_2_S_2_O_3_ (20.0 mL). The mixture was stirred for 1 h and extracted with DCM (40.0 mL×3). The combined organic layers were washed with brine (30.0 mL). After dried over Na_2_SO_4_, the solution was concentrated *in vacuo* and purified by silica gel flash chromatography (petrol ether/EtOAc, 5:1 - 1:2) to afford the desired compound **11** as a white solid (119.0 mg, 80% for two steps).

^1^H NMR (400 MHz, CDCl_3_) δ 10.69 (s, 2H), 10.34 (s, 2H), 7.96 (d, *J* = 8.0 Hz, 2H), 7.53 (d, *J* = 2.0 Hz, 2H), 7.29 (dd, *J* = 8.0, 2.0 Hz, 2H), 4.53 (s, 4H);

^13^C NMR (100 MHz, CDCl_3_) δ 191.84, 191.06, 162.71, 138.83, 134.91, 130.20, 119.70, 115.01, 66.92;

HRMS (ESI) *m/z* calculated for C_18_H_15_O_6_ (M+H)^+^ 327.0863, found 327.0855.

### 1H NMR and 13C NMR spectra

^1^H NMR of compound 2 (400 MHz, CDCl_3_)

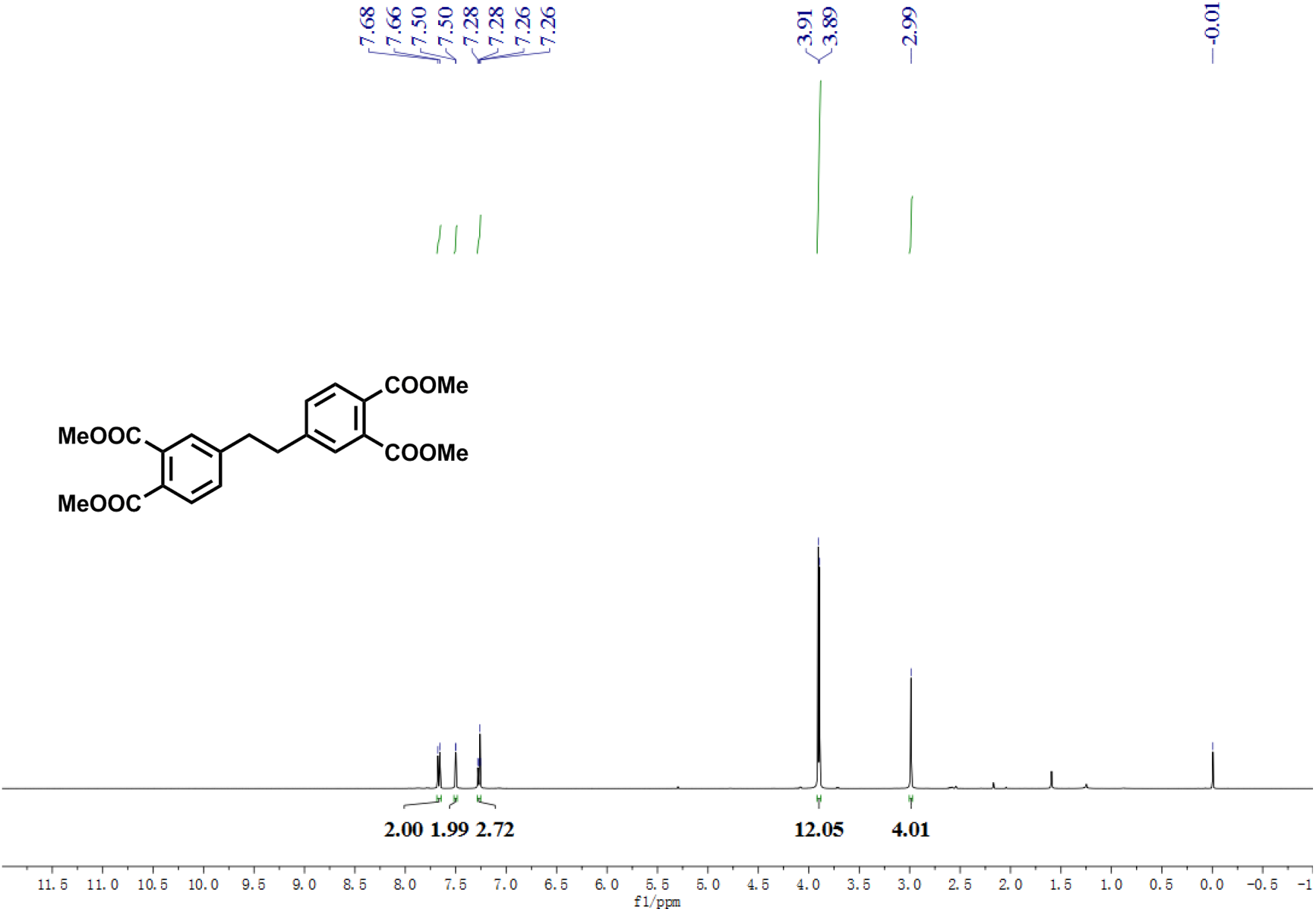

^13^C NMR of compound 2 (100 MHz, CDCl_3_)

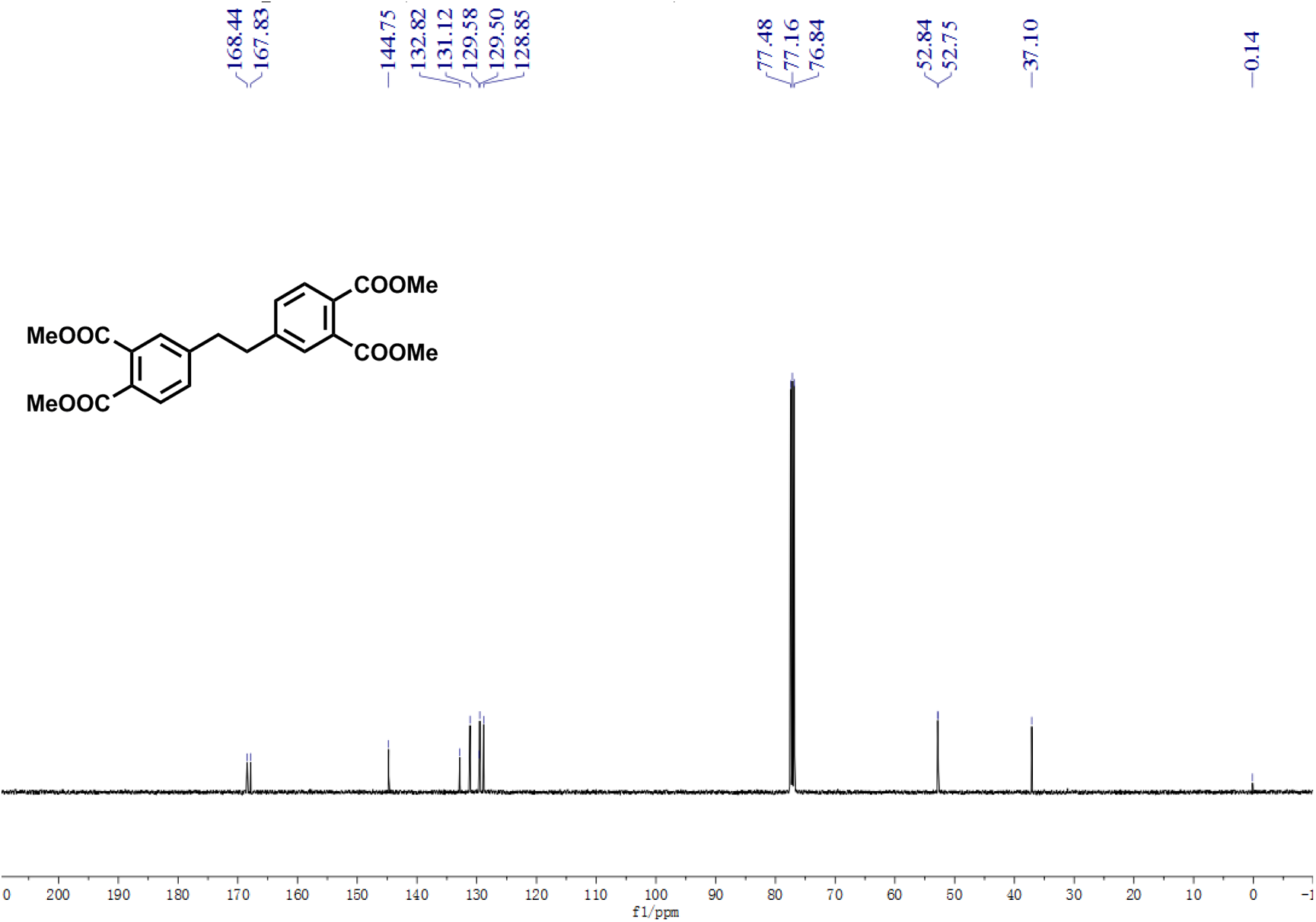

^1^H NMR of compound 4 (500 MHz, CDCl_3_)

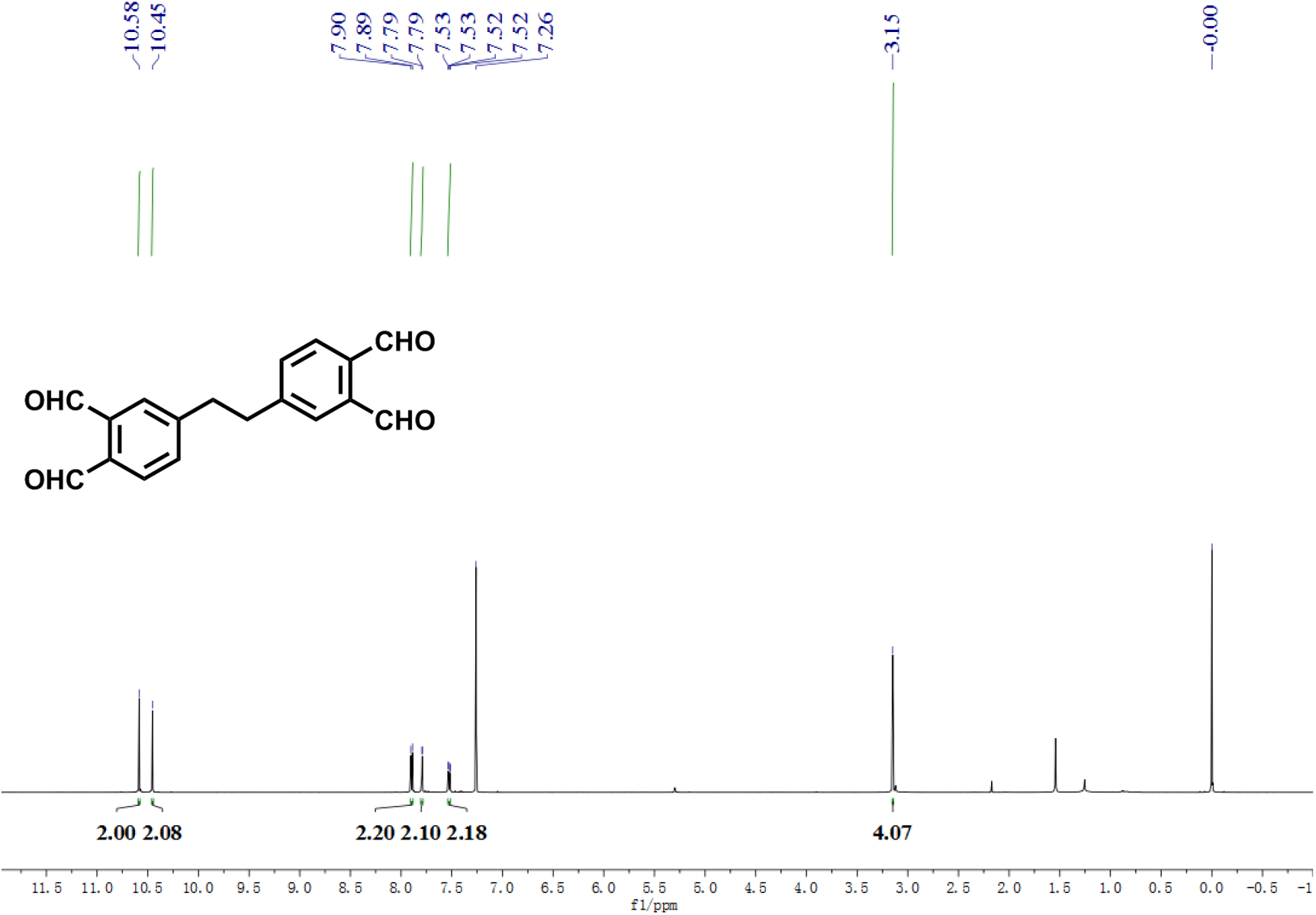

^13^C NMR of compound 4 (125 MHz, CDCl_3_)

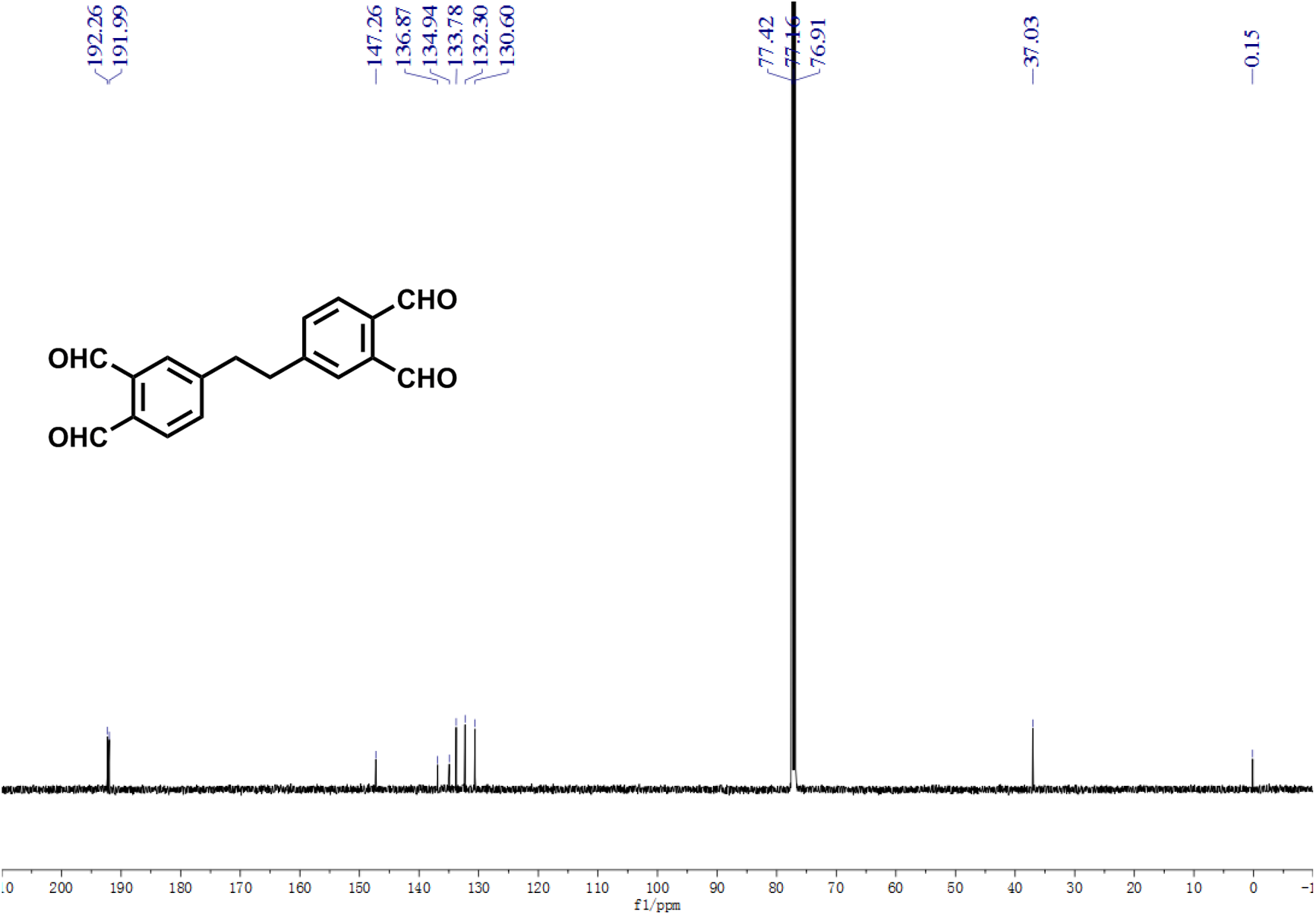

^1^H NMR of compound 6 (400 MHz, CDCl_3_)

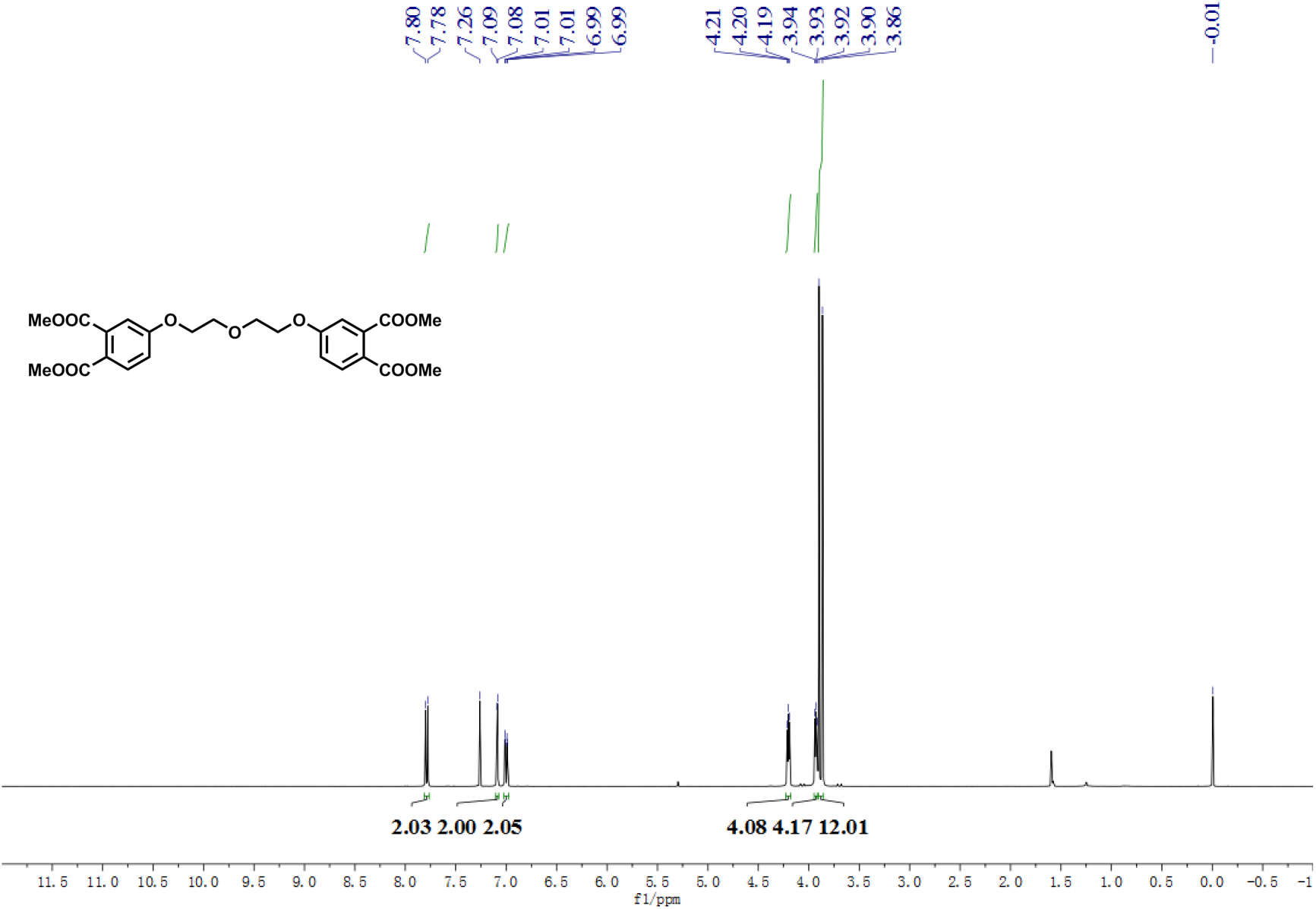

^13^C NMR of compound 6 (100 MHz, CDCl_3_)

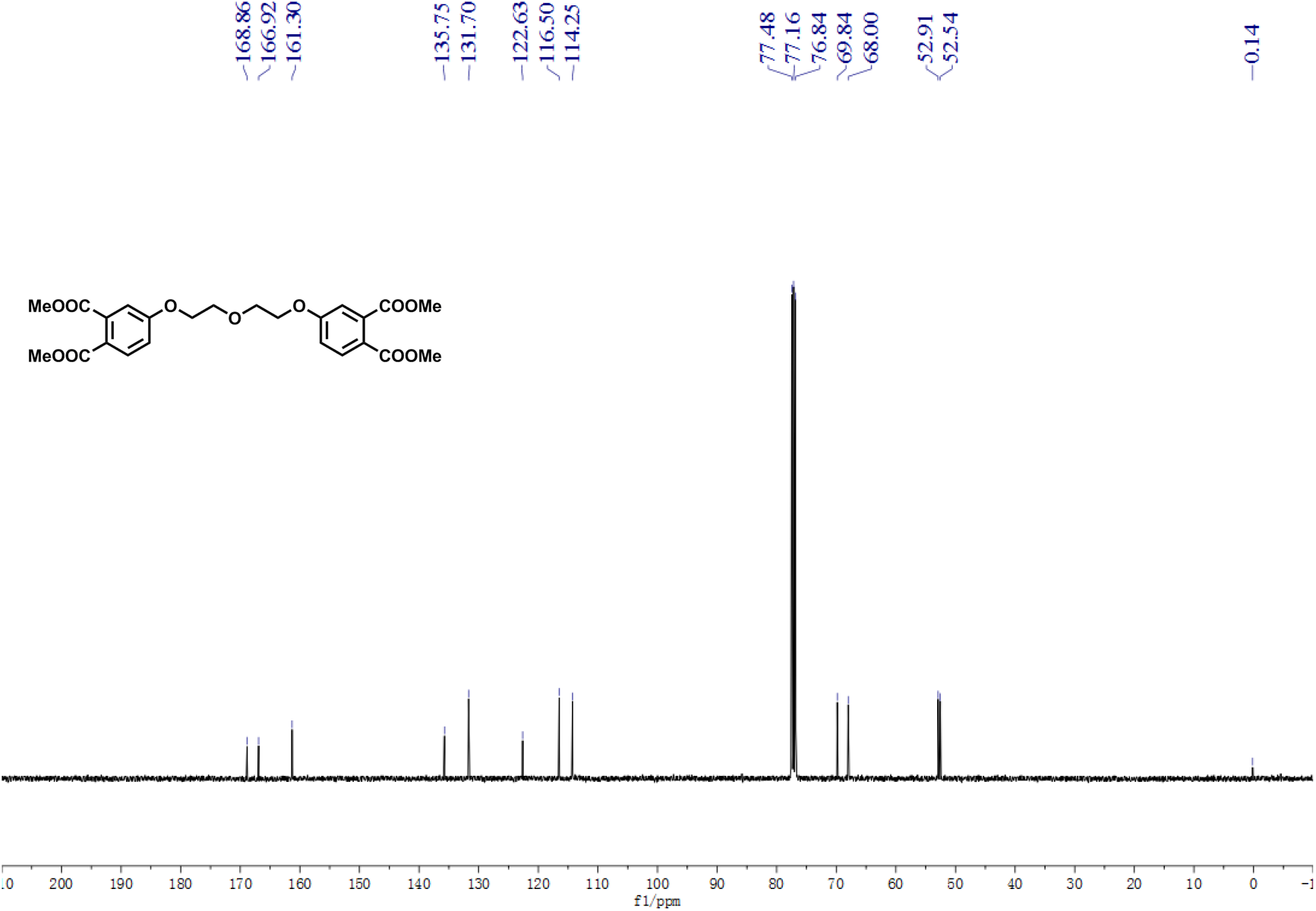

^1^H NMR of compound 8 (500 MHz, CDCl_3_)

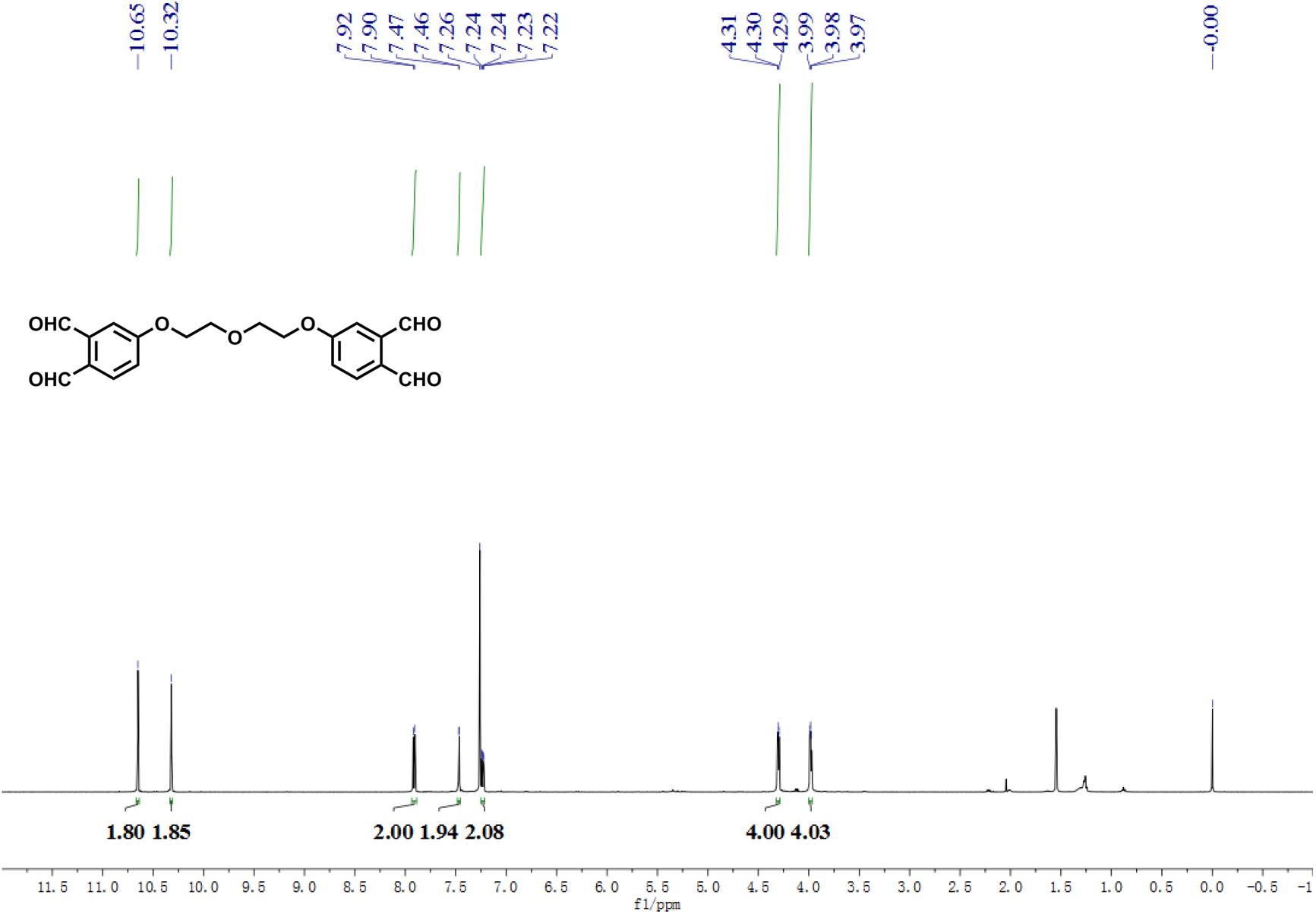

^13^C NMR of compound 8 (125 MHz, CDCl_3_)

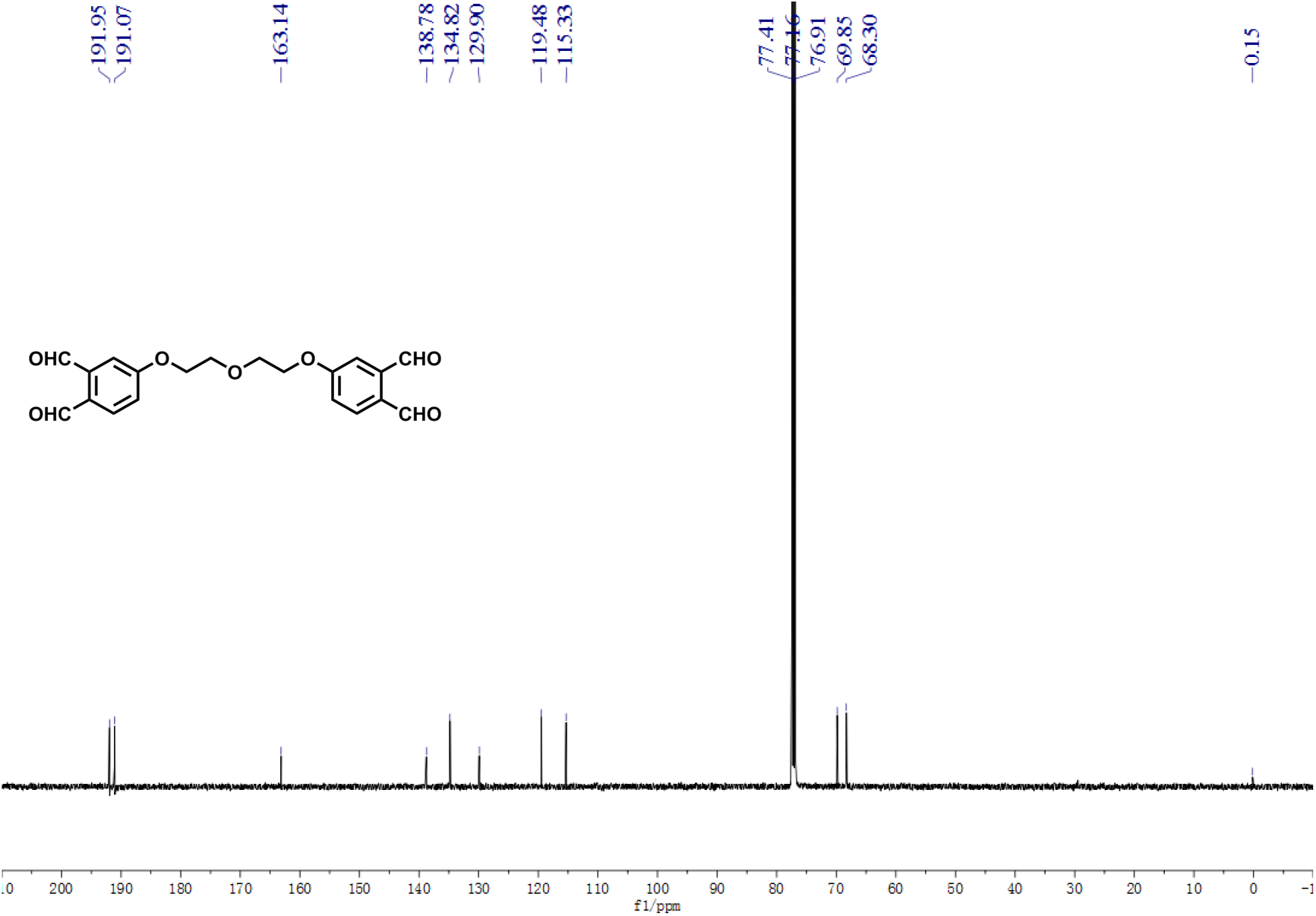

^1^H NMR of compound 9 (400 MHz, CDCl_3_)

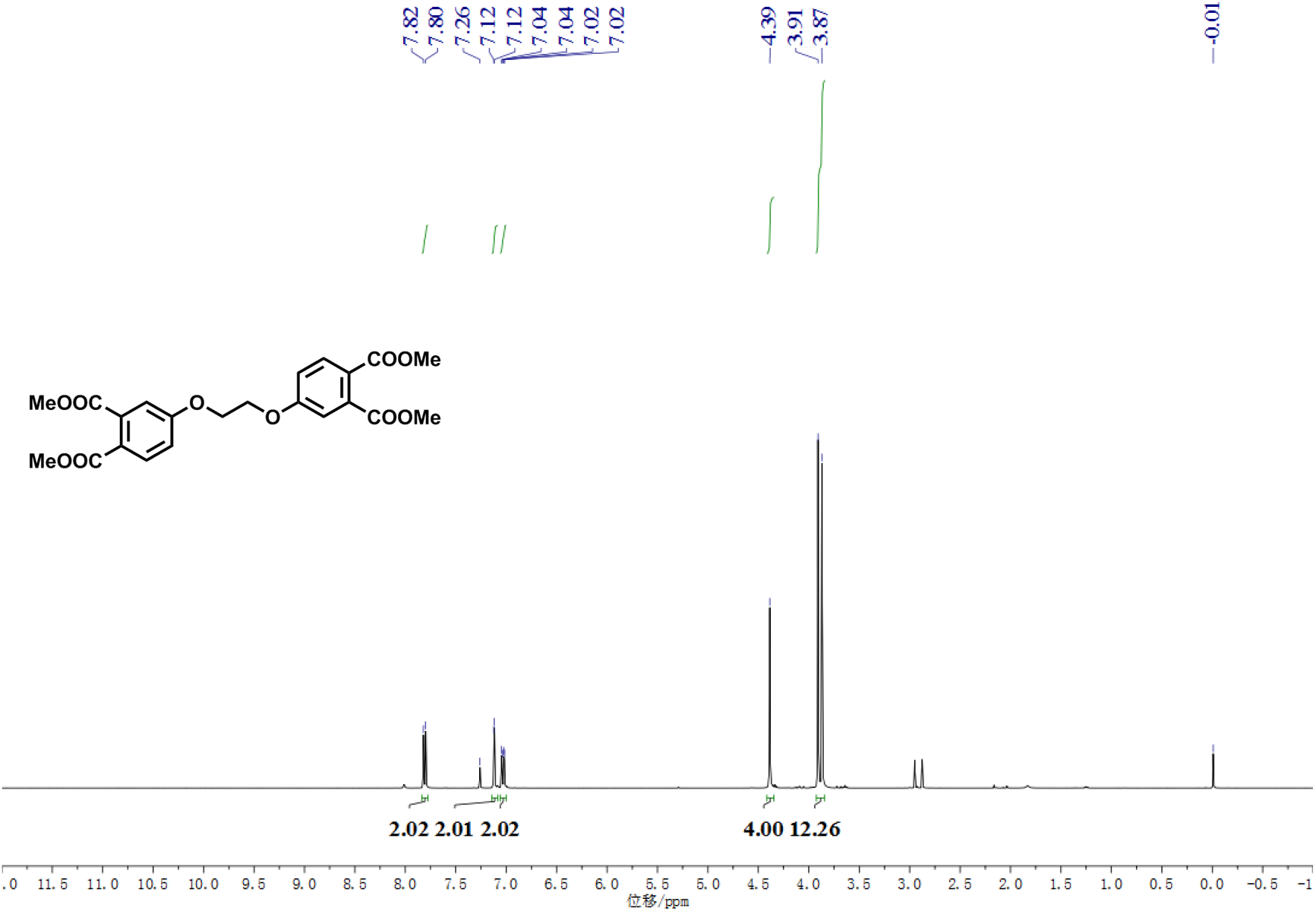

^13^C NMR of compound 9 (100 MHz, CDCl_3_)

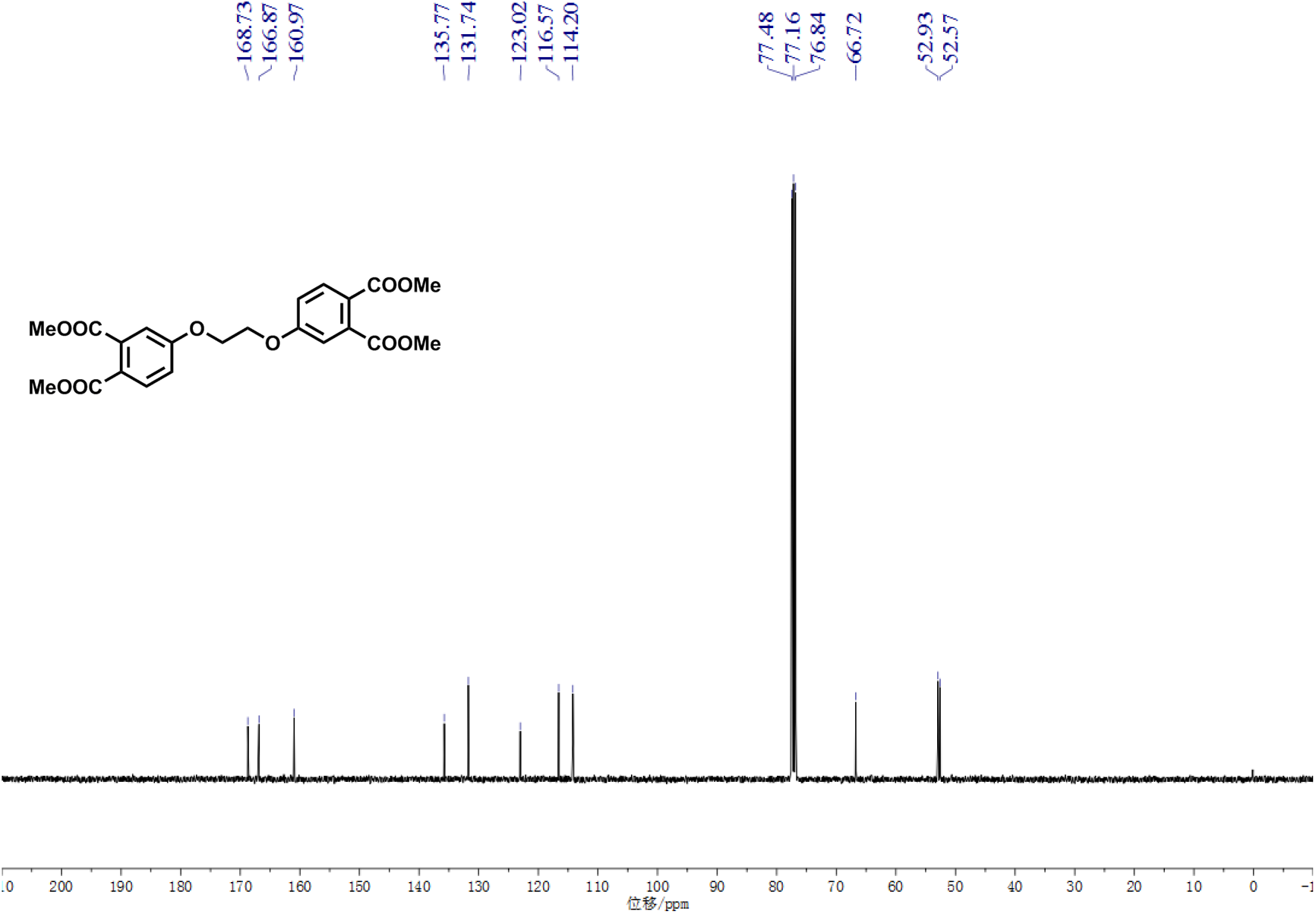

^1^H NMR of compound 11 (400 MHz, CDCl_3_)

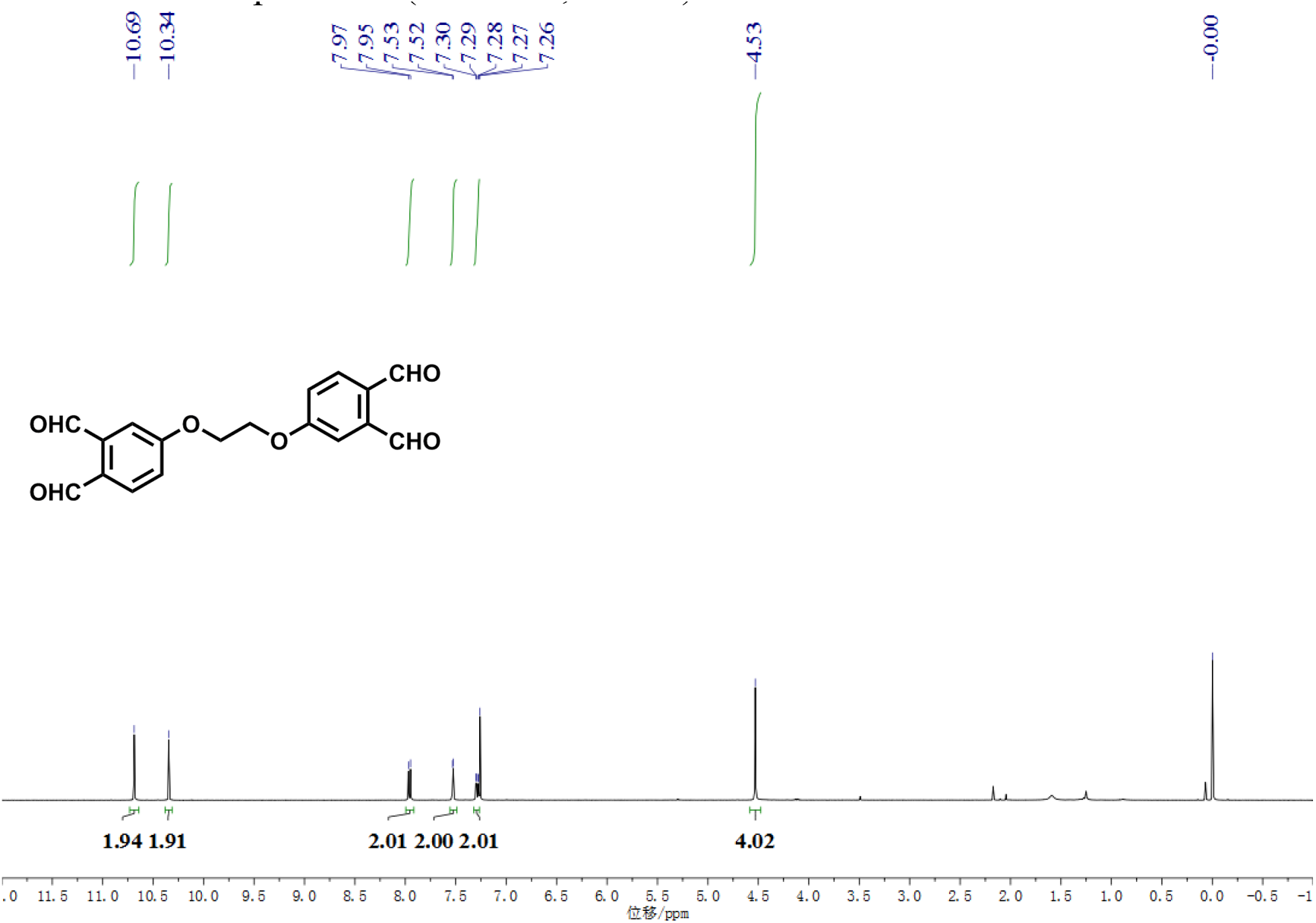

^13^C NMR of compound 11 (100 MHz, CDCl_3_)

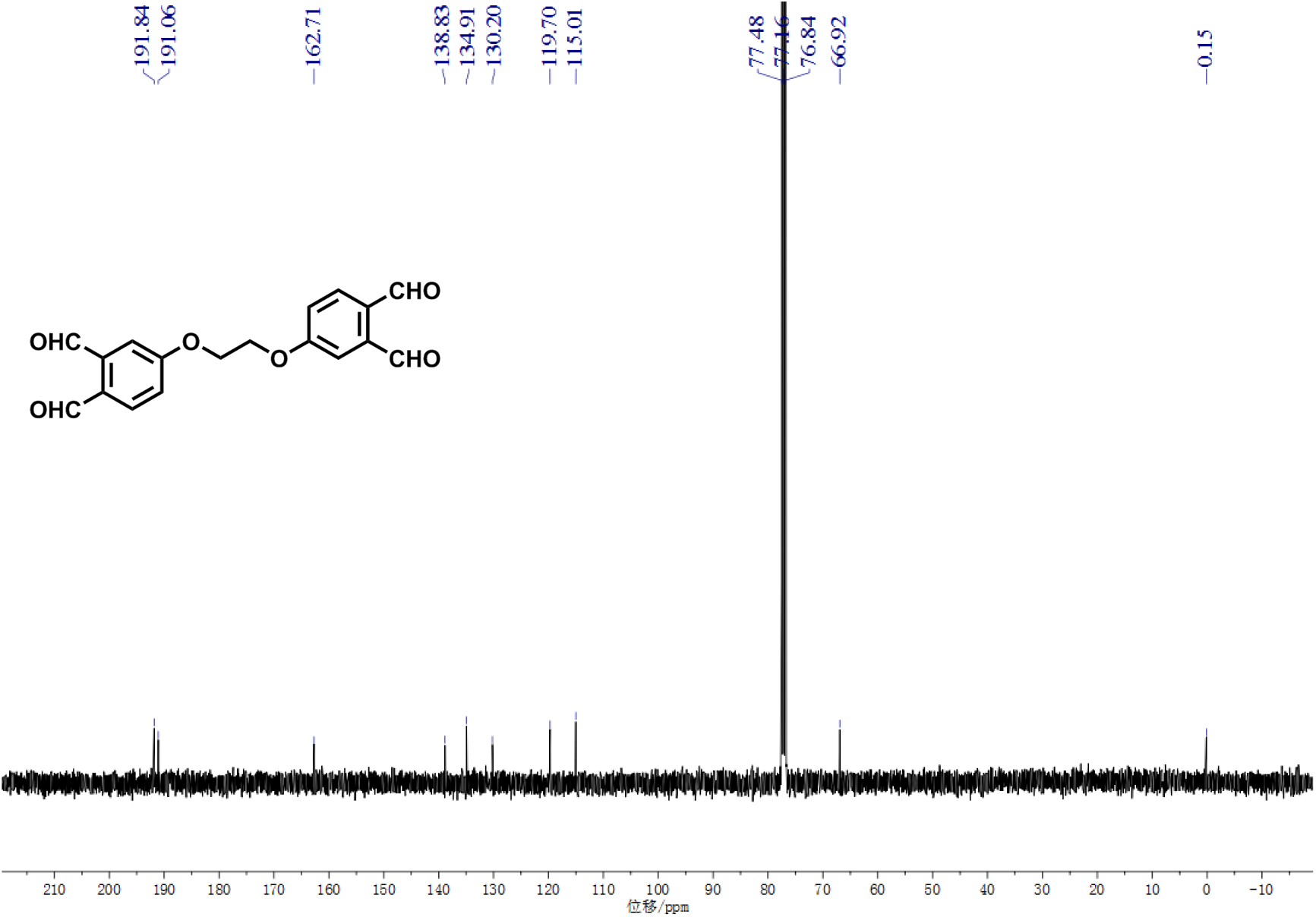

## References

1. Henzler-Wildman, K. & Kern, D. Dynamic personalities of proteins. Nature 450, 964–972 (2007).

2. Li, H., Xie, Y., Liu, C. & Liu, S. Physicochemical bases for protein folding, dynamics, and protein-ligand binding. Sci China Life Sci 57, 287–302 (2014).

3. Boehr, D.D., Nussinov, R. & Wright, P.E. The role of dynamic conformational ensembles in biomolecular recognition. Nat Chem Biol 5, 789–796 (2009).

4. Ubbink, M. The courtship of proteins: understanding the encounter complex. FEBS Lett 583, 1060–1066 (2009).

5. Englander, S.W. & Mayne, L. The nature of protein folding pathways. Proc Natl Acad Sci U S A 111, 15873–15880 (2014).

6. Wlodawer, A., Minor, W., Dauter, Z. & Jaskolski, M. Protein crystallography for aspiring crystallographers or how to avoid pitfalls and traps in macromolecular structure determination. FEBS J 280, 5705–5736 (2013).

7. Bai, X.C., McMullan, G. & Scheres, S.H. How cryo-EM is revolutionizing structural biology. Trends Biochem Sci 40, 49–57 (2015).

8. Sekhar, A. & Kay, L.E. NMR paves the way for atomic level descriptions of sparsely populated, transiently formed biomolecular conformers. Proc Natl Acad Sci U S A 110, 12867–12874 (2013).

9. Ishima, R. & Torchia, D.A. Protein dynamics from NMR. Nat Struct Biol 7, 740–743 (2000).

10. Heyduk, T. Measuring protein conformational changes by FRET/LRET. Curr Opin Biotechnol 13, 292–296 (2002).

11. Kajihara, D. et al. FRET analysis of protein conformational change through position-specific incorporation of fluorescent amino acids. Nat Methods 3, 923–929 (2006).

12. Nesmelov, Y.E. & Thomas, D.D. Protein structural dynamics revealed by site-directed spin labeling and multifrequency EPR. Biophys Rev 2, 91–99 (2010).

13. Jain, R. et al. X-ray scattering experiments with high-flux X-ray source coupled rapid mixing microchannel device and their potential for high-flux neutron scattering investigations. The European physical journal. E. Soft matter 36, 109 (2013).

14. Jain, R. & Techert, S. Time-resolved and in-situ X-ray scattering methods beyond photoactivation: Utilizing high-flux X-ray sources for the study of ubiquitous non-photoactive proteins. Protein and peptide letters 23, 242–254 (2016).

15. Jain, R. et al. Insights into open/closed conformations of the catalytically active human guanylate kinase as investigated by small-angle X-ray scattering. European biophysics journal: EBJ 45, 81–89 (2016).

16. Hamuro, Y. et al. Rapid analysis of protein structure and dynamics by hydrogen/deuterium exchange mass spectrometry. J Biomol Tech 14, 171–182 (2003).

17. Wang, L. & Chance, M.R. Structural mass spectrometry of proteins using hydroxyl radical based protein footprinting. Anal Chem 83, 7234–7241 (2011).

18. Yang, B. et al. Identification of cross-linked peptides from complex samples. Nat Methods 9, 904–906 (2012).

19. Liu, F., Rijkers, D.T., Post, H. & Heck, A.J. Proteome-wide profiling of protein assemblies by cross-linking mass spectrometry. Nat Methods 12, 1179–1184 (2015).

20. Yu, C. & Huang, L. Cross-linking mass spectrometry: an emerging technology for interactomics and structural biology. Anal Chem 90, 144–165 (2018).

21. O’Reilly, F.J. & Rappsilber, J. Cross-linking mass spectrometry: methods and applications in structural, molecular and systems biology. Nat Struct Mol Biol 25, 1000–1008 (2018).

22. Sinz, A. Cross-Linking/mass spectrometry for studying protein structures and protein-protein interactions: where are we now and where should we go from here? Angew Chem Int Ed Engl 57, 6390–6396 (2018).

23. Chavez, J.D. & Bruce, J.E. Chemical cross-linking with mass spectrometry: a tool for systems structural biology. Curr Opin Chem Biol 48, 8–18 (2019).

24. Ding, Y.H. et al. Modeling protein excited-state structures from “over-length” chemical cross-links. J Biol Chem 292, 1187–1196 (2017).

25. Liu, T. et al. Structural Insights of WHAMM’s Interaction with Microtubules by Cryo-EM. J Mol Biol 429, 1352–1363 (2017).

26. Pilch, P.F. & Czech, M.P. Interaction of cross-linking agents with the insulin effector system of isolated fat cells. Covalent linkage of 125I-insulin to a plasma membrane receptor protein of 140,000 daltons. J Biol Chem 254, 3375–3381 (1979).

27. Staros, J.V. N-hydroxysulfosuccinimide active esters: bis(N-hydroxysulfosuccinimide) esters of two dicarboxylic acids are hydrophilic, membrane-impermeant, protein cross-linkers. Biochemistry 21, 3950–3955 (1982).

28. Lauber, M.A. & Reilly, J.P. Structural analysis of a prokaryotic ribosome using a novel amidinating cross-linker and mass spectrometry. J Proteome Res 10, 3604–3616 (2011).

29. Kao, A. et al. Development of a novel cross-linking strategy for fast and accurate identification of cross-linked peptides of protein complexes. Mol Cell Proteomics 10, M110 002212 (2011).

30. Jones, A.X. et al. Improving mass spectrometry analysis of protein structures with arginine-selective chemical cross-linkers. Nat Commun 10, 3911 (2019).

31. Gutierrez, C.B. et al. Development of a Novel Sulfoxide-Containing MS-Cleavable Homobifunctional Cysteine-Reactive Cross-Linker for Studying Protein-Protein Interactions. Anal Chem 90, 7600–7607 (2018).

32. Leitner, A. et al. Chemical cross-linking/mass spectrometry targeting acidic residues in proteins and protein complexes. Proc Natl Acad Sci U S A 111, 9455–9460 (2014).

33. Zhang, X. et al. Carboxylate-selective chemical cross-Linkers for mass spectrometric analysis of protein structures. Anal Chem 90, 1195–1201 (2018).

34. Ding, Y.H. et al. Increasing the Depth of Mass-Spectrometry-Based Structural Analysis of Protein Complexes through the Use of Multiple Cross-Linkers. Anal Chem 88, 4461–4469 (2016).

35. Leitner, A. et al. Probing native protein structures by chemical cross-linking, mass spectrometry, and bioinformatics. Mol Cell Proteomics 9, 1634–1649 (2010).

36. Thiele, J. & Schneider, J. Ueber condensationsproducte des o-Phtalaldehyds. Justus Liebigs Ann Chem 369, 287–299 (1909).

37. Viets, J.W., Deen, W.M., Troy, J.L. & Brenner, B.M. Determination of serum protein concentration in nanoliter blood samples using fluorescamine or 9-phthalaldehyde. Anal Biochem 88, 513–521 (1978).

38. Aso, C., Tagami, S. & Kunitake, T. Polymerization of aromatic aldehydes. IV. Cationic copolymerization of phthalaldehyde isomers and styrene. J Polym Sci Part A-1: Polym Chem 8, 1323–1336 (1970).

39. Lerones, C., Mariscal, A., Carnero, M., Garcia-Rodriguez, A. & Fernandez-Crehuet, J. Assessing the residual antibacterial activity of clinical materials disinfected with glutaraldehyde, o-phthalaldehyde, hydrogen peroxide or 2-bromo-2-nitro-1,3-propanediol by means of a bacterial toxicity assay. Clin Microbiol Infect 10, 984–989 (2004).

40. Tung, C.L., Wong, C.T., Fung, E.Y. & Li, X. Traceless and chemoselective amine bioconjugation via phthalimidine formation in native protein modification. Org Lett 18, 2600–2603 (2016).

41. Zhang, Y., Zhang, Q., Wong, C.T.T. & Li, X. Chemoselective peptide cyclization and bicyclization directly on unprotected peptides. J Am Chem Soc 141, 12274–12279 (2019).

42. Neira, J.L. & Rico, M. Folding studies on ribonuclease A, a model protein. Fold Des 2, R1–11 (1997).

43. Yan, Y.B., Jiang, B., Zhang, R.Q. & Zhou, H.M. Two-phase unfolding pathway of ribonuclease A during denaturation induced by dithiothreitol. Protein Sci 10, 321–328 (2001).

44. Westmoreland, D.G. & Matthews, C.R. Nuclear magnetic resonance study of the thermal denaturation of ribonuclease A: implications for multistate behavior at low pH. Proc Natl Acad Sci U S A 70, 914–918 (1973).

45. Garel, J.R., Nall, B.T. & Baldwin, R.L. Guanidine-unfolded state of ribonuclease A contains both fast- and slow-refolding species. Proc Natl Acad Sci U S A 73, 1853–1857 (1976).

46. Chen, Z.L. et al. A high-speed search engine pLink 2 with systematic evaluation for proteome-scale identification of cross-linked peptides. Nat Commun 10, 3404 (2019).

47. Kotrba, P., Inui, M. & Yukawa, H. Bacterial phosphotransferase system (PTS) in carbohydrate uptake and control of carbon metabolism. J Biosci Bioeng 92, 502–517 (2001).

48. Deutscher, J., Francke, C. & Postma, P.W. How phosphotransferase system-related protein phosphorylation regulates carbohydrate metabolism in bacteria. Microbiol Mol Biol Rev 70, 939–1031 (2006).

49. Garrett, D.S., Seok, Y.J., Peterkofsky, A., Gronenborn, A.M. & Clore, G.M. Solution structure of the 40,000 Mr phosphoryl transfer complex between the N-terminal domain of enzyme I and HPr. Nat Struct Biol 6, 166–173 (1999).

50. Garrett, D.S., Seok, Y.J., Peterkofsky, A., Clore, G.M. & Gronenborn, A.M. Identification by NMR of the binding surface for the histidine-containing phosphocarrier protein HPr on the N-terminal domain of enzyme I of the Escherichia coli phosphotransferase system. Biochemistry 36, 4393–4398 (1997).

51. Cai, M. et al. Solution structure of the phosphoryl transfer complex between the signal-transducing protein IIAGlucose and the cytoplasmic domain of the glucose transporter IICBGlucose of the Escherichia coli glucose phosphotransferase system. J Biol Chem 278, 25191–25206 (2003).

52. Suh, J.Y., Tang, C. & Clore, G.M. Role of electrostatic interactions in transient encounter complexes in protein-protein association investigated by paramagnetic relaxation enhancement. J Am Chem Soc 129, 12954–12955 (2007).

53. Reizer, J. et al. Functional interactions between proteins of the phosphoenolpyruvate:sugar phosphotransferase systems of Bacillus subtilis and Escherichia coli. J Biol Chem 267, 9158–9169 (1992).

54. Gong, Z. et al. Visualizing the ensemble structures of protein complexes using chemical cross-Linking coupled with mass spectrometry. Biophys Rep 1, 127–138 (2015).

55. Tang, C., Iwahara, J. & Clore, G.M. Visualization of transient encounter complexes in protein-protein association. Nature 444, 383–386 (2006).

56. Yang, B. et al. Proximity-enhanced SuFEx chemical cross-linker for specific and multitargeting cross-linking mass spectrometry. Proc Natl Acad Sci U S A 115, 11162–11167 (2018).

57. Xing, Q. et al. Visualizing an ultra-weak protein-protein interaction in phosphorylation signaling. Angew Chem Int Ed Engl 53, 11501–11505 (2014).

58. Li, D. et al. pFind: a novel database-searching software system for automated peptide and protein identification via tandem mass spectrometry. Bioinformatics 21, 3049–3050 (2005).

59. Wang, L.H. et al. pFind 2.0: a software package for peptide and protein identification via tandem mass spectrometry. Rapid Commun Mass Spectrom 21, 2985–2991 (2007).

60. Matthew Allen Bullock, J., Schwab, J., Thalassinos, K. & Topf, M. The Importance of Non-accessible Crosslinks and Solvent Accessible Surface Distance in Modeling Proteins with Restraints From Crosslinking Mass Spectrometry. Mol Cell Proteomics 15, 2491–2500 (2016).

61. Combe, C.W., Fischer, L. & Rappsilber, J. xiNET: cross-link network maps with residue resolution. Mol Cell Proteomics 14, 1137–1147 (2015).

